# A tunable and versatile chemogenetic near infrared fluorescent reporter

**DOI:** 10.1101/2024.04.05.588310

**Authors:** Lina El Hajji, Benjamin Bunel, Octave Joliot, Chenge Li, Alison G. Tebo, Christine Rampon, Michel Volovitch, Evelyne Fischer, Nicolas Pietrancosta, Franck Perez, Xavier Morin, Sophie Vriz, Arnaud Gautier

**Affiliations:** Sorbonne Université, École Normale Supérieure, Université PSL, CNRS, Laboratoire des Biomolécules, LBM, 75005 Paris, France; Institut de Biologie de l’ENS (IBENS), École Normale Supérieure, CNRS, INSERM, Université PSL, 75005 Paris, France; Institut Curie, Université PSL, CNRS UMR144, Paris, France; PASTEUR, Department of Chemistry, École Normale Supérieure, Université PSL, Sorbonne Université, CNRS, 75005 Paris, France; Université Paris Cité, 75006 Paris, France; Neuroscience Paris Seine-Institut de Biologie Paris Seine (NPS-IBPS) INSERM, CNRS, Sorbonne Université, Paris, France; Institut Universitaire de France

## Abstract

Near-infrared (NIR) fluorescent reporters provide additional colors for highly multiplexed imaging of cells and organisms, and enable imaging with less toxic light and higher contrast and depth. Here, we present the engineering of nirFAST, a small tunable chemogenetic NIR fluorescent reporter that is brighter than top-performing NIR fluorescent proteins in cultured mammalian cells. nirFAST is a small genetically encoded protein of 14 kDa that binds and stabilizes the fluorescent state of synthetic, highly cell-permeant, fluorogenic chromophores (so-called fluorogens) that are otherwise dark when free. Engineered to emit NIR light, nirFAST can also emit far-red or red lights through change of chromophore. nirFAST allows the imaging of proteins in live cultured mammalian cells, chicken embryo tissues and zebrafish larvae. Its near infrared fluorescence provides an additional color for high spectral multiplexing. We showed that nirFAST is well-suited for stimulated emission depletion (STED) nanoscopy, allowing the efficient imaging of proteins with subdiffraction resolution in live cells. nirFAST enabled the design of a chemogenetic green-NIR fluorescent ubiquitination-based cell cycle indicator (FUCCI) for the monitoring of the different phases of the cell cycle. Finally, bisection of nirFAST allowed the design of a fluorogenic chemically induced dimerization technology with NIR fluorescence readout, enabling the control and visualization of protein proximity.

## INTRODUCTION

Far-red (FR) and near-infrared (NIR) fluorescent proteins (FPs) have opened new possibilities for imaging biological processes in living cells and organisms^1–4^. Spectrally orthogonal to visible fluorescent proteins, biosensors and optogenetic tools, FR and NIR FPs provide additional colors for highly multiplexed experiments^5,6^. In addition, autofluorescence of most tissues is low in the FR/NIR region, facilitating high contrast imaging. Finally, FR and NIR fluorescence imaging in living systems is facilitated because light above 600 nm is less toxic for cells and penetrates better into biological tissues.

FR/NIR FPs were engineered from bacterial phytochromes, protein photoreceptors that incorporate covalently the endogenous chromophore biliverdin^3^. Highly fluorogenic, biliverdin only fluoresces when embedded in FR/NIR FPs. The apparent brightness of FR/NIR FPs in cells depends however on the availability of biliverdin and its efficacy of incorporation. Efficient incorporation of biliverdin remains a key challenge in the field as biliverdin concentration can vary in cells, and as the kinetics of incorporation is rather slow. A recent comprehensive quantitative study identified emiRFP670, miRFP680, miRFP713 and miRFP720 as the top-performing monomeric FR/NIR FPs in cultured mammalian cells^7^. NIR FP have largely benefitted from protein engineering efforts, aiming at reducing their size (usually around 35 kDa) to reduce the risk of dysfunctional fusions^4,8^, and at enhancing their cellular brightness through improvement of the efficacy of incorporation of biliverdin.

Here, we used a chemogenetic approach to engineer a small tunable NIR fluorescent reporter with quasi-instantaneous fluorescence that can be used as an alternative to biliverdin-based FR/NIR FPs. Dubbed nirFAST (near infrared fluorescence-activating and absorption-shifting tag), this chemogenetic reporter is composed of a small protein of 14 kDa that efficiently and rapidly assembles with the synthetic fluorogenic chromophore 4-hydroxy-3,5-dimethoxyphenylallylidene rhodanine (HPAR-3,5DOM) to form a fluorescent semi-synthetic assembly with emission maximum at 715 nm when excited with red light (**Fig. 1a-e**). When free at physiological pH, HPAR-3,5DOM is protonated and absorbs violet-blue light, and is almost non-fluorescent because of ultra-fast non-radiative decay (**Fig. 1d**). Directed evolution and rational design allowed the engineering of a protein cavity that stabilizes the chromophore in its red-light absorbing deprotonated state, and enables it to adopt an emissive conformation, leading to NIR fluorescence upon red light excitation. In this system, deprotonation increases the electron donation of the phenol moiety leading to red-shifted absorption and emission, while immobilization of the chromophore in a planar conformation slows down non-radiative de-excitation, leading to high fluorescence activation. This unique fluorogenic mechanism provides very high contrast even in the presence of an excess of chromophore, allowing wash-free imaging in live cells or organisms. nirFAST can also form a fluorescent assembly with 4-hydroxy-3-methoxyphenylallidene rhodanine (HPAR-3OM) that emits bright fluorescence at 680 nm. In this study, we show that nirFAST (with HPAR-3OM or HPAR-3,5DOM) is brighter than the top-performing emiRFP670 and miRFP713 in cultured mammalian cells when excited at 633 nm, and can be an efficient NIR fluorescent reporter for multiplexed imaging of proteins in mammalian cells, chicken embryo tissues and in zebrafish larva. Its excellent photophysical properties make it a suitable reporter for advanced imaging techniques such as stimulated emission depletion (STED) super-resolution microscopy. We also demonstrate the potential of nirFAST for the design of cellular sensors. The combination of nirFAST with the chemogenetic reporter pFAST^9^ allowed the generation of a green-NIR fluorescent ubiquitination-based cell cycle indicator (FUCCI) for visualizing the progression of the cell cycle in living cells. Finally, we also present the use of nirFAST for the design of a fluorogenic chemically induced dimerization technology enabling to control and visualize protein proximity in live cells.

**Fig. 1.**
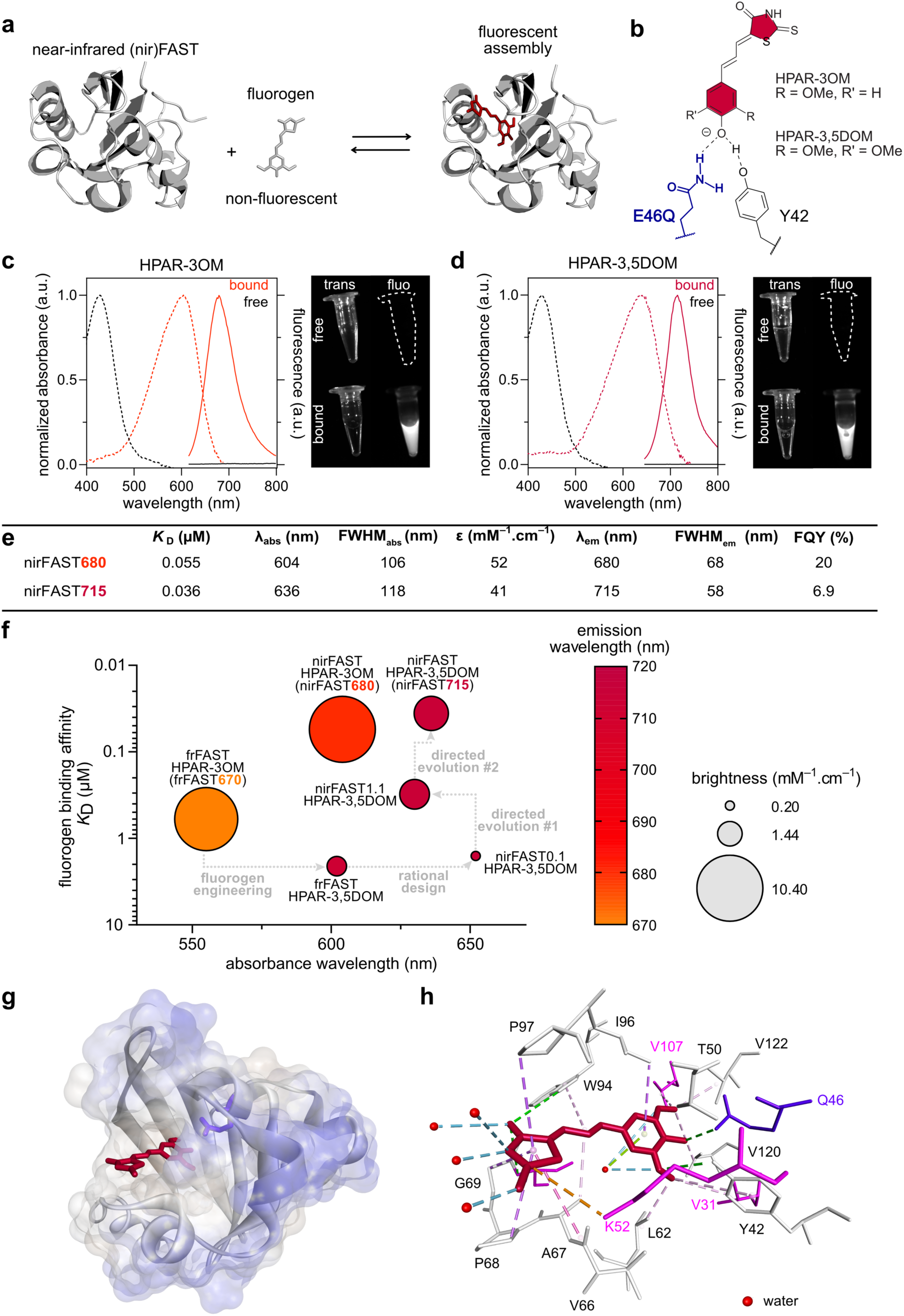
nirFAST, a versatile and tunable near-infrared fluorescent chemogenetic reporter. **(a)** Principle of nirFAST. (**b**) Structure of the anionic state of HPAR-3OM and HPAR-3,5DOM in interaction with residues Y42 and Q46. (**c**) Absorption (dashed line, left axis) and emission (solid line, right axis) spectra of HPAR-3OM free (black) or bound (orange) to nirFAST. Spectra were recorded in pH 7.4 PBS at 25 °C. Image on the right shows the fluorescence of solutions of HPAR-3OM free or bound to nirFAST under illumination with far-red light. (**d**) Absorption (dashed line) and emission (solid line) spectra of HPAR-3,5DOM free (black) or bound (dark red) to nirFAST. Spectra were recorded in pH 7.4 PBS at 25 °C. Image on the right shows the fluorescence of solutions of HPAR-3,5DOM free or bound to nirFAST under illumination with far-red light. (**e**) Properties of nirFAST with HPAR-3OM (nirFAST680) and with HPAR-3,5DOM (nirFAST715) in PBS pH 7.4. Abbreviations are as follows: *K*_D_ thermodynamic dissociation constant, λ_abs_ wavelength of maximal absorption, FWHM_abs_ full width at half-maximal absorption, ε molar absorptivity at λ_abs_, λ_em_ wavelength of maximal emission, FWHM_em_ full width at half-maximal emission, FQY fluorescence quantum yield. (**f**) Engineering of nirFAST from frFAST670: the thermodynamic dissociation constant *K*_D_, absorption wavelength, emission wavelength and brightness of the main engineering intermediates are indicated. (**g,h**) Structural model of nirFAST715 generated by homology modeling and molecular dynamics using the crystal structure of the *Halorhodospira halophila* Photoactive Yellow Protein (PYP) (PDB: 6P4I). HPAR-3,5DOM is in red, while Gln 46 is in blue. (**h**) Interactions networks involved in HPAR-3,5DOM binding and recognition within nirFAST. Residues introduced during the engineering process are shown in magenta. See also **Supplementary Figure 6** for additional details.

## RESULTS

### Pushing the spectral properties to the near-infrared

nirFAST was engineered from the chemogenetic fluorescent reporter far-red (fr)FAST^10^ (**Fig. 1f**), a reporter of the fluorescence-activating and absorption-shifting tag (FAST) family (see **Supplementary Figure 1** for a summarizing family tree)^11^. frFAST binds and stabilizes the anionic fluorescent state of HPAR-3OM. The elongated π-electron conjugation in HPAR-3OM gives a fluorescent assembly, hereafter named frFAST670, with a maximal absorption at α_abs_ = 555 nm and a maximal emission at α_em_ = 670 nm (**Fig. 1f**). Despite its FR fluorescence emission, the use of frFAST670 as FR fluorescent reporter is limited by its low absorption in the red region. To obtain a reporter emitting NIR fluorescence (> 700 nm) with high efficiency upon excitation with the 633 or 640 nm lasers classically used in fluorescence microscopy and cytometry, we introduced changes in both the fluorogen and the protein (**Fig. 1f**). **Supplementary Text 1** details the full engineering process. In brief, the use of HPAR-3,5DOM, that bears a second methoxy group in ortho position of the phenol ring, and the mutation of Glu46, which normally stabilizes the phenolate state of the chromophore in frFAST, into a glutamine led to a new fluorescent assembly, named nirFAST0.1:HPAR-3,5DOM, with a 95 nm red-shifted absorbance of the anionic phenolate state, and a 45 nm red-shifted emission with respect to frFAST. The assembly nirFAST0.1:HPAR-3,5DOM displayed an absorbance peak at α_abs_ = 650 nm and an emission maximum at α_em_ = 715 nm (**Fig. 1f**, **Supplementary Figure 2a** and **Supplementary Table 1**), which were ideal for the engineering of a NIR fluorescent reporter excitable with red light. The presence of two methoxy group in ortho position of the phenol in HPAR-3,5DOM (rather than one in HPAR-3OM) shifts absorption and emission to the red by 50 and 40 nm respectively. The introduction of a glutamine in position 46 rather than a glutamic acid additionally shifts the absorption band of the phenolate to the red by 50 nm, presumably because the reduced hydrogen-bond donating ability of glutamine versus glutamic acid increases the electron-donating ability of the anionic phenolate (see **Supplementary Text 1** for detailed explanation). However, the introduced modifications resulted in a significant decrease in brightness. Indeed, in nirFAST0.1:HPAR-3,5DOM, the fluorogen is mainly protonated as evidenced by the presence of a strong absorption band at 430 nm, characteristic of the protonated phenol state, and a much weaker band at 650 nm, characteristic of the deprotonated phenolate (**Supplementary Figure 2a**). Since only the deprotonated chromophore absorbs red light, nirFAST0.1:HPAR-3,5DOM displays a low brightness at 650 nm. In addition to decreasing brightness, the modifications of the fluorogen and the mutation E46Q also resulted in a decrease in fluorogen binding affinity (**Supplementary Figure 2b**).

### Improvement by directed evolution

To increase molecular brightness and fluorogen binding affinity, we used yeast surface display^12^ to evolve a protein variant able to efficiently bind and stabilize only the deprotonated state of HPAR-3,5DOM (see also **Supplementary Text 1**). We created a combinatorial library of variants, which we expressed at the surface of yeast cells. The yeast-displayed library was screened by fluorescence-activated cell sorting (FACS) using a 640 nm excitation laser to sort variants forming bright NIR fluorescent assemblies with HPAR-3,5DOM. After five rounds of enrichment, we isolated and characterized clones with improved brightness. The best clone was further refined by site-directed mutagenesis (**Supplementary Figures 3 and 4, Supplementary Tables 1** and **2**, **Supplementary Text 1**). The improved protein, called nirFAST1.1, efficiently binds HPAR-3,5DOM (*K*_D_ = 0.31 µM) and forms an assembly 10-fold brighter than nirFAST0.1:HPAR-3,5DOM (**Fig. 1f**, **Supplementary Figure 2a** and **Supplementary Figure 2b**).

HPAR-3,5DOM was however still partially protonated when embedded in nirFAST1.1 (**Supplementary Figure 2a**), therefore we performed a second round of directed protein evolution to maximize deprotonation of the chromophore and thus enhance molecular brightness (see also **Supplementary Text 1**). This second round of engineering, further refined by site-directed mutagenesis and an additional screening step in mammalian cells (**Supplementary Figure 5**), allowed us to identify nirFAST (with the mutations I31V, D36N, Q41K, N43S, E46Q, D48G, R52K, T70S, K78I, I107V relative to frFAST) (**Supplementary Figures 3 and 4** and **Supplementary Table 3**), a protein able to efficiently bind and fully stabilize the deprotonated state of HPAR-3,5DOM. nirFAST shows a 10-fold higher fluorogen binding affinity (*K*_D_ = 36 nM) relative to nirFAST1.1, ideal for efficient labeling in cells (**Supplementary Figure 2b**). With absorption/emission peaks at α_abs_ = 636 nm and α_em_ = 715 nm, a fluorescence quantum yield FQY = 6.9 % and a maximal molar absorption coefficient χ_636 nm_ = 41,000 M^−1^.cm^−1^, nirFAST:HPAR-3,5DOM shows great molecular brightness in the NIR window while displaying an optimal absorption maximum for excitation with common 633 and 640 nm red lasers (**Fig. 1d-f** and **Supplementary Figure 2**). Noteworthily, the absorption spectrum of nirFAST:HPAR-3,5DOM was narrower than those of other variants from the engineering process, due to the introduction of the R52K mutation. Overall, nirFAST binds HPAR-3,5DOM 50-fold tighter than nirFAST0.1, and this optimized complex is over 10-fold brighter when excited at 633 nm (**Fig. 1f**). In addition, free HPAR-3,5DOM is completely non-fluorescent when exciting with red light making it an ideal fluorogen for labeling applications in live systems. Hereafter, nirFAST assembly with HPAR-3,5DOM is named nirFAST715 for simplicity (the suffix 715 indicates the wavelength of maximal emission).

Interestingly, nirFAST also binds the far-red fluorogen HPAR-3OM with a binding affinity tenfold higher than frFAST (*K*_D_ = 55 nM versus 0.6 µM) (**Supplementary Figure 2b**). Compared to frFAST670, nirFAST:HPAR-3OM (hereafter called nirFAST680) displays similar fluorescence quantum yield (FQY = 20 %), but is characterized by a 50 nm red-shifted absorption (α_abs_ = 604 nm), a 10 nm red-shifted emission (α_em_ = 680 nm) and a narrower absorption spectrum (**Fig. 1c,e**, **Supplementary Figure 2a** and **Supplementary Tables 2** and **4**), making it overall a better FR fluorescent chemogenetic reporter.

### Structural model of nirFAST

Homology modelling using the crystal structure of the evolutionary related photoactive yellow protein (PYP) from *Halorhodospira halophila* (see family tree on **Supplementary Figure 1**) coupled to molecular dynamics allowed the generation of an atomic-resolution model of nirFAST715. Within the assembly with minimal energy, the bound HPAR-3,5DOM adopts a quasi-planar conformation in agreement with enhanced fluorescence quantum yield (**Fig. 1g,h** and **Supplementary> Figure 6a**). HPAR-3,5DOM is stably bound in its deprotonated phenolate state with the phenolate establishing hydrogen-bonds with Gln46 (from the rationally introduced E46Q mutation) and Tyr42 (conserved in PYP and all FAST variants) (**Fig. 1h**), in agreement with the red shifted absorption and emission wavelengths of nirFAST715. The presence of a structural water molecule within the binding cavity stabilizes also the chromophore in its deprotonated state (**Fig. 1h** and **Supplementary Figure 6b**), very likely by influencing the electrostatic environment of the phenolate and thus the local p*K*_A_ of the phenol moiety. The Ser70 (from the mutation T70S) establishes hydrogen-bonds with this structural water molecule (**Supplementary Figure 6b**), stabilizing the overall assembly. All the newly introduced residues are within the chromophore binding pocket or nearby (**Supplementary Figure 6a**), and participate in the shaping of the binding pocket. Noteworthily, in addition to Gln46 (from E46Q mutation) that directly interacts with the phenolate of HPAR-3,5DOM, the mutations I31V and I107V widen the bottom of the binding pocket and create a hydrophobic clamp that interacts with the methyl of the methoxy groups of HPAR-3,5DOM (**Fig. 1h** and **Supplementary Figure 6c**). This allows a deeper positioning of the phenolate into the binding cavity, favouring the stabilization of the phenolate by Gln46. The Lys52 (from R52K mutation) also participates to the binding of the chromophore by directly interacting with the rhodanine head of HPAR-3,5DOM (**Fig. 1h**). In addition, the mutation R52K enables to reduce the size of the binding pocket by allowing the residues 53 to 58 to adopt an α-helix conformation (**Supplementary Figure 6d**).

### Characterization with prototypical FAST fluorogens

Systematic testing of nirFAST with hydroxybenzylidene rhodanine (HBR) chromophores commonly used with prototypical FAST^13,14^, the parent protein of frFAST and nirFAST, showed that nirFAST was able to bind the fluorogens HMBR, HBR-2,5DM, HBR-3,5DM and HBR-3,5DOM, giving respectively, green, yellow, orange and red fluorescence, with moderate or good affinities (**Supplementary Table 5**). Two opposite behaviors were however observed: nirFAST failed to activate efficiently the fluorescence of HMBR and HBR-2,5DM, while it formed bright assemblies with HBR-3,5DM (FQY = 37 %) and HBR-3,5DOM (FQY = 24 %). The ability of nirFAST to form a tight and bright assembly with HBR-3,5DOM, hereafter called nirFAST600, opened great prospect for tunable imaging in the red-NIR region of the spectrum as it is possible to generate red, FR and NIR fluorescence with a single protein tag (nirFAST) through a simple change of fluorogen (*vide infra*). On the other hand, the inability of nirFAST to activate the green fluorescence of HMBR opened interesting prospects for two-color imaging, leveraging the spectral orthogonality of nirFAST715 and pFAST:HMBR (hereafter called pFAST540) to simultaneously visualize two distinct biological events (*vide infra*).

### Imaging of nirFAST in cultured mammalian cells

Next, we showed that nirFAST could be an efficient NIR fluorescent reporter in cultured mammalian cells. nirFAST expressed in live mammalian cells and labeled with HPAR-3OM or HPAR-3,5DOM displayed a homogenous fluorescence distribution, in agreement with high intracellular stability (**Fig. 2a**). Fusions of nirFAST to the nuclear histone H2B, to the microtubule-associated protein MAP4, to cytochrome b5 (Cb5) (for anchoring at the membrane of the endoplasmic reticulum) and to the mitochondrial targeting sequence (mito) from the subunit VIII of human cytochrome C oxidase (for expression in the matrix of mitochondria) were correctly localized when expressed in live HeLa cells (**Fig. 2a**), showing that nirFAST does not perturb cellular localization. Quantitative fluorescence measurements revealed that labeling of nirFAST after cell fixation was as efficient and as contrasted as in live cells (**Fig. 2b,c**). Fast timelapse confocal microscopy revealed that labeling with HPAR-3OM and HPAR-3,5DOM was complete within twenty seconds of fluorogen addition (**Supplementary Figure 7** and **Supplementary Movie 1**). Given the typical timescale for small molecule cell uptake and diffusion, these rapid kinetics suggest that labeling is primarily limited by the rate at which the fluorogens can be taken up and diffused into cells. Combined with the observation that HPAR-3OM and HPAR-3,5DOM are non-toxic for mammalian cells at the dose used for imaging (**Supplementary Figure 8**), this set of experiments showed that nirFAST was an excellent protein tag for the selective and efficient labeling of proteins in cultured mammalian cells.

**Fig. 2.**
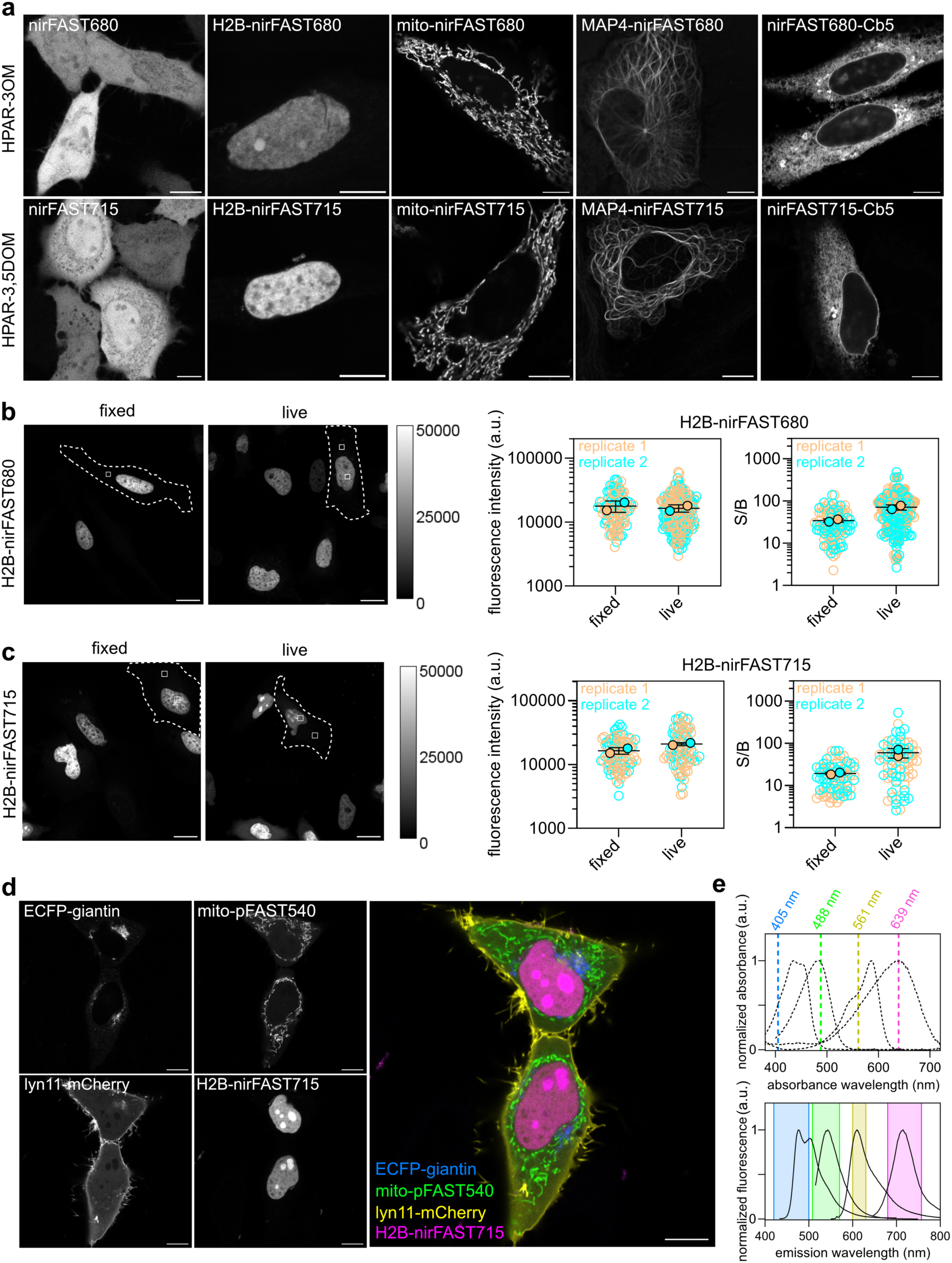
Selective imaging of nirFAST in various localizations in mammalian cells. (**a**) Confocal micrographs of live HeLa cells expressing nirFAST-P2A-EGFP or nirFAST fused to: histone H2B, mito (mitochondrial targeting motif), microtubule-associated protein (MAP) 4 and cytochrome b5 (Cb5) for endoplasmic reticulum membrane targeting, and labeled with 10 μM of HPAR-3OM (top row) (to assemble nirFAST680) or HPAR-3,5DOM (bottom row) (to assemble nirFAST715). Scale bars, 10 μm. Representative micrographs from three independent experiments (n > 13 cells). **b-c** Labeling of H2B-nirFAST in live and fixed cells. Nuclear fluorescence intensity and comparison of signal (nucleus) to background (cytosol) ratio (S/B) in live and fixed cells incubated with 10 µM of HPAR-3OM (to assemble nirFAST680) (**b**) or HPAR-3,5DOM (to assemble nirFAST715) (**c**), and imaged with identical microscope settings. On the graphs, each cell is color-coded according to the biological replicate it came from. The solid circles correspond to the mean of each biological replicate. The black line represents the mean ± SD of the two biological replicates. For nuclear fluorescence evaluation, 103 (respectively 93) fixed cells and 220 (respectively 83) live cells from two independent experiments were used for HPAR-3OM (respectively HPAR-3,5DOM). For S/B evaluation 103 (respectively 79) fixed cells and 189 (respectively 76) live cells from two experiments were used for HPAR-3OM (respectively HPAR-3,5DOM) analysis. Scale bars, 20 µm. (**d**) Confocal micrographs of live HeLa cells co-expressing ECFP fused to giantin, pFAST fused to a mitochondria targeting sequence (mito), mCherry fused to lyn11 (inner membrane-targeting motif) and nirFAST fused to H2B. Cells were labeled with 1 µM HMBR to assemble pFAST540 and 10 µM HPAR-3,5DOM to assemble nirFAST715. Micrographs in the cyan, green, red and near-infrared channels are represented as well as an overlay of the four channels. Representative micrographs from two independent experiments (n = 22 cells). Scale bars, 10 µm. (**e**) Imaging settings and spectral properties of the reporters used in (**d**). Absorption (dashed lines) and emission spectra (solid lines) of ECFP (from FPbase.org), pFAST540, mCherry (from FPbase.org) and nirFAST715 are shown. The four excitation wavelengths used in (**d**) are shown on the absorption spectra graph and the four spectral windows are indicated on the emission spectra graph as colored areas. See **Supplementary Table 8** for detailed imaging settings.

### Multicolor imaging in live cells

NIR fluorescent reporters allow multiplexed imaging through combination with visible fluorescent reporters that are spectrally distinct. Efficient multicolor imaging requires in particular efficient discrimination between NIR fluorescent reporters and red fluorescent reporters. We showed that, due to its red excitation and NIR emission, nirFAST715 (α_abs_ = 636 nm, α_em_ = 715 nm) can be efficiently separated from the red fluorescent protein mCherry (α_abs_ = 587 nm, α_em_ = 610 nm) by spectral discrimination, while maximizing the signal of the two reporters (**Supplementary Figure 9**). nirFAST715 was superior to nirFAST680 or frFAST670 that both show fluorescence bleed-through in the mCherry channel.

We also demonstrated the possibility of two-color imaging using spectrally orthogonal nirFAST715 and pFAST540 (α_abs_ = 481 nm, α_em_ = 540 nm) (**Supplementary Figure 10**). Fluorescence microscopy of HeLa cells expressing nirFAST and pFAST in the nucleus and mitochondria after treatment with 1 µM of HMBR and 10 µM of HPAR-3,5DOM showed green fluorescence only in mitochondria and NIR fluorescence only in the nucleus, demonstrating the selective labeling of pFAST with HMBR and nirFAST with HPAR-3,5DOM.

Finally, we showed that nirFAST715 could be combined with the enhanced cyan fluorescent protein (ECFP) (α_abs_ = 434 nm, α_em_ = 477 nm), pFAST540 and mCherry for achieving multicolor imaging of four cellular targets in live cells. We selectively visualize ECFP localized at the Golgi apparatus through fusion with giantin, the mitochondria-localized mito-pFAST (labeled with HMBR), the inner plasma membrane-anchored lyn11-mCherry and the nuclear H2B-nirFAST (labeled with HPAR-3,5DOM) in HeLa cells using four different excitation lines (405 nm, 488 nm, 561 nm, 639 nm respectively) and four detection windows (**Fig. 2d,e**), demonstrating that nirFAST715 could be combined with visible fluorescent reporters for spectral multiplexed imaging of up to four biological targets.

### Comparison with top-performing NIR FPs in cultured mammalian cells

Next, we compared nirFAST680 and nirFAST715 to emiRFP670 and miRFP713, two of the top-performing NIR FPs in cultured mammalian cells (**Fig. 3**). The two NIR FPs are characterized by fluorescence quantum yields comparable or lower than those of nirFAST680 and nirFAST715, however their molecular brightness *in vitro* is higher because of higher molar absorption coefficients. emiRFP670 absorbs maximally at 642 nm (with a molar absorption coefficient of 87,400 M^−1^.cm^−1^) and emits maximally at 670 nm (with a FQY of 14 %), while miRFP713 absorbs maximally at 690 nm (with a molar absorption coefficient of 99,000 M^−1^.cm^−1^) and emits maximally at 713 nm (with a FQY of 7 %) (**Fig. 3e** and **Supplementary Table 6**)^6^. In cells, the brightness of NIR FPs depends however on the intracellular concentration of biliverdin and its efficacy of incorporation. We thus compared the cellular brightness of emiRFP670 and miRFP713 to that of nirFAST680 and nirFAST715. The four proteins were co-expressed stoichiometrically with the green fluorescent protein EGFP using a polycistronic expression cassette containing a viral P2A sequence for ribosomal skipping during translation, allowing us to normalize the NIR fluorescent signal by the protein expression level using the EGFP signal. In HeLa cells, nirFAST680 was 2.5-fold brighter than emiRFP670 when exciting at 633 nm, although absorption at this wavelength is optimal for emiRFP670 and suboptimal for nirFAST680 (**Fig. 3a,b,e**). Similarly, nirFAST715 was 1.4-fold and 2-fold brighter than emiRFP670 and miRFP713, respectively, when exciting at 633 nm (**Fig. 3c-e**). Photostability studies in cells showed furthermore that nirFAST680 and nirFAST715 displayed excellent photostability, comparable to that of emiRFP670 (**Supplementary Figure 11**). The possibility to exchange bleached chromophores with new ones might explain the high photostability, as previously reported for other non-covalent systems^15–17^. Overall, this set of experiments showed that nirFAST was an excellent alternative to NIR FPs for efficient protein imaging in cultured mammalian cells.

**Fig. 3.**
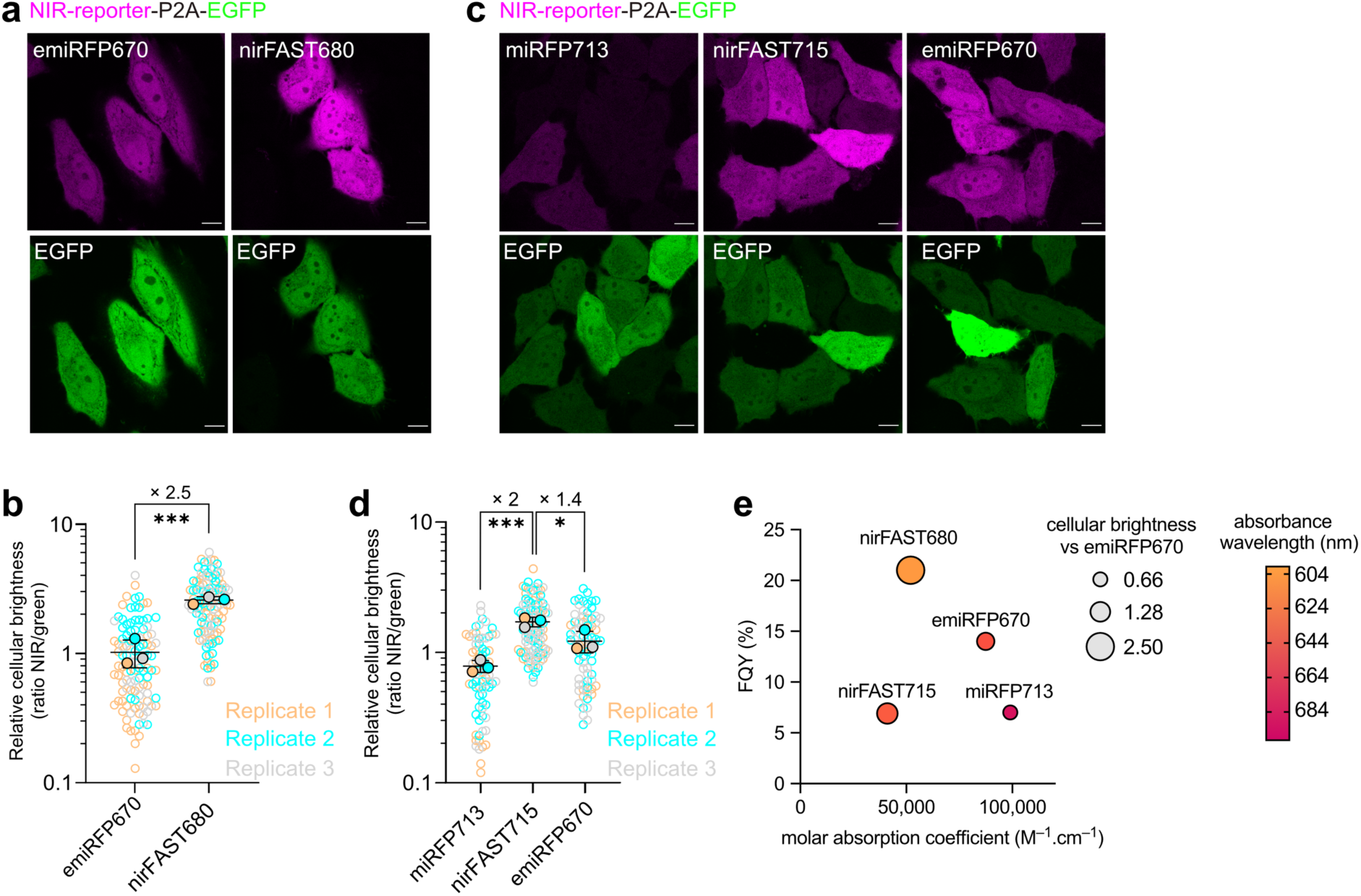
Comparison of nirFAST with top-performing emiRFP670 and miRFP713. (**a**) Confocal micrographs of Hela cells expressing, nirFAST-P2A-EGFP (labeled with 10 µM HPAR-3OM to assemble nirFAST680) and emiRFP670-P2A-EGFP. Scale bars, 10 µm. Representative micrographs of n > 100 cells from three independent experiments. Identical microscope settings were used for enabling side-by-side comparison. (**b**) Relative cellular brightness of nirFAST680 and emiRFP670 computed by normalizing the fluorescence of the NIR reporters with the green fluorescence of the stoichiometrically expressed EGFP (ratio NIR/green). Each cell is color-coded according to the biological replicate it came from. The solid circles correspond to the mean of each biological replicate. The solid circles correspond to the mean of each biological replicate. The black line represents the mean ± SD of the three biological replicates. n = 115 (respectively n = 108) live cells from three independent experiments were used for nirFAST680 (respectively emiRFP670). Unpaired two-tailed t-test assuming equal variance was used to compare the two distributions (***P = 0.0008). **(c)** Confocal micrographs of Hela cells expressing miRFP713-P2A-EGFP, nirFAST-P2A-EGFP (labeled with 10 µM HPAR-3,5DOM to assemble nirFAST715) and emiRFP670-P2A-EGFP. Scale bars, 10 µm. Representative micrographs of n > 70 cells from three independent experiments. Identical microscope settings were used for enabling side-by-side comparison. Unpaired two-tailed t-test assuming equal variance was used to compare miRFP713 and nirFAST715 distributions (***P = 0.0006, *P = 0.0340). (**d**) Relative cellular brightness of miRFP713, nirFAST715 and emiRFP670 computed by normalizing the fluorescence of the NIR reporters with the green fluorescence of the stoichiometrically expressed EGFP (ratio NIR/green). Each cell is color-coded according to the biological replicate it came from. The solid circles correspond to the mean of each biological replicate. The black line represents the mean ± SD of the three biological replicates. n = 106 (respectively n = 78 and n = 74) live cells from three independent experiments were used for nirFAST715 (respectively miRFP713 and emiRFP670). (**e**) Photophysical properties of emiRFP670^6^, nirFAST680, miRFP713^6^ and nirFAST715. FQY of these reporters are plotted against their molar absorption coefficient. Each data point size is scaled to the cellular fluorescence brightness of the corresponding fluorescent reporter and the color is indicative of the absorption wavelength. See **Supplementary Table 8** for detailed imaging settings.

### Subdiffraction imaging of nirFAST fusions with stimulated emission depletion nanoscopy

Super-resolution microscopy techniques such as stimulated emission depletion (STED) microscopy allow for imaging with spatial resolution beyond the diffraction limit. In STED, super-resolution is achieved by shining a depletion doughnut-shaped laser beam right after excitation^18^. This depletion laser beam switches off fluorophores in the doughnut region through stimulated emission, leaving only the fluorophores at the center in an emissive state. STED requires bright and photostable fluorophores to achieve improved resolution. As nirFAST efficiently formed a bright assembly (nirFAST600) with HBR-3,5DOM (**Supplementary Figure 12**), a chromophore which previously allowed STED nanoscopy of pFAST-tagged proteins using a green excitation laser and a 775 nm depletion laser^9^, we first demonstrated the STED compatibility of nirFAST using this chromophore. STED microscopy of microtubule-associated MAP4-nirFAST in HeLa cells treated with HBR-3,5DOM allowed us to demonstrate that nirFAST600 was an efficient STED label for the visualization of fine cellular structures in live cells (**Fig. 4a**), with improved resolution in comparison with regular confocal microscopy (**Fig. 4b**). Using a parameter-free image resolution estimation method based on image partial phase autocorrelation^19^, the resolution enhancement over confocal microscopy was estimated to be 1.6-fold (**Fig. 4c**).

**Fig. 4.**
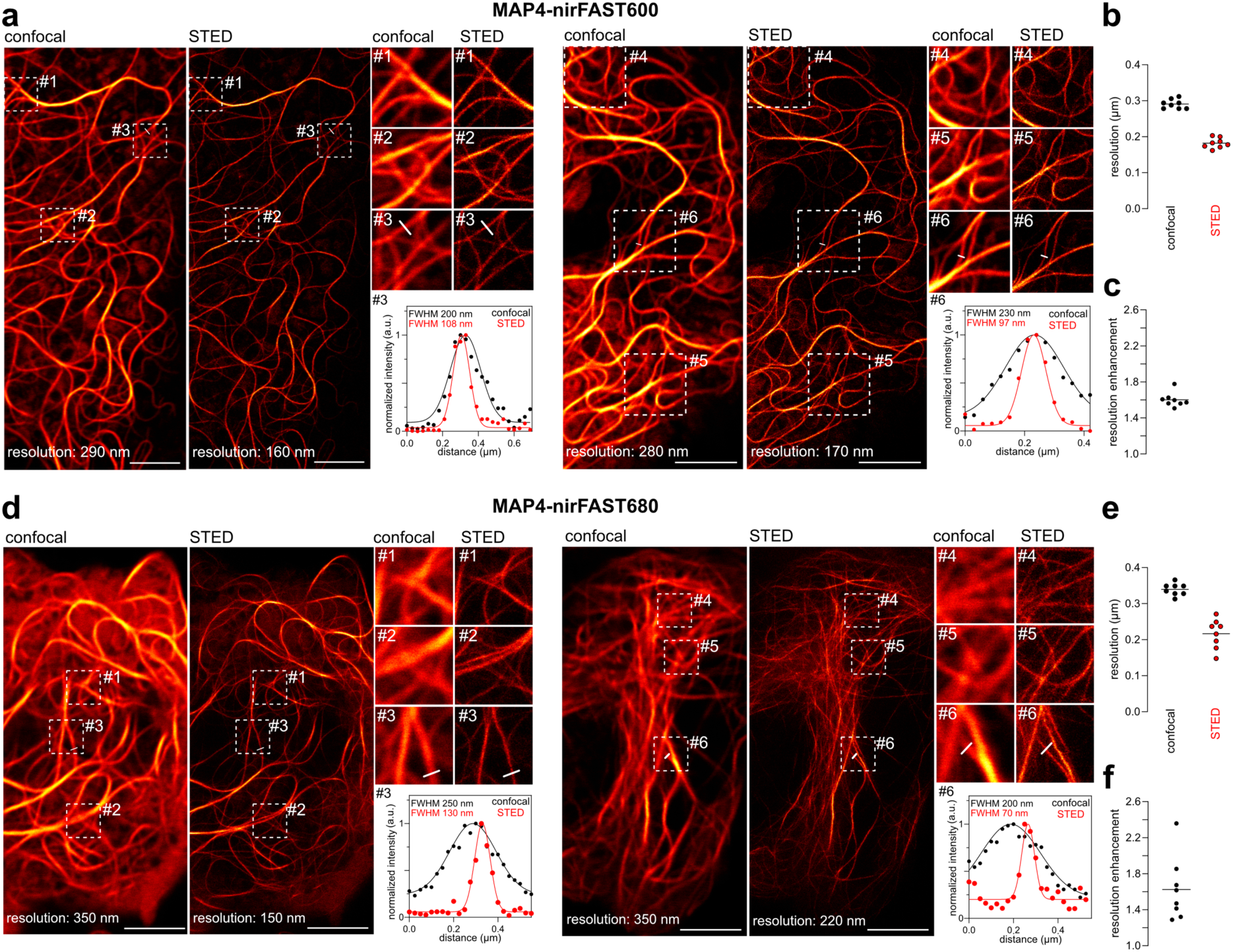
STED imaging of nirFAST-tagged proteins in live mammalian cells. Confocal and STED micrographs of two live HeLa cells expressing nirFAST fused to MAP4 and labeled with 10 µM of HBR-3,5DOM (to assemble nirFAST600) (**a-c**) or 10 µM of HPAR-3OM to assemble (nirFAST680) (**d-f**). For each cell, three regions of interest were selected for close-up comparison between confocal and STED. Line profile across microtubule filaments from one close-up was used to compare gain in resolution between confocal and STED. Graph shows raw data of line profile from confocal and STED images (points), in addition to the corresponding gaussian fit (line). Micrographs are representative of n = 8 cells from two independent experiments for HBR-3,5DOM (**a**) and HPAR-3OM (**d**). Scale bars, 5 µm. (**b**) Comparison of resolution of confocal and STED MAP4-nirFAST680 micrographs using image decorrelation analysis^19^ (n = 8 cells). (**c**) Enhancement of resolution of MAP4-nirFAST600 micrographs, calculated as the ratio between confocal resolution and STED resolution from (**b**). (**e**) Comparison of resolution of confocal and STED MAP4-nirFAST680 micrographs using image decorrelation analysis^19^ (n = 8 cells). (**f**) Enhancement of resolution of MAP4-nirFAST680 micrographs, calculated as the ratio between confocal resolution and STED resolution from (**e**). See **Supplementary Table 8** for detailed imaging settings and methods section for image decorrelation analysis parameters.

Although FR and NIR light are less toxic and more compatible with biological imaging, achieving high resolution with FR and NIR reporters is more challenging because of the wavelength-dependence of spatial resolution. In the context of STED, the use of FR and NIR reporters can also be hindered by their possible re-excitation by the 775 nm depletion laser. In this context, we reasoned that nirFAST680 should be well suited for STED nanoscopy because of its excellent photostability, brightness and spectral properties. With a 604 nm excitation peak, nirFAST680 is less prone to re-excitation by the 775 nm depletion beam compared to other NIR reporters with more red-shifted absorption, which should thus increase resolution. In addition, nirFAST680 is efficiently excited at 640 nm, a more favorable wavelength for imaging at high power in living cells. Accordingly, we successfully imaged fine microtubule structures labeled with MAP4-nirFAST680 in live HeLa cells using STED nanoscopy (**Fig. 4d**), achieving a 1.6-fold resolution enhancement over confocal microscopy (**Fig. 4e,f**,). Overall, this set of experiments showed that nirFAST600 and nirFAST680 were excellent STED probes for sub-diffraction imaging in live cells.

### Imaging of nirFAST in chicken embryo tissues

Next, we investigated the suitability of nirFAST for protein labeling in multicellular organisms using chicken embryo tissues as model. We expressed nirFAST and frFAST in each side of the neural tube (using a sequential *in ovo* bilateral electroporation strategy) for direct comparison of the two proteins within the very same sample. After dissection of the neural tube, the sample was cultured in an open-book configuration, and labeled with either HPAR-3OM or HPAR-3,5DOM. Simultaneous imaging showed that, with HPAR-3OM, labeling was achieved earlier than 30 minutes following fluorogen addition with robust NIR signal detected for both frFAST and nirFAST (**Fig. 5a,b**), confirming efficient permeation of the fluorogen across cell membranes. Fluorescence appears earlier for nirFAST, in agreement with a more efficient binding of HPAR-3OM. Labeling with HPAR-3,5DOM showed high labeling efficiency of nirFAST but poor signal for frFAST (**Fig. 5c**), in agreement with their relative properties with this fluorogen (**Supplementary Table 1**). Comparison of the in-tissue brightness of nirFAST715 and emiRFP670 using the same protocol (**Fig. 5d**) showed that the two proteins displayed comparable fluorescence in chicken embryo tissues, in agreement with the behavior observed in live cultured mammalian cells.

**Fig. 5.**
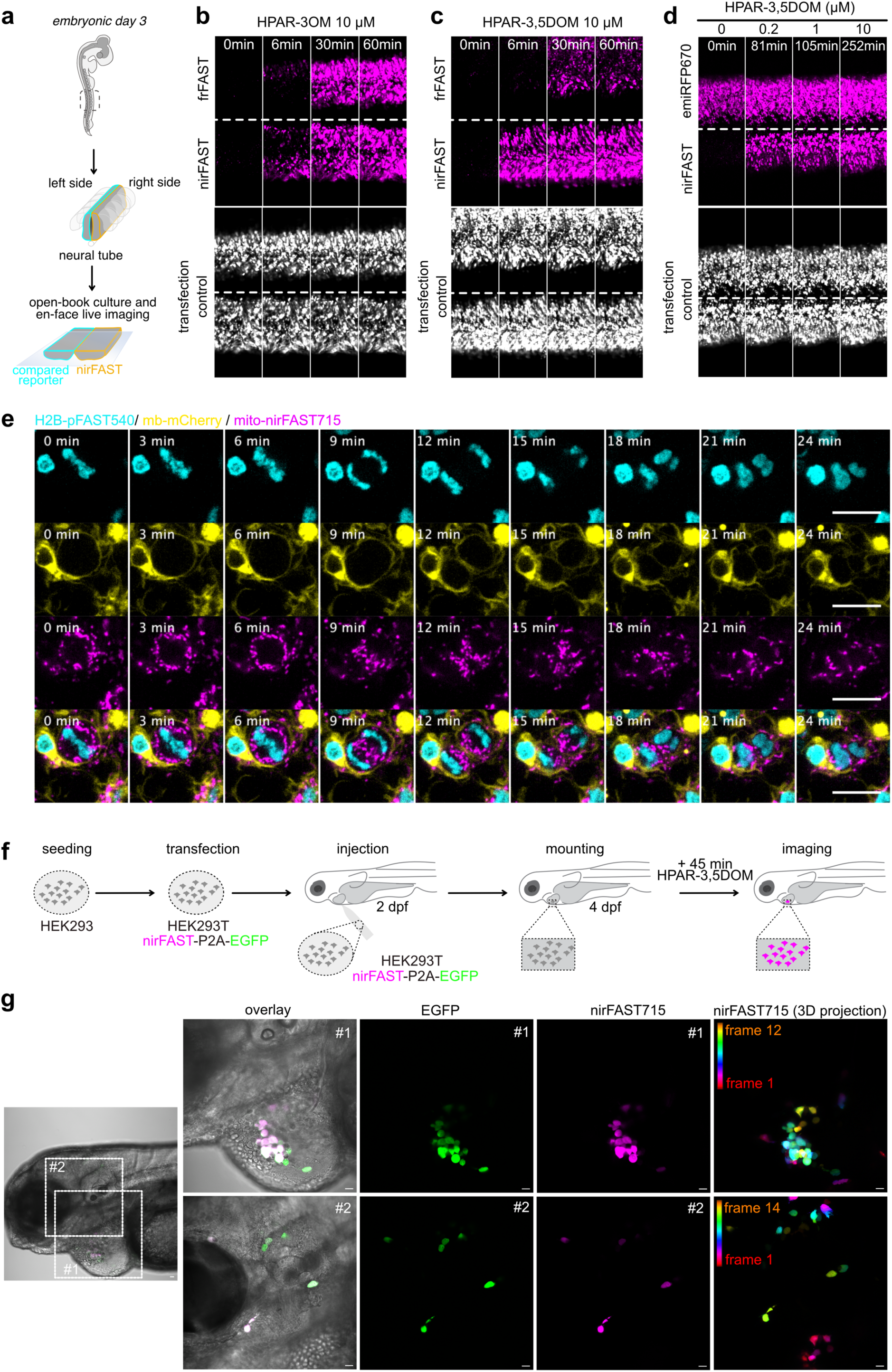
Selective imaging of nirFAST in chicken embryo and zebrafish larvae. (**a-c**) Plasmids encoding nirFAST-P2A-EGFP and frFAST-P2A-EGFP were electroporated in each side of the neural tube in ovo at embryonic day 2. 24 h later, embryos with homogenous bilateral reporter expression in the neural tube were dissected and imaged upon addition of 10 µM of HPAR-3OM or HPAR-3,5DOM by timelapse spinning disk microscopy. Scale bars 100 µm. Representative timelapse micrographs of n = 3 independent experiments. (**d**) Plasmids encoding nirFAST-P2A-EGFP and emiRFP670-P2A-EGFP were electroporated in each side of the neural tube in ovo at embryonic day 2. 24 h later, embryos with homogenous bilateral reporter expression in the neural tube were dissected and the neuroepithelium was imaged in en-face view by time-lapse spinning disk microscopy upon sequential addition of 0.2, 1 and 10 μM HPAR-3,5DOM. Scale bars 100 µm. Representative micrographs of n = 3 independent experiments. (**e**) Plasmids encoding H2B-pFAST (cyan), memb-mCherry (yellow) and mito-nirFAST (magenta) were electroporated in the neural tube in ovo, at embryonic day 2 (E2, HH stage 13–14). 24 h later, embryos were dissected, and the neuroepithelium was imaged in en-face view in the presence of 1 μM HMBR (to assemble pFAST540) and 10 μM HPAR-3,5DOM (to assemble nirFAST715) using a spinning disk microscope (see also **Supplementary Movie 2**). Scale bars, 10 µm. Representative micrographs of n = 3 independent experiments. (**f,g**) Mammalian HEK293T were transfected with plasmid encoding nirFAST-P2A-EGFP. After 24h, they were injected near the heart of 2 dpf zebrafish larvae. Larvae with green fluorescence signal were selected and imaged at 4 dpf after 45 min incubation with HPAR-3,5DOM (to assemble nirFAST715). Representative micrographs of n = 3 independent experiments. Scale bars, 20 µm. See **Supplementary Table 8** for detailed imaging settings.

Next, we performed multicolor imaging in chicken embryo tissues. We simultaneously followed the dynamics of nuclear H2B-pFAST540, membrane-anchored mCherry and mitochondria-localized nirFAST715 during cell division in the neural tube by en-face timelapse imaging of the neuroepithelium (**Fig. 5e**, **Supplementary Movie 2**). Labeling of mitochondria with nirFAST715 allowed us to efficiently follow their dynamics during cytokinesis: in metaphase, mitochondria were distributed in the cytoplasm around the metaphase plate; moving to anaphase, mitochondria were recruited at the cell equator at the site of cleavage furrow and depleted at the cell poles, allowing accurate inheritance of mitochondria between the two daughter cells.

These experiments demonstrated the excellent suitability of nirFAST for imaging biological processes in complex tissues and highlighted the interest of reporters with NIR-shifted spectral properties for efficient multiplexed imaging.

### Imaging of nirFAST in zebrafish embryo

Widely used model for the study of gene function, zebrafish is also an interesting model for cancer-related studies, in particular for investigating cell invasion and metastasis. Injection of cancer cells in zebrafish larvae allows the monitoring of cell growth and proliferation as well as their interaction with their environment by fluorescence imaging^20,21^. In this context, we reasoned that the NIR excitation and emission of nirFAST would make it an interesting alternative to more blue-shifted reporters because of the low autofluorescence of zebrafish tissues in this region of the spectrum. To demonstrate our ability to detect nirFAST-expressing cells in zebrafish larvae, we injected HEK293T cells co-expressing nirFAST and EGFP, near the heart of larvae at 2 dpf (day post fertilization) stage (**Fig. 5f,g**). Larvae were then imaged two days later by confocal microscopy. After 45 min of treatment with HPAR-3,5DOM, individual cells emitting green and NIR fluorescence were efficiently detected mainly near the heart. Some cells were also detected from areas close to the inner ear, suggesting cell migration. This experiment showed that nirFAST was well suited for labeling xenografts of mammalian cells injected in zebrafish larvae.

### A green-NIR fluorescent cell cycle indicator

Next, we took advantage of the orthogonality of nirFAST715 and pFAST540 to design a green-NIR fluorescent ubiquitination cell cycle indicator (FUCCI). Based on the fusion of two spectrally orthogonal fluorescent reporters to fragments of Geminin and Cdt1, two cell-cycle regulators whose levels inversely oscillate during cell cycle, FUCCI indicators enable to identify the different phases of the cell cycle^22^. While the use of FP-based FUCCI was generalized to different types of organisms, the study of short cell cycles remains highly challenging because of the slow fluorescence maturation of FP. An alternative relies on the use of fast activated fluorescent chemogenetic reporters such as FAST variants^23^. We reasoned that the use of pFAST and nirFAST, which both show quasi-instantaneous labeling (provided that their fluorogens are present), could allow the design of cell cycle indicator (hereafter named ^FAST^FUCCI) circumventing the limitation of FP-based FUCCI. nirFAST was fused to the N-terminus domain of zCdt1(1-190) and pFAST was fused to the N-terminus of zGeminin(1-100). Cdt1 and Geminin are substrates of the two complexes SCF^Skp2^ and APC^Cdh1^, which display cell-cycle phase-dependent activities. These two complexes are involved in the tight control of DNA replication, which occurs during the S phase of the cycle. In particular, geminin is degraded during G1 while Cdt1 is degraded during S/G2/M. Thus, fluorescent reporters fused to fragments of Cdt1 and Geminin can directly report the accumulation or degradation of these two proteins, allowing delineation of cell cycle phases^22^. Long-term timelapse imaging of HEK293T cells showed that ^FAST^FUCCI enabled to report on the different stages of the cell cycles, while having no deleterious effect on cell viability and division (**Supplementary Figure 13, Supplementary Movie 3**). The observation of asynchronized cells revealed several types of transitions (**Fig. 6**, **Supplementary Movie 4**). Cells right after division displayed NIR fluorescence, in agreement with the accumulation of Cdt1 during G1 phase. Over time, the NIR signal decreased while the green fluorescence increased with an overlap of the two signals, revealing transition to the S phase. Full disappearance of the NIR fluorescence and constant green fluorescent signal allowed us to identify cells having reached the G2 phase. ^FAST^FUCCI thus is a suitable indicator for delineating different phases of the cell cycle with green and NIR readout.

**Fig. 6.**
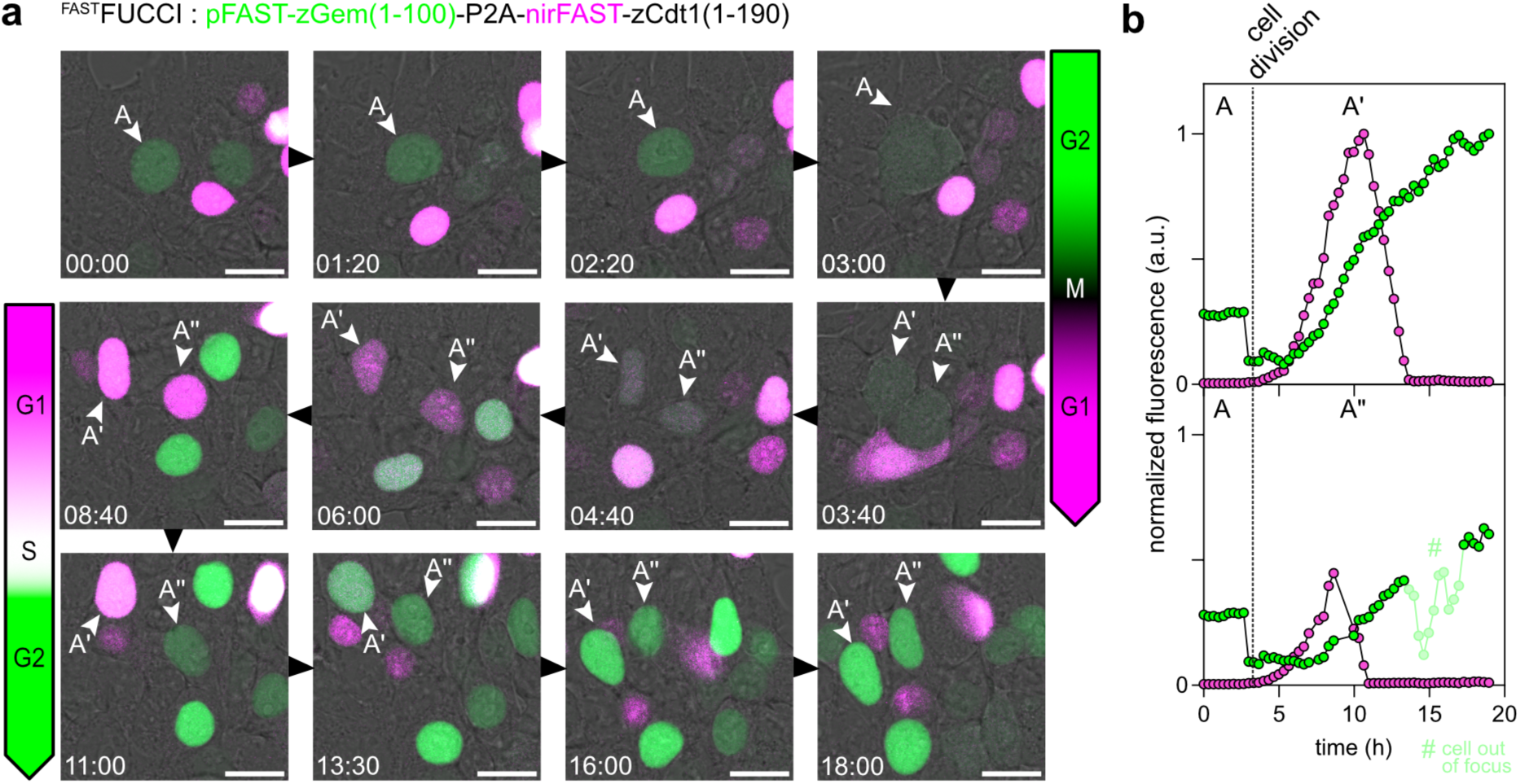
^FAST^FUCCI, a green-NIR fluorescent chemogenetic cell-cycle indicator. (**a**) HEK293 cells expressing ^FAST^FUCCI and treated with 1 µM HMBR (to assemble pFAST540) and 10 µM HPAR-3,5DOM (to assemble nirFAST715) were imaged by time-lapse confocal microscopy (1 frame every 20 min) for 24 h. Relevant timepoints of designated field of view are shown to highlight transitions between the different cell cycle stages. Representative micrographs from three independent experiments (see also **Supplementary Movie 4**). (**b**) Fluorescence intensity evolution of tracked cell (cell A), which upon mitosis gives cells A’ and cells A”, is shown over time. Cell tracking and lineage was achieved through Trackmate plugin (Fiji)^41,42^. See **Supplementary Table 8** for detailed imaging settings.

### Control of protein proximity using bisected nirFAST

Bisection of fluorescent reporters in two complementary fragments is a general approach for the development of methods to image or control the proximity of proteins^24^. Using the split site 114-115 previously used for splitting FAST variants^25–27^, we split nirFAST into two complementary fragments dubbed nirFAST_1-114_ and nirFAST_115-125_. We evaluated the complementation of this split system (dubbed split-nirFAST) by fusing nirFAST_1-114_ to the C-terminus of the FKBP-rapamycin-binding domain (FRB) of the mechanistic target of rapamycin (mTOR), and nirFAST_115-125_ to the C-terminus of the FK506-binding protein (FKBP). As FKBP and FRB interact in the presence of rapamycin^28^, the quantification of NIR fluorescence without and with rapamycin allowed us to quantify the efficiency of complementation of split-nirFAST in the absence and in the presence of interaction at various fluorogen concentrations (**Supplementary Figure 14**). Flow cytometry analysis revealed that split-nirFAST displayed efficient complementation in cells with both HPAR-3OM and HPAR-3,5DOM regardless of the presence of rapamycin (**Supplementary Figure 14a,b,e**). Similarly to what was observed for other split-FASTs, self-complementation of the ternary fluorescent assembly increases with fluorogen concentration. High self-complementation of split-nirFAST680 and split-nirFAST715 in cells was also observed by confocal microscopy (**Supplementary Figure 14c,d,f,g**).

Although high self-complementation precluded the use of split-nirFAST680 or split-nirFAST715 as reporter of protein-protein interactions, this behavior could be leveraged to chemically induce protein proximity using the recently described CATCHFIRE (chemically assisted tethering of chimera through fluorogenic induced recognition) approach. CATCHFIRE uses ^FIRE^mate (a.k.a. pFAST_1-114_) and ^FIRE^tag (a.k.a. pFAST_115-125_) as dimerization domains that can interact together in the presence of various fluorogens (a.k.a. ‘matches’)^27^. nirFAST_115-125_ and ^FIRE^tag only differ by one mutation at position 117 (S117R). Characterization of full-length nirFAST^S117R^ showed that this variant conserved good affinity for HPAR fluorogens (**Supplementary Table 7**). We showed that the replacement of nirFAST_115-125_ by ^FIRE^tag did not hamper fluorogen-induced complementation. Co-expression in HEK293T cells of FRB-nirFAST_1-114_ along with mCherry-nirFAST_115-125_ or mCherry-^FIRE^tag, respectively allowed us to demonstrate that fluorogen-induced complementation was more efficient with ^FIRE^tag than with nirFAST_115-125_ (**Supplementary Figure 15**). Optimal fluorogen-induced complementation was observed when combining nirFAST_1-114, FIRE_tag and HPAR-3,5DOM (**Supplementary Figure 15b,f-h**).

This set of experiments suggested the possibility to control the proximity of two proteins with a NIR fluorescence readout using nirFAST_1-114_ and ^FIRE^tag as dimerizing domains and HPAR fluorogens as fluorogenic inducers of proximity, and we called the resulting system ^nir^CATCHFIRE. We fused nirFAST_1-114_ (hereafter called ^nir-FIRE^mate) to the C-terminus of the N-terminal domain of the mitochondrial outer membrane protein TOM20 (TOM20_1-34_) and co-expressed it with EGFP-^FIRE^tag in HeLa cells (**Fig. 7a**). Correct localization of ^nir-FIRE^mate fusion at the mitochondria was assessed through insertion of the enhanced cyan fluorescent protein (ECFP). In the absence of fluorogen, EGFP-^FIRE^tag displayed cytosolic localization, demonstrating the absence of complementation of ^nir-FIRE^mate and ^FIRE^tag in the absence of fluorogen. Addition of HPAR-3OM or HPAR-3,5DOM induced the rapid translocation of the green fluorescent EGFP-^FIRE^tag from the cytosol to the surface of the mitochondria and the formation of the ternary complex between ^nir-FIRE^mate, ^FIRE^tag and the fluorogen, as evidenced by an increase of the green fluorescence signal and NIR signal at the outer membrane of mitochondria in timelapse confocal microscopy experiments (**Fig. 7b,d and Supplementary Movies 5 and 6**). NIR signal reached 50 % of maximum fluorescence two to three minutes after fluorogen addition, demonstrating fast fluorogen-induced association of ^nir-FIRE^mate and ^FIRE^tag (**Fig. 7c,e**).

**Fig. 7.**
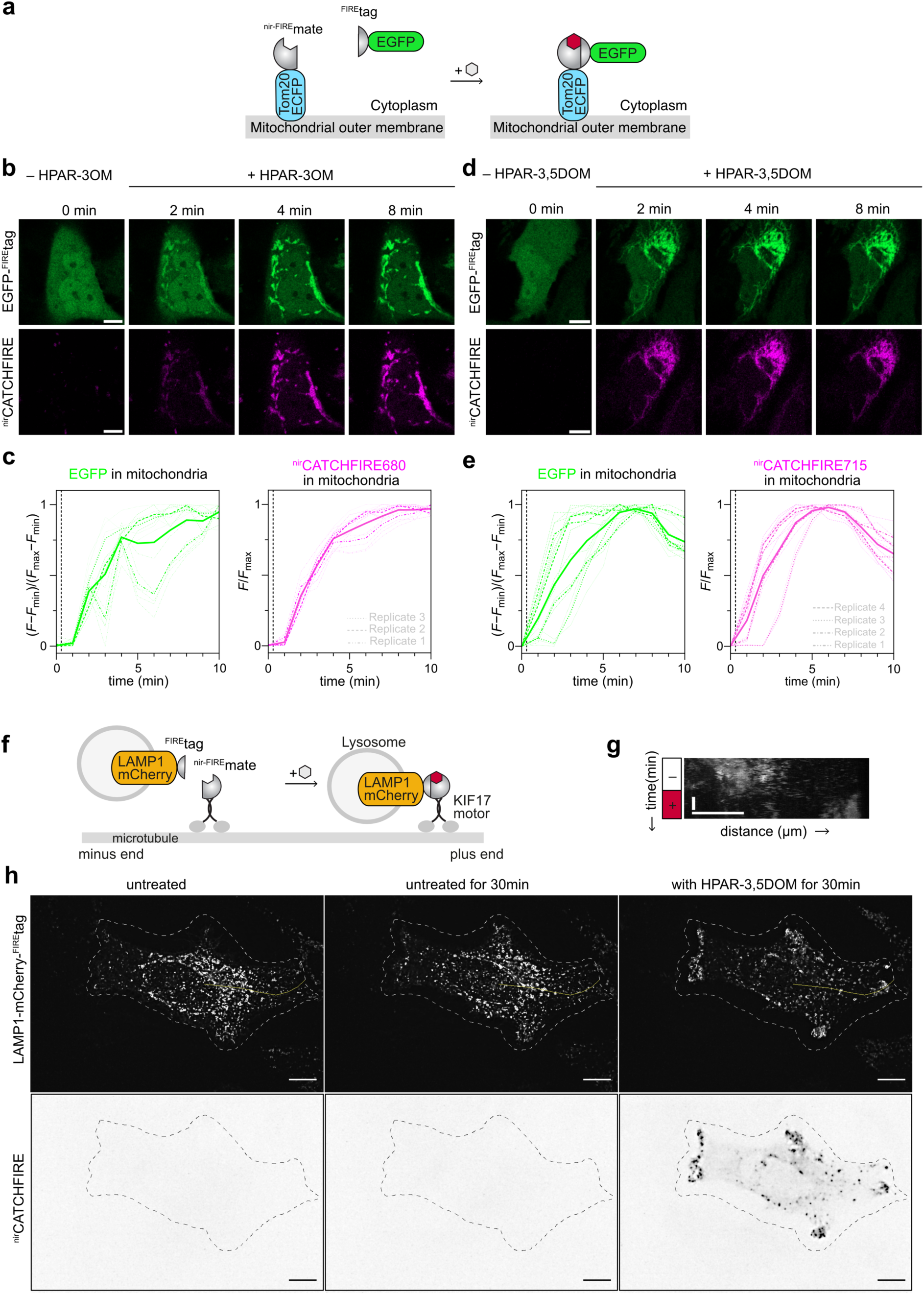
^nir^CATCHFIRE: a fluorogenic chemically induced dimerization tool with NIR fluorescence readout. (**a**) Schematic illustrating the fluorogen-induced interaction between EGFP-^FIRE^tag and Tom20-ECFP-^nir-FIRE^mate for chemically induced recruitment at the outer mitochondrial membrane. (**b-e**) HeLa cells co-expressing EGFP-^FIRE^tag together with Tom20-ECFP-^nir-FIRE^mate were treated with 10 µM of either HPAR-3OM (**b,c**) or HPAR-3,5DOM (**d,e**), and imaged by timelapse confocal microscopy. (**b,d**) Representative confocal micrographs before and after addition of the fluorogen (see also **Supplementary Movies 5 and 6**). Scale bar 10 µm. (**c,e**) Temporal evolution of the mitochondrial EGFP and ^nir^CATCHFIRE fluorescence of (**c**) n = 15 cells (from three independent experiments) and (**d**) n = 14 cells (from four independent experiments). The dashed line indicates fluorogen addition. (**f**) Schematic illustrating how fluorogen-induced interaction between LAMP1–mCherry–^FIRE^tag and ^nir-FIRE^mate–KIF17 allows the chemically induced anterograde transport of lysosomes. (**g**) HeLa cells coexpressing LAMP1–mCherry–^FIRE^tag and ^nir-FIRE^mate–KIF17 were imaged by spinning disk microscopy for 30 min, then HPAR-3,5DOM was added. Representative micrographs of each step (see also **Supplementary Movie 7**). Experiments were repeated three times with similar results. Scale bars 10 µm. (**h**) Kymograph showing the LAMP1–mCherry–^FIRE^tag along the yellow line in (**e**) over time. See **Supplementary Table 8** for detailed imaging settings.

To demonstrate the possibility of controlling cellular process, we applied ^nir^CATCHFIRE to control the positioning of lysosomes (**Fig. 7f**). ^nir-FIRE^mate was fused to the C-terminus of the motor domain of kinesin-like KIF17, while ^FIRE^tag was fused to lysosomal-associated membrane protein (LAMP)1 at the surface of lysosomes. Co-expression of the two proteins in live cells allowed us to control lysosome positioning. Addition of HPAR-3,5DOM led to the displacement of lysosomes from the perinuclear region to the cell periphery, demonstrating the successful fluorogen-induced recruitment of the molecular motor (**Fig. 7g-h** and **Supplementary Movie 7**).

This set of experiments opens exciting prospects for controlling the localization and motion of proteins and organelles using the ^nir^CATCHFIRE technology. Its NIR fluorescence properties offer the additional advantage of possibly reporting additional biological events using green or red-emitting reporters.

## DISCUSSION

NIR fluorescent reporters allow observations in living tissues with high contrast and lower phototoxicity. The need for reporters that fully maximize the advantages of the NIR spectral window drove the development of a toolbox of NIR FPs for various applications in bioimaging^1–6^. In this work, we present the development of nirFAST, a NIR fluorescent chemogenetic reporter, made of a 14 kDa protein tag able to tightly bind and stabilize a new fluorogenic push-pull chromophore called HPAR-3,5DOM displaying efficient permeation across cell membranes. To engineer nirFAST, we took advantage of the bimodular nature of chemogenetic systems to conduct a concerted strategy of fluorogen and tag engineering. By altering the chromophore structure and refining the protein structure through two rounds of directed evolution, we maximized NIR fluorescence brightness, and obtained spectral properties finely adjusted to excitation at 633 nm or 640 nm readily available on most microscope setups.

Although nirFAST was engineered and optimized to take advantage of the benefits of NIR excitation and emission, efficient binding and fluorescence activation can also be achieved with HPAR-3OM yielding far-red fluorescence (nirFAST680), and HBR-3,5DOM yielding red fluorescence (nirFAST600). The fluorescence of nirFAST can thus be finely tuned from red to NIR only by changing the fluorogen, highlighting the versatility and high tunability of chemogenetic systems.

nirFAST680 and nirFAST715 were shown to be brighter than the spectrally equivalent top-performing NIR FPs emiRFP670 and miRFP713 in cultured mammalian cells. As their fluorescence is only reliant on the addition of highly permeant exogenous fluorogens, they can be used as effective alternatives of NIR FP in a variety of biological contexts where scarce levels of biliverdin hamper their use (e.g. neurons^29^). nirFAST displays the additional advantage of being half the size of the top-performing NIR FPs, opening interesting prospects for labeling proteins with robust fluorescence and minimal perturbation. Finally, full labeling of nirFAST in cells is achieved within seconds of fluorogen addition, indicating that the rate of labeling is primarily constrained by fluorogen cell uptake and diffusion. This means that in cells pre-treated with the fluorogen, labeling is nearly instantaneous, which should enable the reporting of dynamic and rapid biological processes such as protein neosynthesis and turnover. This contrasts with biliverdin-based NIR FPs, which can be limited by their longer maturation time due to slower covalent incorporation of biliverdin^3^.

In this study, we showed that nirFAST is advantageous for simultaneous imaging of several biological targets, as it can be combined with various fluorescent reporters emitting in the visible. In addition to being unmistakably distinguishable from widely used red-emitting fluorescent reporters due to its red-shifted spectral properties, nirFAST is also orthogonal to pFAST, another chemogenetic reporter of the FAST family capable of yielding green fluorescence. nirFAST allowed spectral multiplexed imaging in cells as well as in chicken embryo tissues, where the dynamics of three subcellular proteins were simultaneously monitored during cell division.

The orthogonality of nirFAST and pFAST allowed the design of a FUCCI-like cell cycle indicator delineating cell-cycle stages with green and NIR fluorescence readout. The photostability and low toxicity of the ^FAST^FUCCI indicator allowed the visualization of M/G1 as well as G1/S/G2 transitions over long periods of time while having the red channel readily available for monitoring additional processes. Because of the quasi-instantaneous labeling of nirFAST and pFAST, the ^FAST^FUCCI indicator should be ideal for the monitoring of very short cell cycles, such as those in early stages of embryogenesis^23^.

In addition of requiring less toxic light for imaging, NIR fluorescent reporters facilitate imaging in biological tissues, which are less autofluorescent in this spectral window, enabling thus observations without interference from inherent fluorescent structures. In this context, nirFAST enabled unambiguous high contrast observation of injected cells inside zebrafish larvae. In these experiments we used non-tumorigenic HEK293T cells as a proof of concept of the successful labeling of injected cells inside zebrafish larvae. The advantageous spectral properties of nirFAST should allow its combination with the variety of existing EGFP-expressing transgenic zebrafish lines for the study of angiogenesis or response to anti-cancer drugs of tumorigenic injected cells.

While red, FR and NIR lights are associated with better in-depth penetration and lower phototoxicity, they are often linked to lower spatial resolution in microscopy. In this study, we showed that the excellent spectral properties, photostability and brightness of nirFAST600 and nirFAST680 make them well-suited reporters for STED, and enable live-cell subdiffraction imaging of fine cellular structures such as microtubules with a 1.6-fold resolution enhancement.

Finally, we used nirFAST to design a fluorogenic chemically inducible dimerization tool with NIR fluorescence readout. The bisection of members of the FAST family previously led to chemogenetic tools enabling the functional study of protein-protein interactions^10,23,25–27^. Bisection led to either systems displaying low self-complementation in the presence of the fluorogen, well suited for reporting protein-protein interactions^26^ and contact sites between organelles^30,31^ or to systems displaying high self-complementation in the presence of the fluorogen, which were used to develop CATCHFIRE (chemically assisted tethering of chimera through fluorogenic induced recognition)^27^, a chemically induced dimerization tool with intrinsic fluorescence readout for controlling and visualizing protein proximity. Bisection of nirFAST allowed us to generate a CATCHFIRE system with NIR fluorescence readout, dubbed ^nir^CATCHFIRE, in which HPAR-3OM and HPAR-3,5DOM act as fluorogenic molecular glues promoting the dimerization of the two fragments. This system can thus be used to chemically control and also visualize the proximity of two proteins fused to the two fragments.

Either used in its full-length or split version, nirFAST opens exciting prospects for the development of novel single wavelength or FRET-based tunable biosensors with NIR readout for unravelling biological processes in complex model organisms. We expect the promising properties of nirFAST to make it an interesting addition to the existing toolbox of NIR fluorescent reporters.

## METHODS

### Organic synthesis

#### General

Commercially available reagents were used as obtained. ^1^H and ^13^C NMR spectra were recorded at 300K on a Bruker AM 300 spectrometer; chemical shifts are reported in ppm with protonated solvent as internal reference; coupling constants *J* are given in Hz. Mass spectra were performed by the Service de Spectrométrie de Masse de Chimie Paris Tech (France). The synthesis of HPAR-3OM^10^, HBR-3,5DOM^14^, HBR-3,5DM^14^ and HMBR^13^ was previously reported. These fluorogens are commercially available from the Twinkle Factory under the name ^TF^Poppy, ^TF^Coral, ^TF^Amber and ^TF^Lime.

#### HPAR-3,5DOM ((Z)-5-((E)-3-(4-hydroxy-3,5-dimethoxyphenyl)allylidene)-2-thioxothiazolidin-4-one

To a stirred solution of rhodanine (133 mg, 1.0 mmol) and 4-hydroxy-3,5-dimethoxycinnamaldehyde (208 mg, 1.0 mmol) in ethanol (3 mL) was added 4-dimethylaminopyridine (12 mg, 0.11 mmol). The solution was stirred at reflux for 20 h. Solution was neutralized by adding 1N HCl. After cooling to 4°C and standing overnight, the precipitate was filtered and the crude solid was washed with EtOH/H_2_O 1:5. HPAR-3,5DOM was obtained as a red powder (303 mg, 94% yield). ^1^H NMR (300 MHz, CD_3_SOCD_3_, δ in ppm): 13.55 (s, 1H), 9.08 (s, 1H), 7.32 (d, *J* = 11.7 Hz, 1H), 7.21 (d, *J* = 15.0 Hz, 1H), 6.99 (s, 2H), 6.91 (dd, *J* = 11.7, 15.0 Hz, 1H), 3.82 (s, 6H). ^13^C NMR (75 MHz, CD_3_SOCD_3_, δ in ppm): 195.2, 168.7, 148.1(2C), 146.3, 138.5, 133.1, 126.2, 124.6, 121.2, 106.1(2C), 56.2(2C). MS (ESI): m/z 322.3 [M–H]^−^ (calculated mass for [C_14_H_12_NO_4_S_2_]^−^: 322.3).

#### Biology

The presented research complies with all relevant ethical regulations

#### General

Synthetic oligonucleotides used for cloning were purchased from Integrated DNA Technology. PCR reactions were performed with Q5 polymerase (New England Biolabs) in the buffer provided. PCR products were purified using QIAquick PCR purification kit (QIAGEN). DNase I, T4 ligase, fusion polymerase, Taq ligase and Taq exonuclease were purchased from New England Biolabs and used with accompanying buffers and according to the manufacturer’s protocols. Isothermal assemblies (Gibson Assembly) were performed using a homemade mix prepared according to previously described protocols^32^. Small-scale isolation of plasmid DNA was conducted using a QIAprep miniprep kit (QIAGEN) from 2 mL overnight bacterial culture supplemented with appropriate antibiotics. Large-scale isolation of plasmid DNA was conducted using the QIAprep maxiprep kit (QIAGEN) from 150 mL overnight bacterial culture supplemented with appropriate antibiotics. All plasmid sequences were confirmed by Sanger sequencing with appropriate sequencing primers (GATC Biotech). All the plasmids used in this study are listed in **Supplementary Table 9**, as well as DNA sequences. The construction of the combinatorial libraries for the directed evolution of nirFAST, as well as the screening protocol by FACS, are described in **Supplementary Methods**.

#### Cloning

The plasmids used in this study have been generated using isothermal Gibson Assembly or restriction enzymes cloning. The construction of the plasmids for the characterization of clones from the directed evolution experiments is described in **Supplementary Methods**.

Plasmid pAG1253 for bacterial expression of 6×His–TEVcs–nirFAST under the control of a T7 promoter was obtained by introducing R52K mutation in the sequence coding for nirFAST2.0 (see **Supplementary Methods**).

Plasmids for mammalian expression of emiRFP670 and miRFP713 under the control of CMV promoter were obtained from Addgene vectors #136556 and #136559. Plasmids pAG1329, pAG1334, pAG1335 and pAG1337 for mammalian expression of nirFAST-P2A-EGFP-cMyc, frFAST-P2A-EGFP-cMyc, emiRFP670-P2A-EGFP-cMyc and miRFP713-P2A-EGFP-cMyc respectively were obtained by replacement of the sequence of FAST by the sequences of nirFAST, frFAST, emiRFP670 and miRFP713 respectively in the plasmid pAG453 coding for FAST-P2A-EGFP (not published). The plasmid pAG1371 for mammalian expression of nirFAST under the control of a CMV promoter was obtained by replacing the sequence of frFAST by that of nirFAST in plasmid pAG504^10^ encoding frFAST. The plasmid pAG1372 for mammalian expression of H2B-nirFAST-cMyc was obtained by replacing the sequence of pFAST by nirFAST in the plasmid pAG657 for mammalian expression of H2B-pFAST^9^. The plasmid pAG1375 for mammalian expression of mito-nirFAST-cMyc was obtained by replacing the sequence of pFAST by that of nirFAST in the plasmid pAG671^9^ for the expression of mito-pFAST-cMyc. Similarly, the plasmid pAG1380 for mammalian expression of MAP4-nirFAST-cMyc was obtained by replacing the sequence of pFAST by that of nirFAST in plasmid pAG665^9^ for mammalian expression of MAP4-pFAST. The plasmid pAG1377 for mammalian expression of nirFAST-Cb5 was obtained by replacing the sequence of FAST_1-114_-eCFP by the sequence of nirFAST in the plasmid pAG1168 (not published) for mammalian expression of FAST_1-114_-ECFP-Cb5. Plasmids pAG1329 (nirFAST-P2A-EGFP-cMyc), pAG1334 (frFAST-P2A-EGFP-cMyc), pAG1335 (emiRFP670-P2A-EGFP-cMyc) and pAG1337 (miRFP713-P2A-EGFP-cMyc) described above were used for characterization of nirFAST properties at low magnification in chicken embryo tissues. For imaging of mito-nirFAST at high magnification in the chick neural tube, the CMV promotor in pAG1375 was replaced by the CAGGS promoter from pCAGGS-zH2B-frFAST^10^. The plasmid for expression of pCAGGS-zH2B-pFAST for expression of H2B-pFAST in chicken embryo was previously described^9^. The plasmid X-159 for expression of membrane-localized memb-mCherry in chicken embryo was previously described^33^. ^FAST^FUCCI (pSV1363) was obtained (via NEBuilder DNA Assembly) in two steps, first replacing the sequence of greenFAST by the sequence of nirFAST upstream zCdt1(1-190) in pSV1135 coding for redFAST-zGem(1-100)-P2A-greenFAST-zCdt1(1-190)^23^, then replacing in the resulting plasmid the sequence of redFAST by the sequence of pFAST upstream zGeminin(1-100). The plasmid pAG721 for the mammalian expression of lyn11-mCherry was obtained by Gibson assembly, by replacing the sequence of FAST by the sequence of mCherry in plasmid pAG106 for encoding lyn11-FAST and was used in a previous study^9^. The plasmid pAG1171 for encoding ^FIRE^mate-ECFP-giantin used in this study for the expression of ECFP at the Golgi apparatus was previously described^27^. The plasmid pAG324 used for encoding H2B-mCherry used in this study for the the expression of mCherry in the nucleus was previously described^10^. The plasmid pAG506 used for encoding mito-frFAST was previously described^10^.

The plasmid pAG1384 for mammalian expression of FKBP-nirFAST_115-125_-IRES-EGFP was obtained by replacing the sequence of iRFP670 by the sequence of EGFP in the plasmid pAG577^26^ for mammalian expression of FKBP-FAST_115-125_-IRES-iRFP670. The plasmid pAG1385 for mammalian expression of FRB-nirFAST1-_114-_-IRES-mTurquoise2 was obtained from the plasmid pAG490^26^ for mammalian expression of FRB-FAST_1-114_-IRES-mTurquoise2 by replacing the sequence of FAST_1-114_ by the sequence of nirFAST_1-114_. The plasmid pAG1501 for mammalian expression of TOM20-ECFP-nirFAST_1-114_ was obtained by replacing the sequence of FRB by the sequence of nirFAST_1-114_ in Addgene plasmid #171461 for mammalian expression of TOM20-ECFP-FRB(2021-2113 aa). The plasmid KIF17MD-flag-^nir-FIRE^mate was constructed by insertion of the ^nir-FIRE^mate from the plasmid pAG1501 encoding TOM20-ECFP-nirFAST_1-114_ in a vector for the expression of KIF17MD-flag-FIREmate (pFP5370)^27^.

#### In silico modeling of nirFAST

The homology models of nirFAST and frFAST were generated according to models previously described^34,35^. Briefly, sequence alignments between nirFAST and frFAST and the ultra-high resolution structure of the *Halorhodospira halophila* Photoactive Yellow Protein (PYP) as well as FAST 3D NMR structures (Protein Data Bank (PDB) ID: 6P4I, 7AV6, 7AVA, 7AVB^36,37^) were generated with Clustal W^38^. Alignments were manually refined. Three-dimensional nirFAST and frFAST models were built from these alignments and from crystallographic atomic coordinates of PYP using the automated comparative modeling tool MODELER (Sali and Blundell) implemented in Discovery Studio. The best model according to DOPE score (Discrete Optimized Protein Energy) and potential energy calculated by modeler were solvated with a minimum distance from periodic boundary of 10 Å (water and 0.145 M NaCl) and minimized using Adopted Basis NR algorithm to a final gradient of 0.001. The resulting structure were submitted to a 10 ns NAMD dynamics. Time course analysis has been performed by following RMSD/RMSF/electrostatic energy evolution.

Flexible ligand-rigid protein docking was performed using CDOCKER implemented in Discovery Studio 2020^39^. Random ligand conformations were generated from the initial ligand structure through high-temperature molecular dynamics. The best poses according to their ligscore2^40^ were retained and clustered according to their binding mode. The most significant poses were solvated and minimized using Adopted Basis NR algorithm to a final gradient of 0.001. The resulting structure were submitted to a 10 ns NAMD dynamics. Time course analysis has been performed by following RMSD/RMSF/electrostatic energy evolution. RMSD and RMSF are shown on **Supplementary Figure 6**. Coordinate files of the initial input and final output are provided as supplementary files.

#### Cell culture

HeLa cells (ATCC CRM-CCL2) were cultured in minimal essential medium supplemented with phenol red, Glutamax I, 1 mM of sodium pyruvate, 1% (vol/vol) of non-essential amino acids,10% (vol/vol) fetal bovine serum (FBS) and 1% (vol/vol) penicillin– streptomycin at 37 °C in a 5% CO_2_ atmosphere. HEK293T (ATCC CRL-3216) cells were cultured in Dulbecco’s modified Eagle medium (DMEM) supplemented with phenol red,10% (vol/vol) FBS and 1% (vol/vol) penicillin–streptomycin at 37 °C in a 5% CO_2_ atmosphere. For imaging, cells were seeded in μDish IBIDI (Biovalley) coated with poly-L-lysine. Cells were transiently transfected using Genejuice (Merck) according to the manufacturer’s protocols for 24 h before imaging. Cells were washed with Dulbecco’s PBS (DPBS) and treated with DMEM (without serum and phenol red) supplemented with the compounds at the indicated concentration.

#### Cell viability assay

Hela and HEK293T cells were treated with the indicated fluorogens concentrations for 24h. The cell viability was assessed using the Celltiter-Glo kit (Promega) through measurement of the luminescence signal on a Polar Star Optima plate-reader (BMG Labtech) according to manufacturer’s protocol.

#### Protein expression in bacteria and purification

Plasmids were transformed in BL21 (DE3) competent Escherichia coli (New England Biolabs) or Rosetta (DE3) pLys E. coli (Merck). Cells were grown at 37 °C in lysogen broth medium supplemented with 50 μg .mL^−1^ kanamycin (and 34 μg.mL^−1^ of chloramphenicol for Rosetta) to OD_600nm_ 0.6. Expression was induced overnight at 16 °C by adding isopropyl β-D-1-thiogalactopyranoside (IPTG) to a final concentration of 1 mM. Cells were collected by centrifugation (4,300 × g for 20 min at 4 °C) and frozen. For purification, the cell pellet was resuspended in lysis buffer (PBS supplemented with 2.5 mM MgCl_2_, 1 mM of protease inhibitor phenylmethanesulfonyl fluoride and 0.025 mg.mL^−1^ DNase, pH 7.4) and sonicated (5 min, 20% of amplitude) on ice. The lysate was incubated for 2 h on ice to allow DNA digestion by DNase. Cellular fragments were removed by centrifugation (9,000 × g for 1 h at 4 °C). The supernatant was incubated overnight at 4 °C by gentle agitation with pre-washed Ni-NTA agarose beads in PBS buffer complemented with 20 mM of imidazole. Beads were washed with ten volumes of PBS complemented with 20 mM of imidazole and with five volumes of PBS complemented with 40 mM of imidazole. His-tagged proteins were eluted with five volumes of PBS complemented with 0.5 M of imidazole. The buffer was exchanged to PBS (0.05 M phosphate buffer and 0.150 M NaCl) using PD-10 desalting columns or Midi-Trap G-25 (GE Healthcare). The purity of the proteins was evaluated using SDS–PAGE electrophoresis stained with Coomassie blue.

#### Physicochemical measurements

Steady-state UV-Vis and fluorescence spectra were recorded at 25 °C on a Spark 10 M (Tecan). Data were processed using GraphPad Prism v.10.0.3. Fluorescence quantum yield measurements were determined in 96-well plates using either FAST:HBR-3,5DOM or frFAST:HPAR-3OM as a reference. Solutions of 40 µM of proteins were used, in which the fluorogen was diluted to the right concentration, usually from 6 µM to 0.375 µM, allowing > 99% of complex formation. Absorption coefficients were determined directly by the previous experiments after determination of the optical path length.

Thermodynamic dissociation constants were determined by titration experiments in which we measured the fluorescence of the fluorescent assembly at various fluorogen concentrations using a Spark 10M plate reader (Tecan) and fitting data in Graphpad Prism 9 to a one-site specific binding model^9–14,23^ Detailed derivation of the thermodynamic dissociation constants can be found in ref 13 and the protocol of their measurement is detailed in ref 12. In brief, the dissociation constant of the fluorogen:protein assembly is determined by titration experiment in a 96-well plate format. As the fluorogen:protein assembly is strongly fluorescent, one can directly use fluorescence as a readout to determine the fraction of complex. Titrations are performed varying the fluorogen concentration while keeping the protein concentration constant. Measurement of the fluorescence intensity at each fluorogen concentration enables to quantify the fraction of complex. These experiments are done in large molar excess of fluorogen to be able to neglect the variation of free fluorogen concentration upon complex formation. We used protein concentrations of 35-50 nM. To determine the specific signal from the complex, the contribution of the free fluorogen (baseline) was measured and subtracted. We used twelve different concentrations of fluorogen. The fluorescence titration data were analyzed by fitting with a one-site specific binding model using Graphpad Prism 9 software.

#### Flow cytometry analysis

Flow cytometry analysis of HEK293T cells was performed on a MACSQuant analyzer equipped with 405 nm, 488 nm and 635 nm lasers and seven channels. To prepare samples, cells were first grown in cell culture flasks, then transiently co-transfected 24 h after seeding using Genejuice (Merck) according to the manufacturer’s protocol for 24 h. After 24 h, cells were centrifuged in PBS with BSA (1 mg.mL^−1^) and resuspend in PBS–BSA supplemented with the appropriate amounts of compounds. For each experiment, 20,000 cells positively expressing mTurquoise2 (Ex 405 nm / Em 425-475 nm) and EGFP (Ex 488 nm / Em 500-550 nm) or mCherry (Ex 488 nm / Em 565-605 nm) were analyzed with the following parameters: Ex 635 nm / Em 655-730 nm. Data were analyzed using FlowJo v.10.7.1. A figure presenting the gating strategy is presented in **Supplementary Figure 16**.

#### Fluorescence microscopy

The confocal micrographs of mammalian cells were acquired on an inverted Zeiss LSM 980 Laser Scanning Microscope equipped with a plan apochromat 63×/1.4 NA oil immersion objective, a plan-apochromat 40×/1.4 oil immersion objective and a plan-apochromat 20×/0.8 dry objective and an inverted Leica TCS SP5 confocal laser scanning microscope equipped with a 63×/1.4 NA oil immersion objective. The confocal micrographs of zebrafish were acquired on an upright Zeiss LSM 980 Laser scanning microscope equipped with a plan-apochromat 20×/1.0 DIC (UV)VIS-IR M27 75mm water objective. ZEN and Leica LAS AF softwares were used to collect the data. Icy and Fiji softwares were used to analyze the data. Fluorescence signal measurement was obtained through the Active Contour plugin on Icy. Lineage and tracking of dividing cells as well as corresponding fluorescence signal extraction was performed through Trackmate plugin^41,42^ (Fiji).

Live imaging in the chick embryonic neuroepithelium was performed on an inverted microscope (Nikon Ti Eclipse) equipped with a heating enclosure (DigitalPixel, UK), a spinning disk confocal head (Yokogawa CSUW1) with Borealis system (Andor) and a sCMOS Camera (Orca Flash4LT, Hamamatsu) driven by MicroManager software^43^. Image stacks were obtained at 3-min intervals either with a 10x objective (CFI Plan APO LBDA, NA 0.45, Nikon) with a 5 µm z-step or a 100× oil immersion objective (APO VC, NA 1.4, Nikon) with a 0.3 µm z-step.

Confocal spinning disk images for control of lysosomes positioning were acquired on a Nikon Inverted Eclipse Ti-E (Nikon) microscope equipped with a Spinning disk CSU-X1 (Yokogawa), a Kinetx 22 sCMOS camera (Photometrics) integrated in Metamorph software by Gataca Systems, using a 100× CFI apochromat VC 1.4 NA oil immersion objective (Nikon). The images were analyzed with Fiji (Image J).

#### Super-resolution

The comparative confocal and STED micrographs were acquired on the STEDYCON confocal and STED module (Abberior Instruments GmbH, Germany) connected to the lateral port of an IX83 inverted microscope (Evident Scientific). The STEDYCON SmartControl software was used for data acquisition. Imaging was performed using a UPLXAPO 100× oil immersion objective (Evident Scientific). The specimens were imaged with 561 nm (nirFAST600) or 640 nm (nirFAST680) lasers as well as a pulsed 775 nm depletion laser for STED imaging. Typically, images of 1024 × 1024 pixels were acquired with a pixel size in the range of 25–30 nm. Resolution assessment of confocal and STED images was performed using image decorrelation analysis plugin in Fiji^19^ using default parameters.

#### Zebrafish experiments

Adult zebrafish (*Danio rerio*) were kept at around 27-29°C on a 14 hr-light:10 hr-dark cycle and fed twice daily. Natural crosses obtained fertilized eggs which were raised at 28°C in Volvic water. Experiments were performed using the standard nacre strain. Developmental stages were determined and indicated as days post fertilization (dpf). The animal facility obtained permission from the French Ministry of Agriculture (agreement No. D-75-05-32.) and all animal procedures were performed in accordance with French animal welfare guidelines. HEK293T cells were seeded at 500,000 cells/mL concentration in 25 cm^2^ flasks and transfected after 24h with plasmid encoding nirFAST-P2A-EGFP using GeneJuice (Merck) according to manufacturer’s guidelines. After 24h, cells were harvested at 10,000,000 cells/mL concentration in serum free DMEM.

Cell suspension was loaded into a borosilicate glass needle pulled by a Flaming/Brown micropipette puller (Narishige, Japan, PN-30). 5∼10 nanoliters suspension were implanted into anesthetized (0.02% MS-222 tricaine (Sigma)) 2 dpf zebrafish larvae close to the heart by using an electronically regulated air-pressure microinjector (FemtoJet, Eppendorf). After injection, zebrafish larvae were placed in Volvic water and examined under a stereoscopic microscope for the presence of fluorescent cells and then raised for two more days at 28°C before imaging. For imaging, living zebrafish larvae were anesthetized in MS-222 tricaine solution and embedded in a lateral orientation in low-melting agarose (0.8%). Larval zebrafish were studied before the onset of sexual differentiation and their sex can therefore not be determined.

#### Chicken embryo experiments

JA57 chicken fertilized eggs were provided by EARL Morizeau (8 rue du Moulin, 28190 Dangers, France) and incubated at 38°C in a Sanyo MIR-253 incubator. Embryos used in this study were between E2 (HH14) and E3 (HH14 + 24 h). The sex of the embryos was not determined. Under current European Union regulations, experiments on avian embryos between 2 and 4 days *in ovo* are not subject to restrictions.

*In ovo* electroporation in the chick neural tube was performed at embryonic day 2 (E2, HH stage 14), by applying five pulses of 50 ms at 25V with 100 ms in between, using a square-wave electroporator (Nepa Gene, CUY21SC) and a pair of 5 mm gold-plated electrodes (BTW Genetrode model 512) separated by a 4 mm interval. The DNA solution was injected directly into the lumen of the neural tube via glass capillaries. Bilateral electroporation was achieved by switching the electrodes polarity and repeating the procedure after 3h. DNA constructs were used at 1 µg/µl, except pCX-mbCherry which was used at 0.3 µg/µl; and pCX-zH2B-pFAST and pCX-mito-nirFAST which were used at 0.5 µg/µl. En-face culture of the embryonic neuroepithelium was performed at E3 (24 h after electroporation). After extraction from the egg and removal of extraembryonic membranes in PBS, embryos were transferred to 37 °C F12 medium and pinned down with dissection needles at the level of the hindbrain in a 35 mm Sylgard dissection dish. A dissection needle was used to slit the roof plate and separate the neural tube from the somites from hindbrain to caudal end on both sides of the embryo. The neural tube and notochord were then transferred in a drop of F12 medium to a glass-bottom culture dish (MatTek, P35G-0-14-C) and medium was replaced with a 500µl-1ml of 1% low melting point agarose/F12 medium (maintained at 38°C). Excess medium was removed so that the neural tube would flatten with its apical surface facing the bottom of the dish, in an inverted open book conformation. After 30s of polymerization on ice, an extra layer of agarose medium (200 µl) was added to cover the whole tissue and left to harden. 1.8 mL of 38 °C culture medium was added (F12/1mM Sodium pyruvate) and the culture dish was transferred to the 38 °C chamber of a spinning disk confocal microscope. To image nirFAST, intermediate dilutions of ligands HPAR-3OM and HPAR-3,5DOM were prepared by diluting the original 20 mM stocks in F12 medium, and appropriate volumes were added to the dish to reach the desired final concentration.

## ACKNOWLEDGMENTS

We thank K. D. Wittrup, for providing us with the pCTCON2 vector and the EBY100 yeast strain for the yeast display selection. We thank V. Verkhusha for the plasmids for mammalian expression of emiRFP670 (Addgene 136556) and miRFP713 (Addgene 136559). We thank T. Miyamoto for the plasmid for mammalian expression of TOM20-ECFP-FRB (Addgene #171461). We thank the flow cytometry facility CISA (Cytométrie Imagerie Saint-Antoine) of UMS LUMIC at the Faculty of Medicine of Sorbonne Université, and, more particularly, Annie Munier and Angélique Vinit for their assistance. We thank the microscopy facility of the Institut de Biologie Paris Seine of Sorbonne University, and more particularly France Lam and Chloé Chaumeton for their assistance. We thank Chloé Chaumeton from the microscopy facility of the Institut de Biologie Paris Seine of Sorbonne University, and Frédéric Eghiaian from Aberrior Instruments for their assistance for the STED imaging. This work has been supported by the European Research Council (ERC-2016-CoG-724705 FLUOSWITCH), the Agence Nationale de la Recherche (ANR-23-CE44-0014-01CATCHFIRE), the Institut Universitaire de France. LEH thanks the École Normale Supérieure and the Fondation pour la Recherche Médicale (grant FDT202304016615 to L.E.H.) for PhD funding.

## AUTHOR CONTRIBUTIONS

L.E.H and A.G. designed the overall project and wrote the paper with the help of the other authors. L.E.H, B.B., O.J., C.R., M.V., E.F., N.P., F.P., S.V., X.M. and A.G. designed the experiments. L.E.H, B.B., O.J., C.L., A.G.T., C.R., M.V., E.F., N.P., S.V. performed the experiments. L.E.H, B.B., O.J., C.R., M.V., E.F., N.P., F.P., S.V., X.M. and A.G. analyzed the experiments.

## COMPETING INTERESTS

The authors declare the following competing financial interest: A.G. and F.P. are co-founders and hold equity in Twinkle Bioscience/The Twinkle Factory, a company commercializing the FAST, split-FAST and CATCHFIRE technologies. The other authors declare no competing interests.

### DATA AVAILABILITY

The data supporting the findings of this study are available within the article and supplementary information, and are available from the corresponding authors upon reasonable request. The plasmids developed in this study will be available from Addgene.

