## Supplementary information for "A tunable and versatile chemogenetic near infrared fluorescent reporter"

**This PDF file includes:**

Supplementary legends of Supplementary movies

Supplementary Text 1

Supplementary Figures 1-16

Supplementary Tables 1-9

Supplementary Methods

**Legends of Supplementary Movies**

**Supplementary Movie 1.** Labeling kinetics of nirFAST680 and nirFAST715. HeLa cells expressing nirFAST-P2A-EGFP were imaged at high imaging rate (1 frame / 2s) by confocal microscopy prior and after addition of 10 µM of HPAR-3OM or HPAR-3,5DOM to assemble nirFAST680 or nirFAST715. Scale bar, 10 µm.

**Supplementary Movie 2. Monitoring cellular division in chicken embryo tissues using nirFAST.** Plasmids encoding H2B-pFAST (cyan), memb-mCherry (yellow) and mito-nirFAST (magenta) were electroporated in the neural tube *in ovo*, at embryonic day 2 (E2, HH stage 13–14). 24 h later, embryos were dissected, and the neuroepithelium was imaged in en-face view in presence of 1 µM HMBR (to assemble pFAST540) and 10 µM HPAR3,5DOM (to assemble nirFAST715) using a spinning disk confocal microscope. Scale bar, 10 µm.

**Supplementary Movie 3. Long-term timelapse imaging of $^{\text{FAST}}$FUCCI-expressing cells.** HEK293T cells transfected with $^{\text{FAST}}$FUCCI are imaged over a large field of view for over 24h after labeling with 1 µM HMBR and 10 µM HPAR-3,5DOM**.** Scale bar, 20 µm.

**Supplementary Movie 4. Timelapse imaging of dividing $^{\text{FAST}}$FUCCI-expressing cells.** HEK293T cells transfected with $^{\text{FAST}}$FUCCI are imaged for over 24h after labeling with 1 µM HMBR and 10 µM HPAR-3,5DOM (close-up view)**.** Scale bar, 20 µm

**Supplementary Movie 5. Kinetics of $^{\text{nir}}$CATCHFIRE680.** HeLa cells co-expressing EGFP-$^{\text{FIRE}}$tag and Tom20-ECFP-$^{\text{nir-FIRE}}$mate were treated with 10 µM of HPAR-3OM and imaged using timelapse confocal microscopy over 10 minutes. Scale bar, 10 µm.

**Supplementary Movie 6. Kinetics of $^{\text{nir}}$CATCHFIRE715.** HeLa cells co-expressing EGFP-$^{\text{FIRE}}$tag and Tom20-ECFP-$^{\text{nirFIRE}}$mate were treated with 10 µM of HPAR-3,5DOM and imaged using timelapse confocal microscopy over 10 minutes. Scale bar, 10 µm.

**Supplementary Movie 7. Control and tracking of lysosome positioning.** HeLa cells coexpressing LAMP1–mCherry–$^{\text{FIRE}}$tag and $^{\text{nir-FIRE}}$mate–KIF17 were imaged by spinning-disk microscopy for 30 min, then HPAR-3,5DOM was added. Scale bar, 10 µm.

### Supplementary Text 1: Engineering of nirFAST

#### The FAST family of chemogenetic reporters

nirFAST was engineered from the chemogenetic fluorescent reporter far-red (fr)FAST[1], which belongs to the fluorescence-activating and absorption-shifting tag (FAST) family. The FAST family is a set of small proteins that can be used as fluorescent markers for monitoring gene expression and protein localization in live cells and organisms. Evolved from the 14-kDa photoactive yellow protein (PYP) from the phototactic bacterium *Halorhodospira halophila*, prototypical FAST binds and stabilizes the fluorescent state of hydroxybenzylidene rhodanine (HBR) derivatives[2,3], push-pull chromophores composed of an electron-donating phenol conjugated to an electron-withdrawing rhodanine head. These chromophores are barely fluorescent when free because they dissipate light energy through ultrafast non-radiative de-excitation pathways: rotations around the methylene bridge bonds have been proposed to allow internal conversion through conical intersection as described for the 4-hydroxybenzylidene-imidazolinone (HBI) chromophore found in the green fluorescent protein (GFP)[4] . HBR derivatives however strongly fluoresce visible light when embedded within FAST because they adopt a quasi-planar conformation with an orientation of the phenolic ring that allows the glutamic acid 46 to stabilize the anionic phenolate state[5,6]. This mode of binding leads to high fluorescence quantum yield and a red-shift in absorption that participates to the high fluorogenicity of the system. The spectral properties of FAST could be tuned by a concerted strategy involving chromophore engineering and directed protein evolution[1,5]. This approach allowed the generation of frFAST, a FAST variant able to bind and stabilize the fluorescent state of HPAR-3OM, yielding a fluorescent assembly, hereafter named frFAST670, with a maximal absorption at $\lambda_{abs}$ = 555 nm and a maximal emission at $\lambda_{em}$ = 670 nm.

#### Spectral tuning by rational design

One main advantage of chemogenetic reporters is the ability to perform spectral tuning by either molecular engineering of the fluorogen or protein engineering of the tag. To engineer nirFAST, we first added a second methoxy group in ortho position of the phenol ring of HPAR-3OM, giving a new fluorogen called HPAR-3,5DOM. The introduction of two methoxy groups in ortho position of the phenol in HBR derivatives was previously shown to significantly red-shift both the absorption and emission wavelengths of FAST:fluorogen assemblies[3,7]. Similarly, we observed that the presence of two methoxy groups in ortho position of the phenol in HPAR-3,5DOM red-shifted both absorption and emission wavelengths, $\lambda_{abs}$ = 602 nm and $\lambda_{em}$ = 713 nm, when bound to frFAST (**Supplementary Table 1**). However, the introduction of the second methoxy group led to a 5-fold reduction of fluorogen binding affinity and to a 2.5-fold decrease of the molar absorption coefficient at $\lambda_{abs}$ = 602 nm because of a partial protonation of the fluorogen (as evidenced by the presence of a second absorption band at 430 nm, characteristic of the protonated phenol state of the fluorogen). To further shift the absorption to the red, we mutated the glutamic acid 46 to glutamine. The mutation E46Q was previously shown to red shift the absorption of holo-PYP (in complex with its natural hydroxycinnamoyl prosthetic group)[8]: the reduced hydrogen-bond donating ability of glutamine versus glutamic acid was proposed to increase the electron-donating ability of the anionic phenolate, and consequently to red-shift absorption. Similar behavior was previously observed in FAST variants bearing this mutation[7]. Likewise, in the present work, we observed that introduction of E46Q in frFAST led to 40-50 nm red-shifted absorption of the anionic phenolate of HPAR-3OM and HPAR-3,5DOM (**Supplementary Tables 1** and **2**). However, the E46Q mutation reduced the absorption band of the anionic phenolate and increased the absorption band of the protonated phenol at shorter wavelengths (**Supplementary Tables 1** and **2**, **Supplementary Figure 2**). We assume that the E46Q mutation changes the immediate electrostatic surrounding of the embedded chromophore, changing thus its apparent p$K_A$ and thus its protonation state. We observed similar behavior by introducing E46Q in the variant frFAST$^{I107V}$, a frFAST variant bearing the mutation I107V previously shown to behave as well as frFAST.

**Directed evolution**

In order to improve the affinity for the fluorogen as well as the molecular brightness we used directed evolution. We first used both frFAST$^{E46Q}$ and frFAST$^{E46Q,I107V}$ as starting templates for the construction of a test combinatorial library of variants. After optimization of the conditions, we eventually constructed a final combinatorial library only based on frFAST$^{E46Q,I107V}$ (hereafter called **nirFAST0.1**). This library was constructed by error-prone PCR, and then cloned in a yeast display plasmid. Initial transformation in bacteria to maximize the number of variants in the library was followed by large-scale transformation in yeast cells. The final library in yeast contained $10^6$ different clones. The yeast-displayed library was then screened by fluorescence-activated cell sorting (FACS) using a 640 nm excitation laser to sort variants forming bright NIR fluorescent assemblies with HPAR-3,5DOM. The concentration of HPAR-3,5DOM was decreased during the course of FACS screening, from 10 μM to 5 μM, as a way to increase the pressure of selection and to sort tighter binders. After five rounds of enrichment, we isolated and sequenced 48 different clones, and characterized their fluorescence properties with HPAR-3,5DOM in yeast. Out of the 48 clones sent to sequencing, 21 different sequences were identified with an average of 3 amino acid mutations per sequence, and 8 of these sequences had 2 or more occurrences. All of the clones displayed improved fluorescence compared to the starting clone.

We then chose the five best clones for recombinant expression in bacteria and characterized their spectral properties and binding affinity with HPAR-3OM and HPAR-

3,5DOM. All five different clones displayed improved brightness and enhanced affinities with both fluorogens in comparison with nirFAST0.1 (**Supplementary Table 1**). The best selected clone, called **nirFAST1.0** (with the mutations I31V, E46Q, D48G, T70S, I107V with respect to frFAST), was further improved by introduction of the mutations N43S and K78I found in other selected clones. The resulting protein, called **nirFAST1.1**, efficiently binds HPAR-3,5DOM ($K_D$ = 0.31 $\mu$M) and forms an assembly 10-fold brighter than nirFAST0.1:HPAR-3,5DOM (**Supplementary Tables 1** and **2**). The complex nirFAST1.1:HPAR-3,5DOM is characterized by absorption/emission peaks at $\lambda_{abs}$ = 630 nm and $\lambda_{em}$ = 715 nm, a fluorescence quantum yield FQY = 6.7 % and a molar absorption coefficient $\varepsilon_{630\ nm}$ = 34,000 $M^{-1}.cm^{-1}$. Within nirFAST1.1, embedded HPAR-3,5DOM is mainly deprotonated: the $\varepsilon_{deprotonated}/\varepsilon_{protonated}$ ratio is 2 (**Supplementary Figure 2**). This gain in molar absorption coefficient explains the 10-fold brightness increase.

To further increase the absorbance at 630 nm, we constructed a second combinatorial library by random mutagenesis of nirFAST1.1. This second library contained 3.9 $\times$ $10^6$ different clones. Five rounds of FACS screening, during which the fluorogen concentration was decreased from 2.5 to 1 μM, allowed us to enrich the yeast library in improved variants. After the fifth round, we analysed the fluorescence properties of 132 individual clones in yeast with HPAR-3,5DOM. We then expressed in bacteria and purified the five best variants and characterized their spectral properties and affinity with HPAR-3OM and HPAR-3,5DOM (**Supplementary Tables 3** and **4**). The best variant of this second selection, named **nirFAST2.0** (with the mutations I31V, D36N, Q41K, N43S, E46Q, D48G, T70S, K78I, I107V with respect to frFAST), was further subjected to site-directed mutagenesis using the mutations G21R, G25R, K41R, G47R, R52K, M95T, V107I and S117R found in other selected clones to further improve its properties. Eight single variants of nirFAST2.0 were generated and characterized after expression in bacteria. All eight variants displayed enhanced affinity and enhanced brightness with HPAR-3OM and HPAR-3,5DOM in comparison with nirFAST1.1 (**Supplementary Tables 3** and **4**).

**Evaluation of the best variants in cultured mammalian cells**

The brightness of the six best performing variants was further evaluated in cultured mammalian cells to identify the best one. For this purpose, the variants were co-expressed stoichiometrically with EGFP using a polycistronic expression cassette containing a viral P2A sequence for ribosomal skipping during translation, allowing us to normalize the NIR fluorescent signal by the protein expression level using the EGFP signal. All variants were characterized with HPAR-3OM and HPAR-3,5DOM and the ratio of NIR fluorescence and green fluorescence was evaluated and compared (**Supplementary Figure 5**). From this screening, B.R5-24.5 (dubbed **nirFAST**) appeared to be the brightest variant with HPAR-3OM and among the brightest with HPAR-3,5DOM. In addition, **nirFAST** displayed narrower absorption and emission bands with HPAR-3OM and HPAR-3,5DOM in comparison with the other mutants. Finally, **nirFAST** was the variant with the best combination of molecular brightness and binding affinities for HPAR-3OM and HPAR-3,5DOM. Taking into account both the observations in vitro and in cells, **nirFAST** appeared to be the variant with the best properties for imaging in the NIR region. This variant binds HPAR-3,5DOM with an improved affinity ($K_D$ = 36 nM) and displays spectral properties in the NIR region ($\lambda_{abs}$ = 636 nm and $\lambda_{em}$ = 715 nm). With HPAR-3,5DOM, only one peak of absorption was observed (**Supplementary Figure 2**), corresponding to a full binding and stabilization of the deprotonated state of the fluorogen, which is emissive.

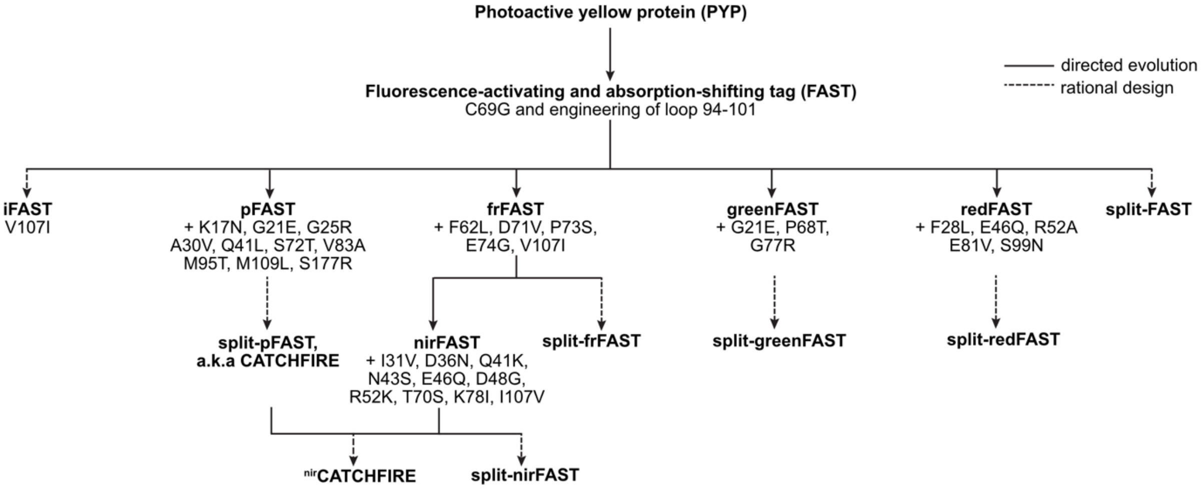


**Supplementary Figure 1. Family tree of the FAST variants.** Engineering efforts have helped expand the family of available FAST variants for various imaging applications. iFAST[9] is a variant with enhanced brightness compared to FAST, while pFAST[5] is a promiscuous variant able to form fluorescent assemblies spanning the visible spectrum, with compatibility with stimulated emission depletion (STED) nanoscopy. greenFAST and redFAST[7] are two orthogonal fluorogen binders enabling multicolor imaging. frFAST[1] bears advantageous spectral properties in the far-red region, and has been used as a starting point to develop nirFAST (this work). Some of these reporters have been split between positions 114-115, enabling the development of protein-protein interaction reporters[1,7,10,11] in addition to protein proximity inducers[12].

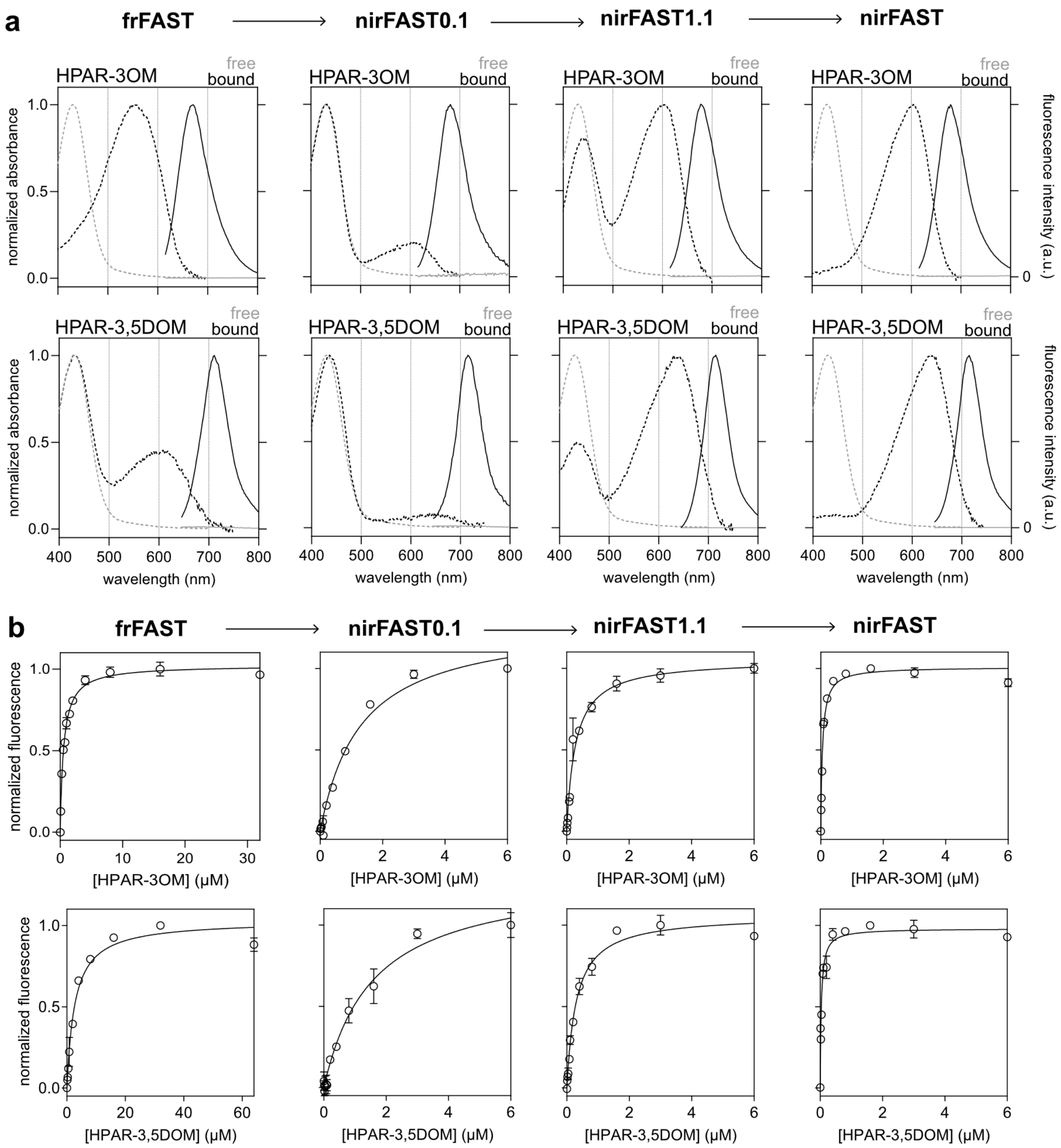


**Supplementary Figure 2. Physicochemical characterisation of the main variants.** (**a**) Absorption (dashed line, left axis) and emission (solid line, right axis) spectra of HPAR-3OM and HPAR-3,5DOM free (grey) or bound (black) to frFAST, nirFAST0.1, nirFAST1.1. and nirFAST. Spectra were recorded in pH 7.4 PBS at 25 °C. Concentrations of fluorogens and protein (here in excess) were chosen to achieve > 95% of fluorescent assembly (**b**) Affinity of frFAST, nirFAST0.1, nirFAST1.1. and nirFAST for HPAR-3OM and HPAR-3,5DOM. Graphs show the normalized fluorescence of bound protein at equilibrium for various fluorogen concentrations. The protein concentration was 50-100 nM. The titration experiments were performed in pH 7.4 PBS at 25 °C**.** Data represent the mean ± SEM (n = 3). Least spares fit (line) gave the dissociation constant $K_D$ of each assembly.

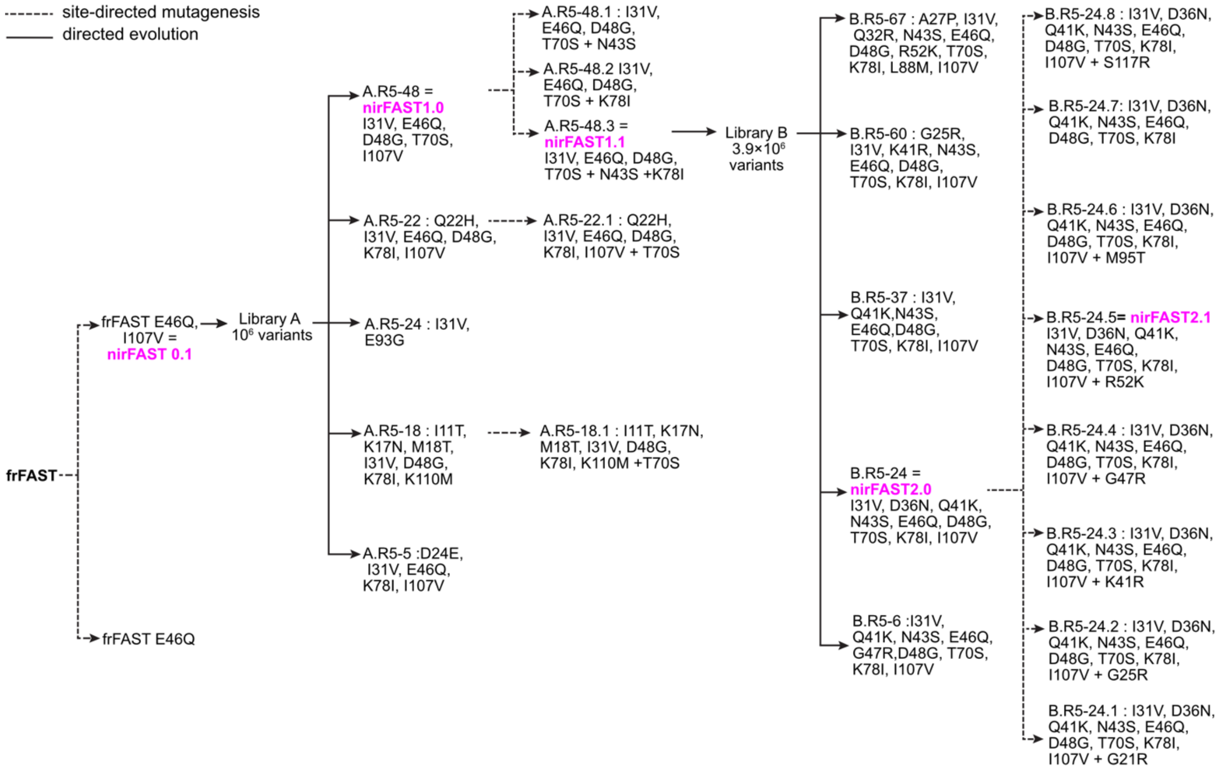


**Supplementary Figure 3. Evolution tree.** This tree presents the different variants generated during this study.

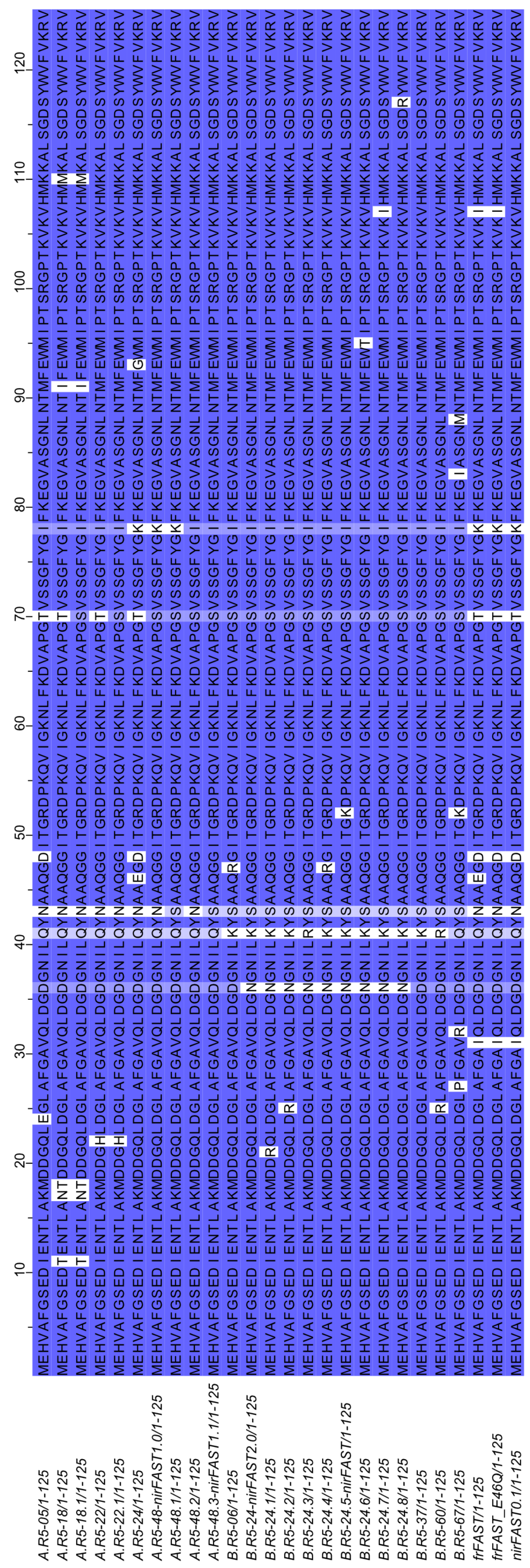

**Supplementary Figure 4. Alignment of the different variants generated during this study.**

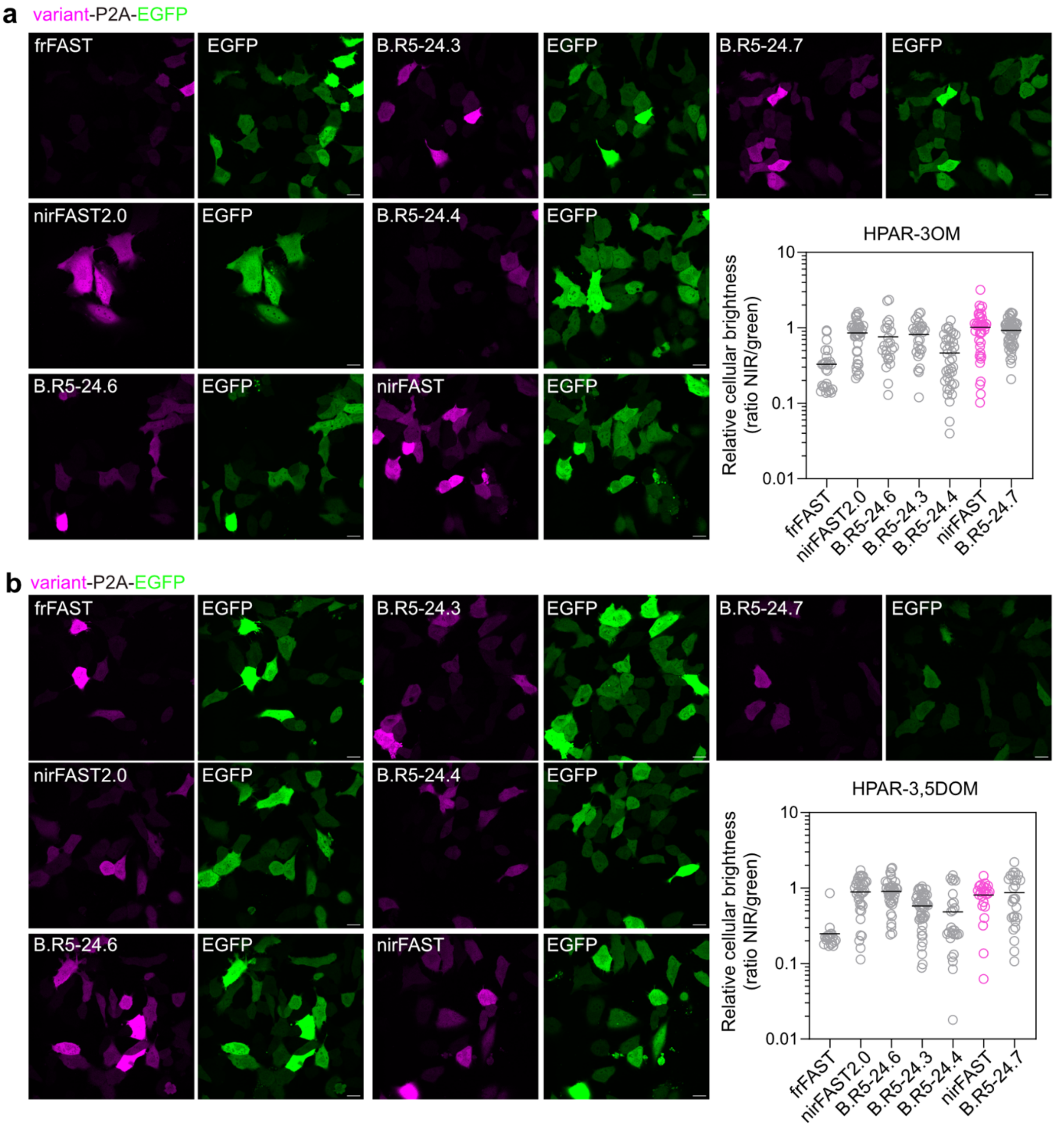


**Supplementary Figure 5. Comparison of the cellular brightness of the selected variants.** Confocal micrographs of HeLa cells expressing variant-P2A-EGFP labeled with 10 μM HPAR-3OM (**a**) or 10 μM HPAR-3,5DOM (**b**). Scale bars, 20 μm. Representative micrographs of n > 18 cells. The graphs show the relative cellular brightness computed by normalizing the fluorescence of the NIR reporters with the green fluorescence of the stoichiometrically expressed EGFP (ratio NIR/green). The black line indicates the mean value. See **Supplementary Table 8** for detailed imaging settings.

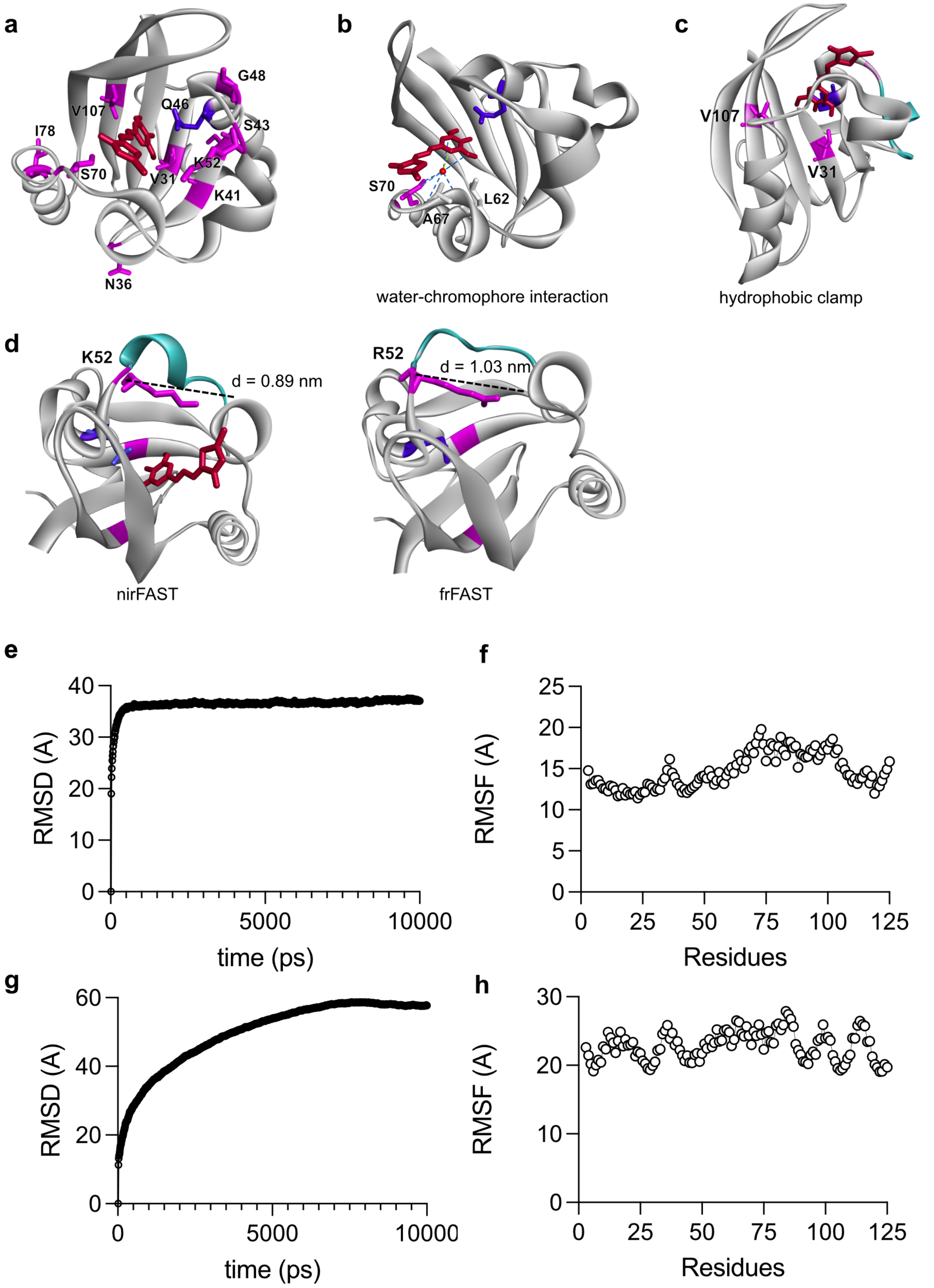


i

| | box dimension (LxWxH) | total number of atoms | total number of water molecules | NaCl concentration |
|---|---|---|---|---|
| nirFAST: HPAR-3,5DOM | 53x52x46 | 14880 | 4324 | 0.145 M |
| frFAST | 75x82x85 | 52673 | 15153 | 0.145 M |

**Supplementary Figure 6. Structural model of nirFAST715**. The model was generated by homology modeling and molecular dynamics using the crystal structure of the *Halorhodospira halophila* Photoactive Yellow Protein (PYP) (PDB: 6P4I). HPAR-3,5DOM is shown in red. Gln 46 is in blue. (**a**) Residues introduced during the engineering process are shown in magenta. (**b**) View showing the interaction of the structural water molecule and the chromophore. The residue Ser70 (from the mutation T70S) establishes hydrogen-bonds with this structural water molecule, stabilizing the overall assembly. (**c**) The mutations I31V and I107V widen the bottom of the binding pocket and create a hydrophobic clamp that interacts with the methyl of the methoxy groups of HPAR-3,5DOM. This allows a deeper positioning of the phenolate into the binding cavity, favouring the stabilization of the phenolate by Gln46. (**d**) The mutation R52K enables to reduce the size of binding pocket by allowing the residues 53 to 58 to adopt an $\alpha$-helix conformation. On the left is shown the model of nirFAST containing Lys52 and on the right the model of frFAST containing Arg52. The shown distances (*d*) are in between the $\alpha$ carbon of residue 52 and the $\alpha$ carbon of Val66. (**e**) Root Mean Square Displacement (RMSD) of nirFAST715 (nirFAST bound to HPAR-3,5DOM) during 10 ns molecular dynamic simulations. (**f**) Root Mean Square Fluctuation (RMSF) of the residues within nirFAST715 (nirFAST bound to HPAR-3,5DOM) during 10 ns molecular dynamic simulation. (**g**) Root Mean Square Displacement (RMSD) of frFAST during 10 ns molecular dynamic simulation. (**h**) Root Mean Square Fluctuation (RMSF) of the residues within frFAST during 10 ns molecular dynamic simulations. (**i**) Simulation set-up summary.

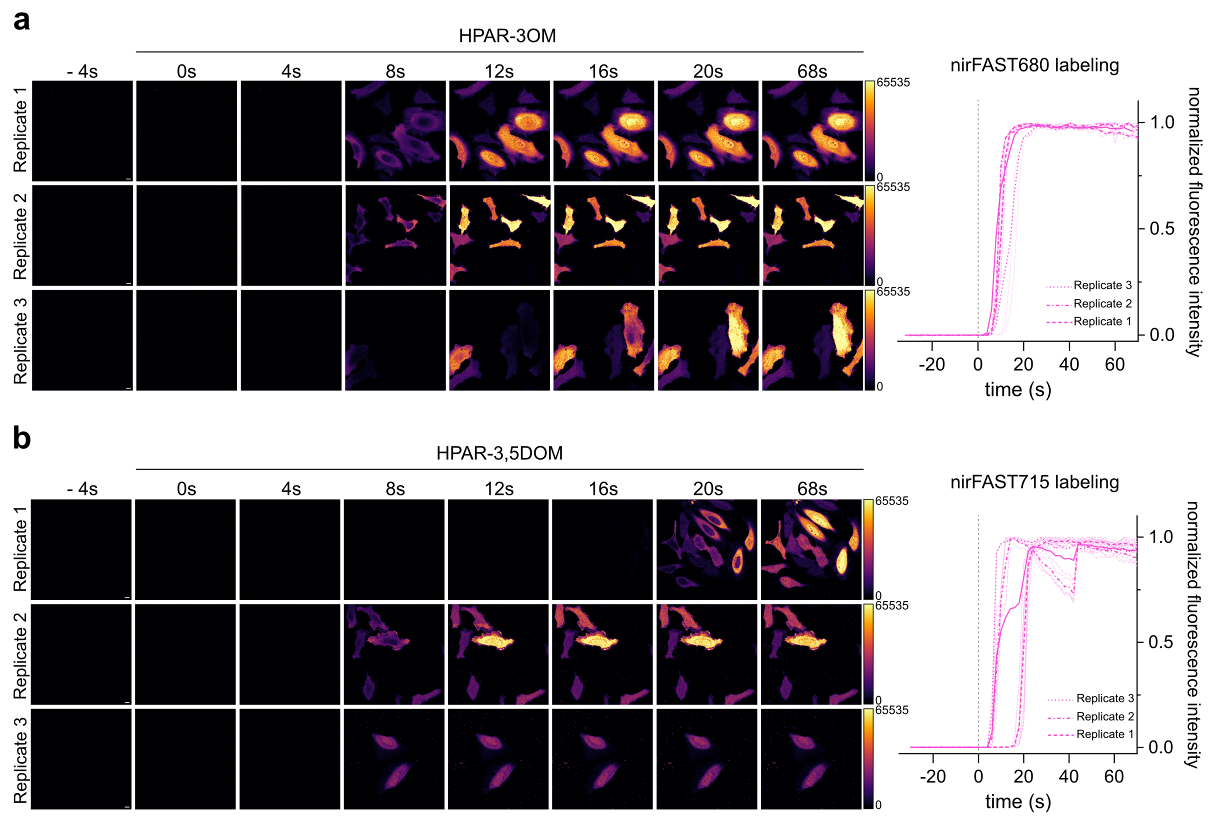


**Supplementary Figure 7. nirFAST labeling kinetics with HPAR-3OM and HPAR-3,5DOM.** Live HeLa cells expressing nirFAST-P2A-EGFP were labeled with 10 µM HPAR-3OM (to assemble nirFAST680) **(a)** or HPAR-3,5DOM (to assemble nirFAST715) **(b)** at t = 0 s (grey dashed line) and imaged every 2 seconds with 639 nm excitation. Micrographs of three independent experiments are shown, in addition to NIR fluorescence evolution over time of n = 11 cells for nirFAST680 and n = 16 cells for nirFAST715 (see also **Supplementary Movie 1**). Note that the loss of NIR signal observed for replicate 2 of nirFAST715 is due to a loss of focus. Scale bars 10 µm.

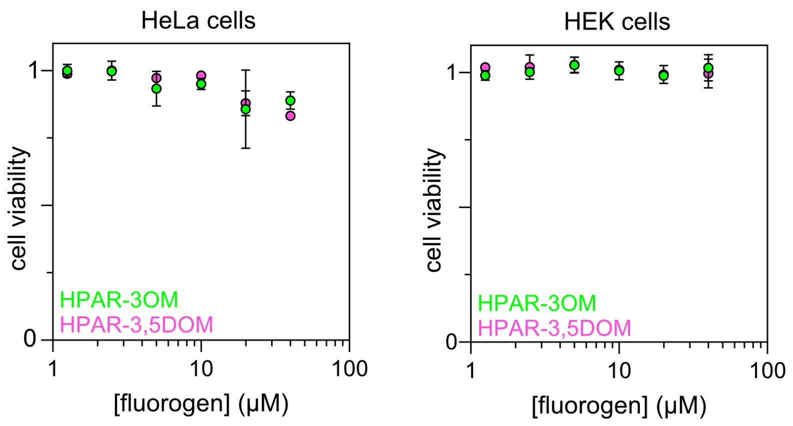


**Supplementary Figure 8. Evaluation of the effect of HPAR-3OM and HPAR-3,5DOM on cell viability.** Cell viability of HeLa cells (5,000 cells per well) and HEK293T cells (40,000 cells per well) treated with various concentrations (1.25, 2.5, 5, 10, 20, 40 µM) of HPAR-3OM or HPAR-3,5DOM for 24 h. Data represent the mean ± SD of n = 2 technical replicates for HeLa cells and n = 2 independent experiments for HEK293T cells.

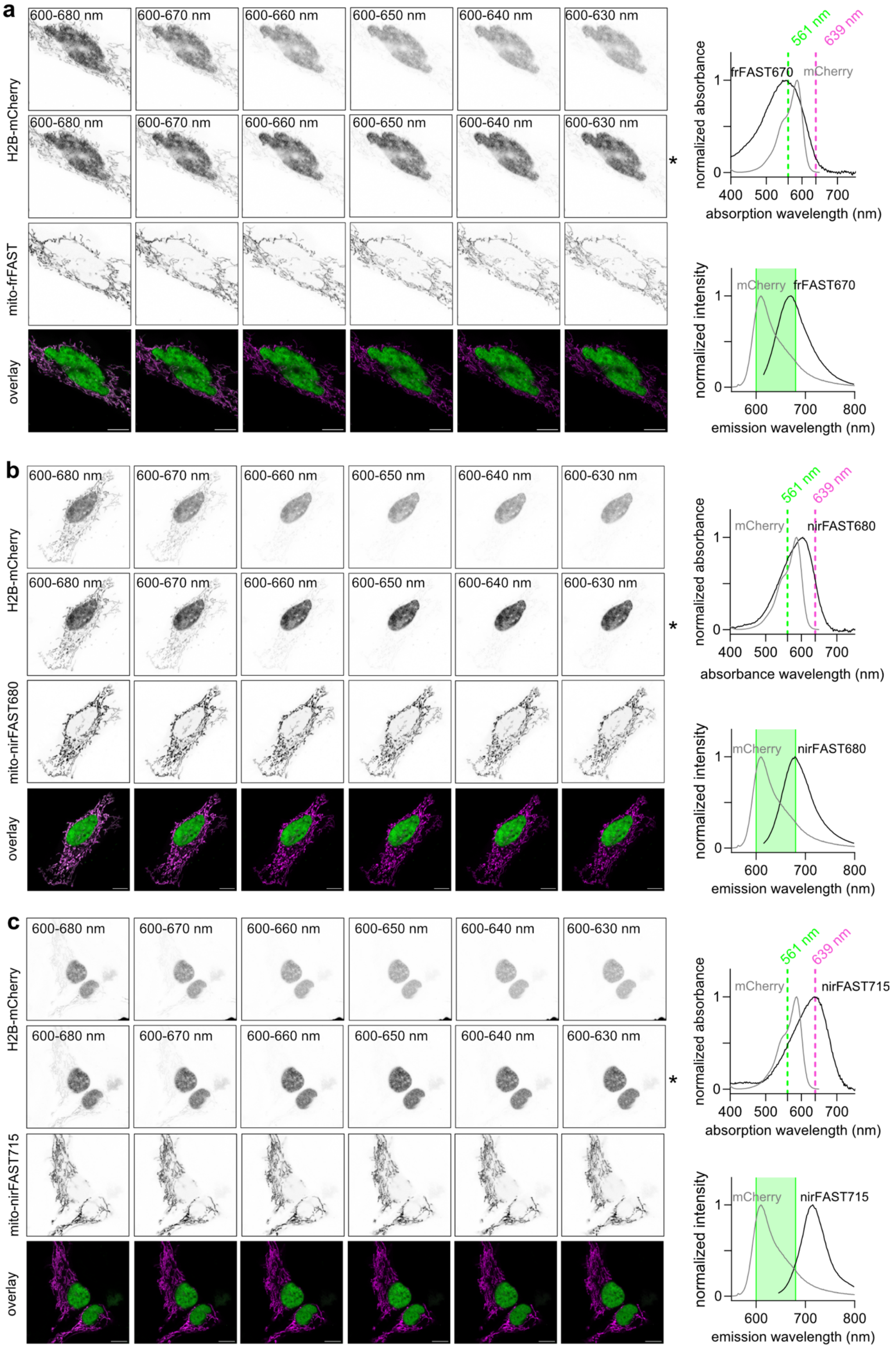
a
600-680 nm
600-670 nm
600-660 nm
600-650 nm
600-640 nm
600-630 nm
H2B-mCherry
mito-frFAST
overlay
*
561 nm
639 nm
normalized absorbance
frFAST670
mCherry
absorption wavelength (nm)
normalized intensity
emission wavelength (nm)
b
mito-nirFAST680
nirFAST680
absorbance wavelength (nm)
c
mito-nirFAST715
nirFAST715

**Supplementary Figure 9. Two-color imaging of frFAST670, nirFAST680 and nirFAST715 together with the red fluorescent protein mCherry.** Micrographs of HeLa cells co-expressing H2B-mCherry and mitochondria-targeted frFAST (**a**), nirFAST (**b**-**c**). Cells were treated with 10 μM HPAR-3OM to assemble frFAST670 (**a**) and nirFAST680 (**b**), and 10 μM HPAR-3,5DOM to assemble nirFAST715 (**c**). mCherry was imaged exciting at 561 nm varying the detection window from 600-680 nm to 600-630 nm, while keeping the detector settings identical. The first row of micrographs shows the raw images, while the second row (indicated with a star) shows images with adjusted levels for obtaining identical mCherry nuclear signal. The NIR reporters were imaged exciting at 639 nm with a 650-790 nm detection window. The overlay was obtained by merging the NIR image with the adjusted mCherry image. Representative micrographs from two independent experiments (n > 5 cells). On the right-hand side, are shown the absorption and emission spectra of each pair of reporters (mCherry spectra were obtained from FPbase.org). On the absorption spectra, are indicated the excitation lasers used for imaging each reporter. On the emission spectra, is indicated the 600-680 nm detection window used to detect mCherry. See **Supplementary Table 8** for detailed imaging settings. Scale bars, 10 μm

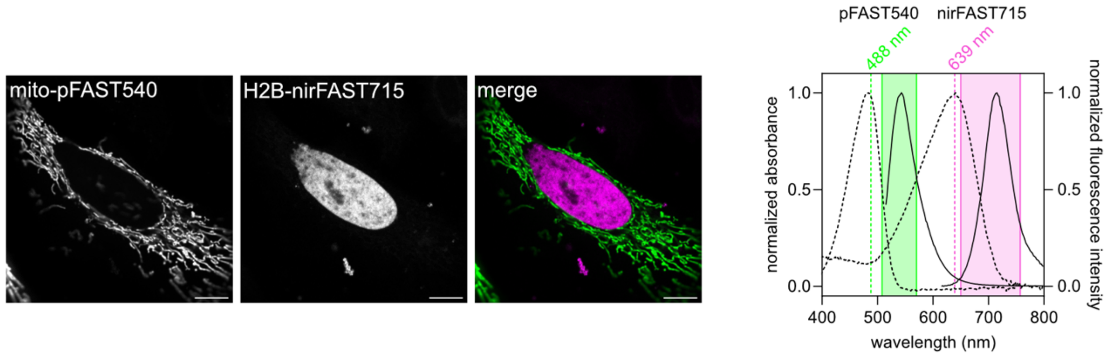


**Supplementary Figure 10. Two-color imaging of pFAST540 and nirFAST715.** Confocal micrographs of live HeLa cells co-expressing pFAST fused to a mitochrondia targeting sequence (mito) and nirFAST fused to H2B. Cells were labeled with 1 µM HMBR to assemble pFAST540 and 10 µM HPAR-3,5DOM to assemble nirFAST715. Representative micrographs from three independent experiments (n > 6 cells). Scale bars, 10 µm. The graph shows the imaging settings and spectral properties of the reporters used. Absorption (dashed lines) and emission spectra (solid lines) of pFAST540 and nirFAST715 are shown. The two excitation wavelengths used are shown on the absorption spectra graph and the two spectral windows are indicated on the emission spectra graph as colored areas. See **Supplementary Table 8** for detailed imaging settings.

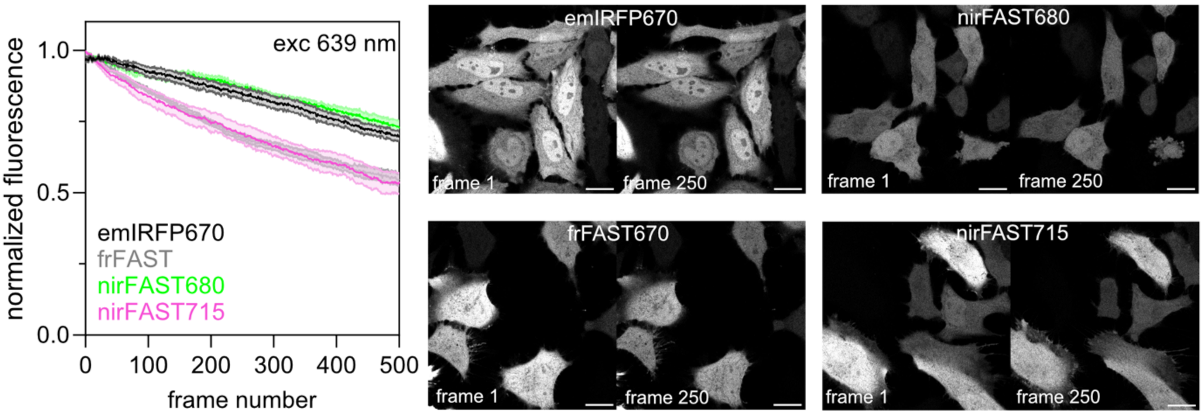


**Supplementary Figure 11. Photostability analysis.** Fluorescence levels of emiRFP670, frFAST670, nirFAST680 and nirFAST715 in live HeLa cells upon long-term observation with 639 nm laser excitation. Cells were labeled with 10 µM HPAR-3OM to assemble frFAST670 and nirFAST715, and 10 µM HPAR-3,5DOM to assemble nirFAST715. The graph shows the average fluorescence intensity of n = 7 (emiRFP670), 8 (frFAST670), 6 (nirFAST680), 6 (nirFAST715) cells from two independent experiments. On the right, are shown representative micrographs at different time points. Scale bars, 20 µm. See **Supplementary Table 8** for detailed acquisition parameters.

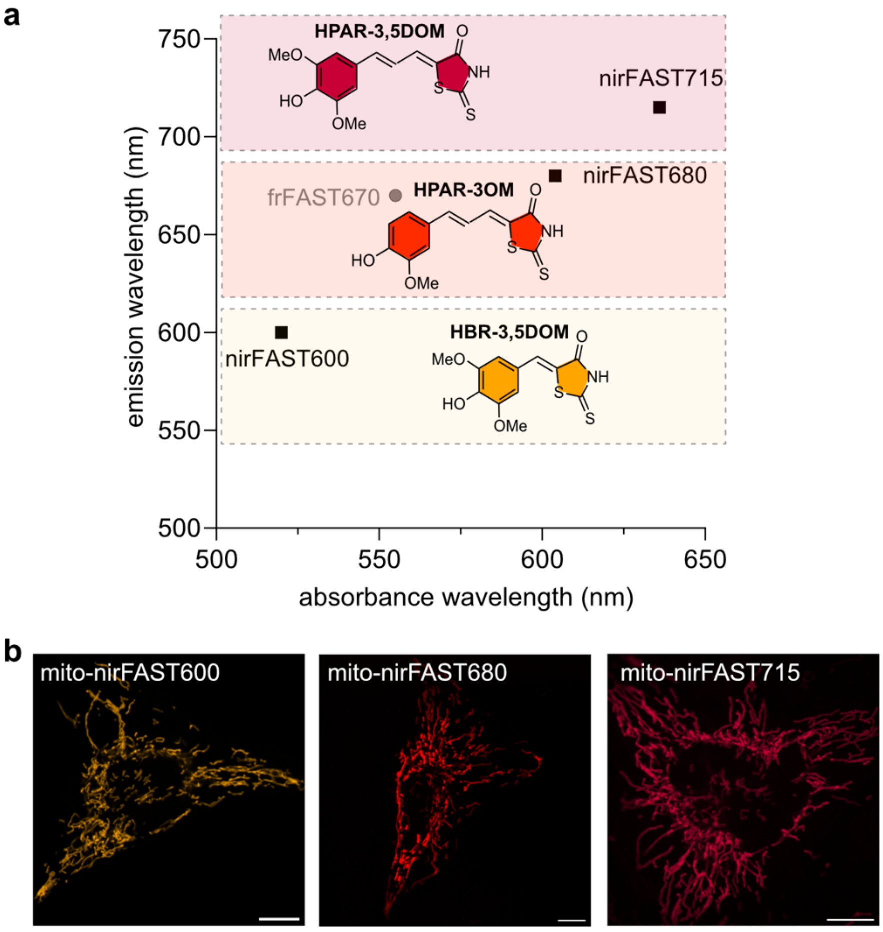


**Supplementary Figure 12. nirFAST spectral properties can be tuned from red to NIR.** (**a**) spectral properties of nirFAST600, nirFAST680 and nirFAST715 formed by assembling nirFAST with HBR-3,5DOM, HPAR-3OM and HPAR-3,5DOM. (**b**) Confocal micrographs of live HeLa cells expressing mitochrondria-targeted nirFAST treated with 10 μM HBR-3,5DOM (to assemble nirFAST600), HPAR-3OM (to assemble nirFAST680) and HPAR-3,5DOM (to assemble nirFAST715). Scale bars, 10 μm. Representative micrographs from two independent experiments for HBR-3,5DOM (n = 7 cells) and over three independent experiments for HPAR-3OM and HPAR-3,5DOM (n > 13 cells). See **Supplementary Table 8** for detailed acquisition parameters.

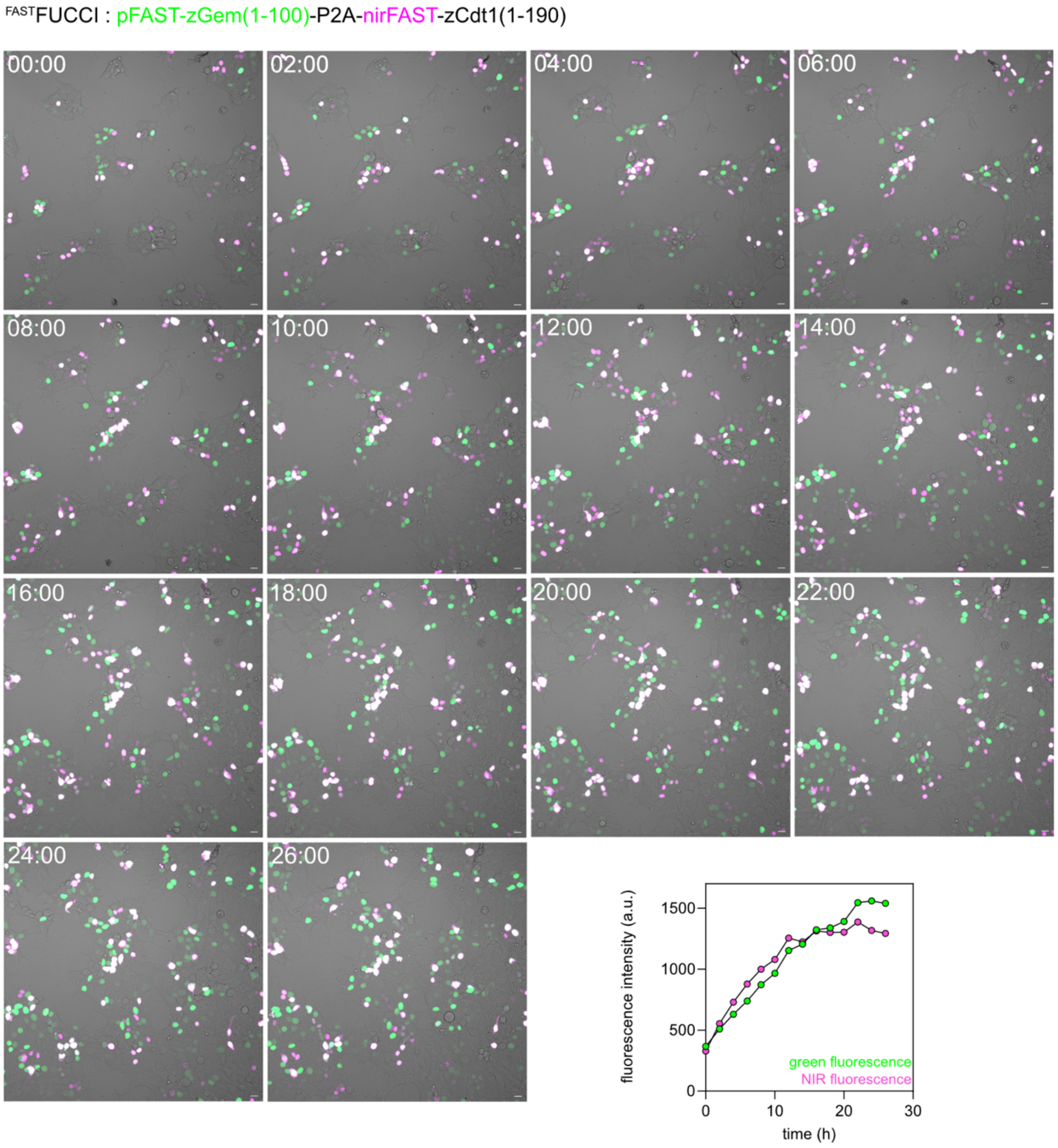


**Supplementary Figure 13. FASTFUCCI, a green-NIR fluorescent chemogenetic cell-cycle indicator.** HEK293T cells expressing FASTFUCCI and treated with 1 µM HMBR (to assemble pFAST540) and 10 µM HPAR-3,5DOM (to assemble nirFAST715) were imaged by time-lapse confocal microscopy (1 frame every 20 min) for 26 h. Relevant timepoints of designated field of view are shown. Representative micrographs from three independent experiments (see also **Supplementary Movie 3**). The graph shows the temporal evolution of the global green and NIR fluorescence, demonstrating proper cell proliferation. Scale bars, 20 µm. See **Supplementary Table 8** for detailed acquisition parameters.

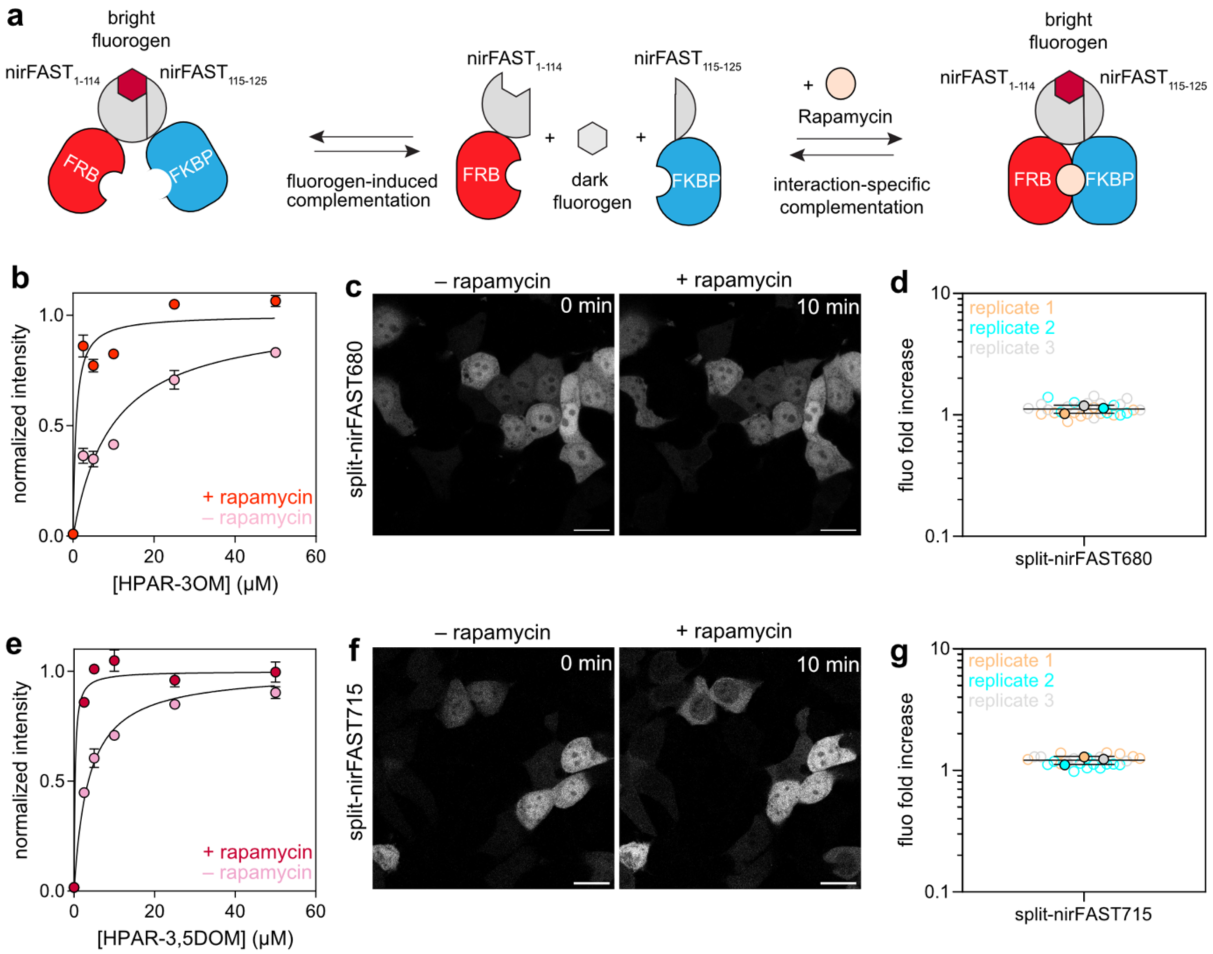


**h**

| | $EC_{50}$, + rapamycin (µM) | $EC_{50}$, – rapamycin (µM) |
|---|---|---|
| split-nirFAST680 | 0.72 | 9.6 |
| split-nirFAST715 | 0.25 | 3.6 |

**Supplementary Figure 14. Design and characterization of split-nirFAST.** split-nirFAST was obtained through bisection of nirFAST between positions 114-115. (**a**) To characterize split-nirFAST, $nirFAST_{115\text{-}125}$ was fused to FK506-binding protein (FKBP) and $nirFAST_{1\text{-}114}$ was fused to FKBP-rapamycin-binding domain (FRB). Upon addition of rapamycin, FRB and FKBP interact, and in presence of the fluorogen, their interaction can be probed with a fluorescence readout. In absence of rapamycin, fluorescence due to fluorogen-induced complementation of the two nirFAST fragments can be observed. (**b**) Normalized average NIR fluorescence (determined by flow cytometry) of about 50,000 HEK293T cells coexpressing FRB-$nirFAST_{1\text{-}114}$ and FKBP-$nirFAST_{115\text{-}125}$ treated without or with 500 nM of rapamycin, and with 0, 2.5, 5, 10, 25 or 50 µM of HPAR-3OM. Data represent the mean ± standard deviation of a technical triplicate of experiments. (**c**) Confocal micrographs of HEK293T cells labeled with 10 µM of HPAR-3OM and co-expressing FRB-$nirFAST_{1\text{-}114}$ and FKBP-$nirFAST_{115\text{-}125}$ prior and 10 minutes after treatment with 100 nM of rapamycin. Scale bars, 20 µm. (**d**) NIR fluorescence fold increase with 10 µM HPAR-3OM after 10 minutes of addition of rapamycin. n = 29 cells from three independent experiments. (**e**) Normalized average fluorescence (determined by flow cytometry) of about 50,000 HEK293T cells coexpressing FRB-$nirFAST_{1\text{-}114}$ and FKBP-$nirFAST_{115\text{-}125}$ treated without or with 500 nM of rapamycin, and with 2.5, 5, 10, 25 or 50 µM of HPAR-3,5DOM. Data represent the mean ± standard deviation of a technical triplicate of experiments. (**f**) Confocal micrographs of HEK293T cells labeled with 10 µM of HPAR-3,5DOM and coexpressing FRB-$nirFAST_{1\text{-}114}$ and FKBP-$nirFAST_{115\text{-}125}$ prior and 10 minutes after treatment with 100 nM of rapamycin. Scale bars, 20 µm. See **Supplementary Table 8** for detailed acquisition parameters. (**g**) NIR fluorescence fold increase with 10 µM HPAR-3,5DOM after 10 minutes of addition of rapamycin. n = 24 cells from three independent experiments. (**h**) Concentration of fluorogen for half-maximal complementation in presence ($EC_{50,+\ rapamycin}$) or absence ($EC_{50,-\ rapamycin}$) of rapamycin, as measured in (**b**) and (**e**). See also **Supplementary Figure 16** for the gating strategy.

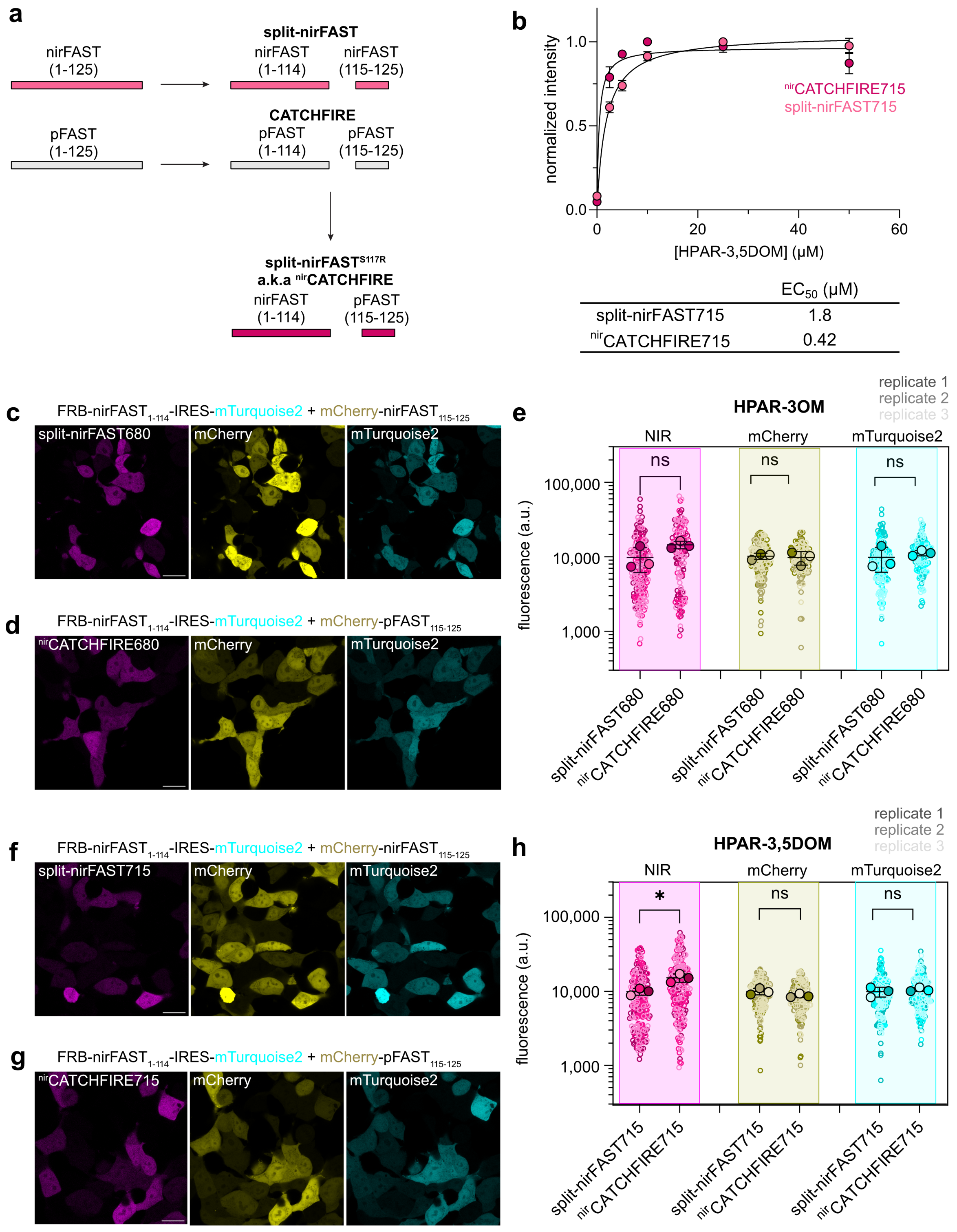


**Supplementary Figure 15. Comparison of split-nirFAST and nirCATCHFIRE.** (**a**) Design of split-nirFAST and nirCATCHFIRE. (**b**) Normalized average NIR fluorescence (determined by flow cytometry) of about 50,000 HEK293T cells co-expressing FRB-nirFAST$_{1-114}$ and either FKBP-nirFAST$_{115-125}$ (split-nirFAST) or FKBP-pFAST$_{115-125}$ (a.k.a. FKBP-$^{FIRE}$tag) treated without or with 500 nM of rapamycin, and with 0, 2.5, 5, 10, 25 or 50 μM of HPAR-3OM. Data represent the mean ± standard deviation of a technical triplicate of experiments. Concentration of fluorogen for half-maximal complementation (EC$_{50}$) values from these titration experiments are reported. See also **Supplementary Figure 16** for gating strategy. (**c-d**) Confocal micrographs of HEK293T cells co-expressing FRB-nirFAST$_{1\text{-}114,}$ and either mCherry-nirFAST$_{115\text{-}125}$ (**c**) or mCherry-pFAST$_{115\text{-}125}$ (a.k.a. mCherry-$^{FIRE}$tag) labeled with 10 µM of HPAR-3OM (**d**). Representative micrographs of n > 130 cells from three independent experiments. (**e**) NIR, red and cyan fluorescence from HEK cells expressing split-nirFAST and $^{nir}$CATCHFIRE, labeled with 10 µM HPAR-3OM. Each cell is color-coded according to the biological replicate it came from. The solid circles correspond to the mean of each biological replicate. The black line represents the mean ± SD of the three biological replicates. For split-nirFAST, 178 cells from three experiments were analyzed. For $^{nir}$CATCHFIRE, 134 cells from three experiments were analyzed. Unpaired two-tailed t-test assuming equal variance was used to compare split-nirFAST680 and $^{nir}$CATCHFIRE680 distributions. $^{ns}$P = 0.1085 (NIR), $^{ns}$P = 0.7489 (mCherry), $^{ns}$P = 0.5492 (mTurquoise2). (**f-g**) Confocal micrographs of HEK293T cells co-expressing FRB-nirFAST$_{1\text{-}114,}$ and either mCherry-nirFAST$_{115\text{-}125}$ (**f**) or mCherry-pFAST$_{115\text{-}125}$ (a.k.a. mCherry-$^{FIRE}$tag) labeled with 10 µM of HPAR-3,5DOM (**g**). Representative micrographs of n > 200 cells from three independent experiments. **(h)** NIR, red and cyan fluorescence from HEK293T cells expressing split-nirFAST and $^{nir}$CATCHFIRE, labeled with 10 µM HPAR-3,5DOM. Each cell is color-coded according to the biological replicate it came from. The solid circles correspond to the mean of each biological replicate. The black line represents the mean ± SD of the three biological replicates. For split-nirFAST, 211 cells from three experiments were analyzed. For $^{nir}$CATCHFIRE, 184 cells from three experiments were analyzed. Unpaired two-tailed t-test assuming equal variance was used to compare split-nirFAST715 and $^{nir}$CATCHFIRE715 distributions. $^{*}$P = 0.0148 (NIR), $^{ns}$P = 0.1335 (mCherry), $^{ns}$P = 0.5076 (mTurquoise2). See **Supplementary Table 8** for detailed acquisition parameters.

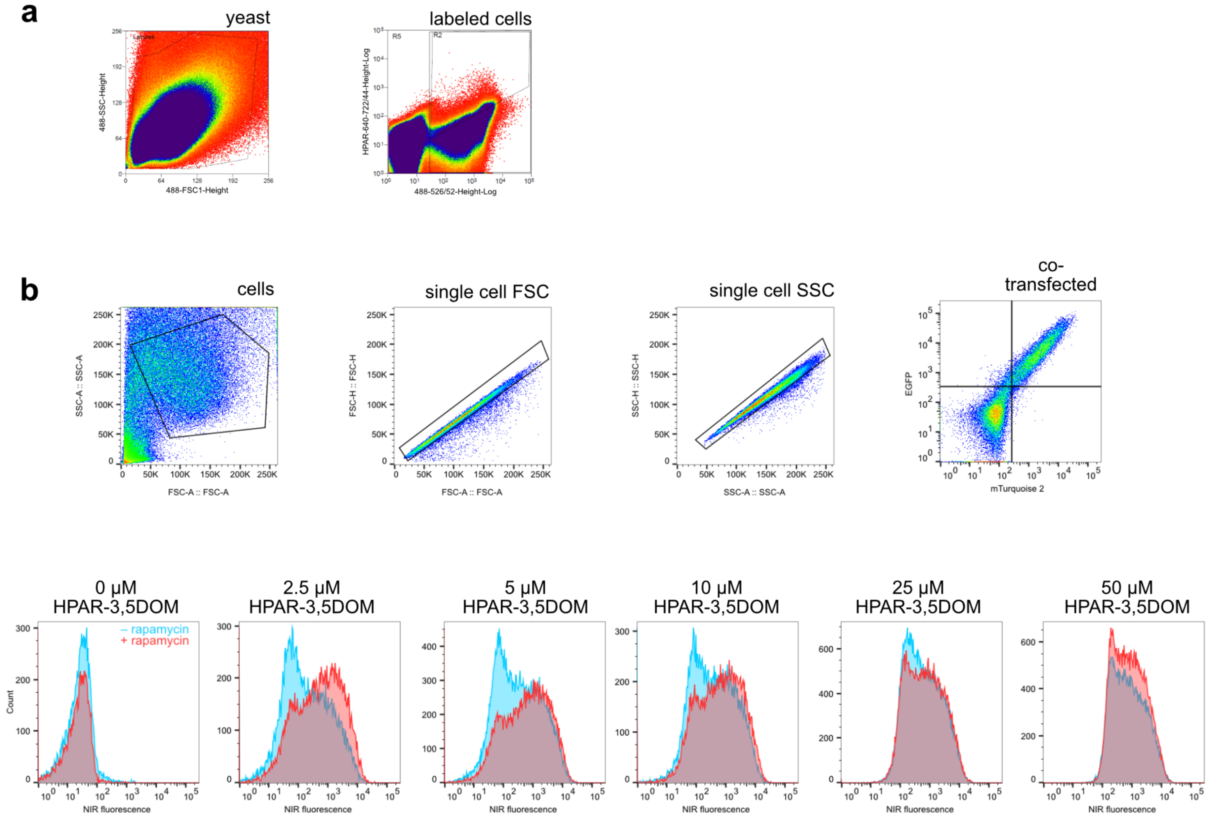


**Supplementary Figure 16. Gating strategy for the cytometry experiments. (a)** Gating strategy for the yeast display selection. **(b)** The plasmids encoding FRB-nirFAST$_{1\text{-}114}$ and FKBP-nirFAST$_{115\text{-}125}$ also enabled the expression of mTurquoise2 and EGFP respectively for transfection reporting. NIR fluorescence of doubly transfected cells enabled the quantification of the efficiency of complementation between FRB-$^{nirFIRE}$mate and FKBP-$^{FIRE}$tag in presence of different fluorogen concentrations (also see **Supplementary Figure 14**). A similar gating strategy was employed for the comparison of the complementation efficiency of FRB-nirFAST$_{1\text{-}114}$ with either mCherry-nirFAST$_{115\text{-}125}$ or mCherry-pFAST$_{115\text{-}125}$ as the plasmids express mTurquoise2 and mCherry for transfection reporting (see also **Supplementary Figure 15**).

**Supplementary Table 1. Library A clones properties with HPAR-3,5DOM in PBS pH 7.4.**

| | Mutations relative to frFAST | plasmid | $K_D$ (μM) | $\lambda_{abs}$ (nm) | ε ($mM^{-1}.cm^{-1}$) | $\lambda_{em}$ (nm) | FQY (%) |
|---|---|---|---|---|---|---|---|
| frFAST | | pAG455 | 2.6 | 610 | 19 | 709 | 6 |
| nirFAST0.1 | E46Q, I107V | pAG456 | 2.1 | 650 | 7.2 | 717 | 5.8 |
| A.R5-5 | D24E, I31V, E46Q, K78I, I107V | pAG1103 | 0.05 | 638 | 19 | 717 | 4.5 |
| A.R5-22 | Q22H, I31V, E46Q, D48G, K78I, I107V | pAG1104 | 0.10 | 642 | 32 | 717 | 4.7 |
| A.R5-48 (=nirFAST1.0) | I31V, E46Q, D48G, T70S, I107V | pAG1105 | 0.20 | 652 | 19 | 715 | 5.4 |
| A.R5-24 | I31V, E93G, | pAG1132 | 0.95 | 612 | 35 | 715 | 6 |
| A.R5-48.1 | N43S, I31V, E46Q, D48G, T70S | pAG1135 | 0.42 | 640 | 25 | 713 | 6.8 |
| A.R5-48.2 | K78I, I31V, E46Q, D48G, T70S | pAG1136 | 0.14 | 650 | 13 | 719 | 5.9 |
| A.R5-48.3 (=nirFAST1.1) | N43S, K78I, I31V, E46Q, D48G, T70S, I107V | pAG1137 | 0.31 | 640 | 34 | 713 | 6.7 |
| A.R5-18.1 | I11T, K17N, M18T, I31V, D48G, K78I, K110M, T70S | pAG1139 | 0.50 | 640 | 16 | 717 | 5.9 |
| A.R5-22.1 | Q22H, I31V, E46Q, D48G, K78I, I107V, T70S | pAG1140 | 0.45 | 650 | 16 | 713 | 5.8 |

Abbreviations are as follows: $K_D$ thermodynamic dissociation constant, $\lambda_{abs}$ wavelength of maximal absorption, ε molar absorptivity at $\lambda_{abs}$ (standard error is typically 10%), $\lambda_{em}$ wavelength of maximal emission, FQY fluorescence quantum yield.

**Supplementary Table 2. Library A clones properties with HPAR-3OM in PBS pH 7.4.**

| Clone | Mutations relative to frFAST | plasmid | $K_D$ (μM) | $\lambda_{abs}$ (nm) | ε ($mM^{-1}.cm^{-1}$) | $\lambda_{em}$ (nm) | FQY (%) |
|---|---|---|---|---|---|---|---|
| frFAST | | pAG455 | 0.6 | 555 | 45 | 670 | 21 |
| nirFAST0.1 | E46Q, I107V | pAG456 | 1.3 | 618 | 8 | 683 | 15 |
| A.R5-5 | D24E, I31V, E46Q, K78I, I107V | pAG1103 | 0.23 | 608 | 15 | 683 | 19 |
| A.R5-22 | Q22H, I31V, E46Q, D48G, K78I, I107V | pAG1104 | 0.23 | 606 | 20 | 681 | 16 |
| A.R5-48 (=nirFAST1.0) | I31V, E46Q, D48G, T70S, I107V | pAG1105 | 0.26 | 612 | 21 | 681 | 16 |
| A.R5-24 | I31V, E93G, | pAG1132 | 0.49 | 576 | 42 | 673 | 20 |
| A.R5-48.1 | N43S, I31V, E46Q, D48G, T70S | pAG1135 | 0.55 | 604 | 24 | 675 | 19 |
| A.R5-48.2 | K78I, I31V, E46Q, D48G, T70S | pAG1136 | 0.21 | 608 | 25 | 683 | 18 |
| A.R5-48.3 (=nirFAST1.1) | N43S, K78I, I31V, E46Q, D48G, T70S, I107V | pAG1137 | 0.33 | 600 | 28 | 677 | 21 |
| A.R5-18.1 | I11T, K17N, M18T, I31V, D48G, K78I, K110M, T70S | pAG1139 | 0.72 | 602 | 11 | 681 | 31 |
| A.R5-22.1 | Q22H, I31V, E46Q, D48G, K78I, I107V, T70S | pAG1140 | 0.4 | 610 | 20 | 681 | 19 |

Abbreviations are as follows: $K_D$ thermodynamic dissociation constant, $\lambda_{abs}$ wavelength of maximal absorption, ε molar absorptivity at $\lambda_{abs}$ (standard error is typically 10%), $\lambda_{em}$ wavelength of maximal emission, FQY fluorescence quantum yield.

**Supplementary Table 3. Library B clones properties with HPAR-3,5DOM in PBS pH 7.4.**

| | Mutations relative to frFAST | plasmid | $K_D$ (µM) | $\lambda_{abs}$ (nm) | $FWHM_{abs}$ (nm) | ε ($mM^{-1}.cm^{-1}$) | $\lambda_{em}$ (nm) | FQY (%) |
|---|---|---|---|---|---|---|---|---|
| B.R5-24 (= nirFAST2.0) | I31V, D36N, Q41K, N43S, E46Q, D48G, T70S, K78I, I107V | pAG1208 | 0.037 | 636 | 136 | 46 | 713 | 6.7 |
| B.R5-6 | I31V, Q41K, N43S, E46Q, G47R,D48G, T70S, K78I, I107V | pAG1225 | 0.059 | 636 | 134 | 43 | 713 | 6.6 |
| B.R5-67 | A27P, I31V, Q32R, N43S, E46Q, D48G, R52K, T70S, K78I, L88M, I107V | pAG1226 | 0.062 | 640 | 116 | 31 | 713 | 6.3 |
| B.R5-60 | G25R, I31V, K41R, N43S, E46Q, D48G, T70S, K78I, I107V | pAG1227 | 0.062 | 634 | 122 | 26 | 715 | 6.7 |
| B.R5-37 | I31V, Q41K, N43S, E46Q,D48G, T70S, K78I, I107V | pAG1228 | 0.055 | 634 | 128 | 31 | 715 | 6.2 |
| B.R5-24.7 | I31V, D36N, Q41K, N43S, E46Q, D48G, T70S, K78I | pAG1236 | 0.018 | 636 | 126 | 64 | 715 | 6.5 |
| B.R5-24.1 | G21R, I31V, D36N, Q41K, N43S, E46Q, D48G, T70S, K78I, I107V | pAG1249 | 0.022 | 632 | 128 | 64 | 713 | 5.8 |
| B.R5-24.2 | G25R,I31V, D36N, Q41K, N43S, E46Q, D48G, T70S, K78I, I107V | pAG1250 | 0.070 | 640 | 124 | 60 | 713 | 6.8 |
| B.R5-24.3 | I31V, D36N, Q41R, N43S, E46Q, D48G, T70S, K78I, I107V | pAG1251 | 0.036 | 636 | 122 | 56 | 715 | 6.6 |
| B.R5-24.4 | I31V, D36N, Q41K, N43S, E46Q, G47R, D48G, T70S, K78I, I107V | pAG1252 | 0.042 | 632 | 130 | 37 | 713 | 7.6 |
| B.R5-24.5 (nirFAST) | I31V, D36N, Q41K, N43S, E46Q, R52K, D48G, T70S, K78I, I107V | pAG1253 | 0.036 | 636 | 118 | 41 | 715 | 6.9 |
| B.R5-24.6 | I31V, D36N, Q41K, N43S, E46Q, D48G, T70S, K78I, M95T, I107V | pAG1254 | 0.059 | 632 | 130 | 40 | 715 | 6.5 |
| B.R5-24.8 | I31V, D36N, Q41K, N43S, E46Q, D48G, T70S, K78I, I107V, S117R | pAG1255 | 0.053 | 638 | 126 | 54 | 713 | 6.9 |

Abbreviations are as follows: $K_D$ thermodynamic dissociation constant, $\lambda_{abs}$ wavelength of maximal absorption, $FWHM_{abs}$ full width at half-maximal absorption, ε molar absorptivity at $\lambda_{abs}$ (standard error is typically 10%), $\lambda_{em}$ wavelength of maximal emission, FQY fluorescence quantum yield.

**Supplementary Table 4. Library B clones properties with HPAR-3OM in PBS pH 7.4.**

| | Mutations relative to frFAST | plasmid | $K_D$ (µM) | $\lambda_{abs}$ (nm) | $FWHM_{abs}$ (nm) | ε ($mM^{-1}.cm^{-1}$) | $\lambda_{em}$ (nm) | FQY (%) |
|---|---|---|---|---|---|---|---|---|
| B.R5-24 (= nirFAST2.0) | I31V, D36N, Q41K, N43S, E46Q, D48G, T70S, K78I, I107V | pAG1208 | 0.01 | 600 | 115 | 52 | 679 | 20 |
| B.R5-6 | I31V, Q41K, N43S, E46Q, G47R,D48G, T70S, K78I, I107V | pAG1225 | 0.067 | 594 | 124 | 55 | 677 | 18 |
| B.R5-67 | A27P, I31V, Q32R, N43S, E46Q, D48G, R52K, T70S, K78I, L88M, I107V | pAG1226 | 0.10 | 604 | 100 | 45 | 681 | 18 |
| B.R5-60 | G25R, I31V, K41R, N43S, E46Q, D48G, T70S, K78I, I107V | pAG1227 | 0.063 | 604 | 116 | 45 | 677 | 18 |
| B.R5-37 | I31V, Q41K, N43S, E46Q,D48G, T70S, K78I, I107V | pAG1228 | 0.070 | 600 | 120 | 43 | 679 | 17 |
| B.R5-24.7 | I31V, D36N, Q41K, N43S, E46Q, D48G, T70S, K78I | pAG1236 | 0.067 | 596 | 116 | 55 | 679 | 19 |
| B.R5-24.1 | G21R, I31V, D36N, Q41K, N43S, E46Q, D48G, T70S, K78I, I107V | pAG1249 | 0.042 | 596 | 116 | 56 | 679 | 17 |
| B.R5-24.2 | G25R,I31V, D36N, Q41K, N43S, E46Q, D48G, T70S, K78I, I107V | pAG1250 | 0.064 | 596 | 118 | 50 | 679 | 20 |
| B.R5-24.3 | I31V, D36N, Q41R, N43S, E46Q, D48G, T70S, K78I, I107V | pAG1251 | 0.056 | 602 | 114 | 47 | 679 | 20 |
| B.R5-24.4 | I31V, D36N, Q41K, N43S, E46Q, G47R, D48G, T70S, K78I, I107V | pAG1252 | 0.077 | 598 | 118 | 52 | 675 | 21 |
| B.R5-24.5 (nir-FAST) | I31V, D36N, Q41K, N43S, E46Q, R52K, D48G, T70S, K78I, I107V | pAG1253 | 0.055 | 604 | 106 | 52 | 679 | 20 |
| B.R5-24.6 | I31V, D36N, Q41K, N43S, E46Q, D48G, T70S, K78I, M95T, I107V | pAG1254 | 0.12 | 602 | 118 | 54 | 675 | 20 |
| B.R5-24.8 | I31V, D36N, Q41K, N43S, E46Q, D48G, T70S, K78I, I107V, S117R | pAG1255 | 0.062 | 600 | 116 | 52 | 677 | 20 |

Abbreviations are as follows: $K_D$ thermodynamic dissociation constant, $\lambda_{abs}$ wavelength of maximal absorption, $FWHM_{abs}$ full width at half-maximal absorption, ε molar absorptivity at $\lambda_{abs}$ (standard error is typically 10%), $\lambda_{em}$ wavelength of maximal emission, FQY fluorescence quantum yield.

**Supplementary Table 5. Physico-chemical properties of nirFAST with HBR fluorogens in PBS pH 7.4**

| Fluorogen | $K_D$ (µM) | $\lambda_{abs}$ (nm) | $\lambda_{em}$ (nm) | FQY (%) |
|---|---|---|---|---|
| HMBR | 0.062 | 448 | 549 | 2 |
| HBR-2,5DM | 0.032 | 510 | 561 | 11 |
| HBR-3,5DM | 0.02 | 520 | 567 | 37 |
| HBR-3,5DOM | 0.14 | 516 | 599 | 24 |

Abbreviations are as follows: $K_D$ thermodynamic dissociation constant, $\lambda_{abs}$ wavelength of maximal absorption, $\lambda_{em}$ wavelength of maximal emission, FQY fluorescence quantum yield.

**Supplementary Table 6. Photophysical properties of emiRFP670, miRFP713, nirFAST680 and nirFAST715.**

| | $\lambda_{abs}$ (nm) | ε ($M^{-1}.cm^{-1}$) | $\lambda_{em}$ (nm) | FQY (%) |
|---|---|---|---|---|
| emiRFP670 | 642 | 87.4 | 670 | 14 |
| miRFP713 | 690 | 99 | 713 | 7 |
| nirFAST680 | 604 | 52 | 680 | 21 |
| nirFAST715 | 636 | 41 | 715 | 6.9 |

Abbreviations are as follows: $\lambda_{abs}$ wavelength of maximal absorption, ε molar absorptivity at $\lambda_{abs}$ (standard error is typically 10%), $\lambda_{em}$ wavelength of maximal emission, FQY fluorescence quantum yield.

**Supplementary Table 7. Binding affinity of nirFASTS117R with HBR and HPAR fluorogens in PBS pH 7.4.**

| Fluorogen | $K_D$ (µM) |
|---|---|
| HMBR | 0.038 |
| HBR-2,5DM | 0.013 |
| HBR-3,5DM | 0.02 |
| HBR-3,5DOM | 0.14 |
| HPAR-3OM | 0.01 |
| HPAR-3,5DOM | 0.016 |

### Supplementary Table 8. Imaging settings used in this study

| Figure | Panel | Fluorescent reporter | Excitation (nm) | Emission (nm) | Imaging setup | Comments |
|---|---|---|---|---|---|---|
| Fig. 1 | | nirFAST680 | 650–675 | 715/30 | Biorad gel imager | |
| | | nirFAST715 | 650–675 | 715/30 | Biorad gel imager | |
| Fig. 2 | panel a | nirFAST680 | 633 | 650-800 | Leica SP5 63 × oil | |
| | | H2B-nirFAST680 | 633 | 650-800 | Leica SP5 63 × oil | |
| | | mito-nirFAST680 | 639 | 650-757 | LSM980 inverted 63 × oil | |
| | | MAP4-nirFAST680 | 633 | 650-800 | Leica SP5 63 × oil | |
| | | nirFAST680-Cb5 | 639 | 650-757 | LSM980 inverted 63 × oil | |
| | | nirFAST715 | 633 | 650-800 | Leica SP5 63 × oil | |
| | | H2B-nirFAST715 | 633 | 650-800 | Leica SP5 63 × oil | |
| | | mito-nirFAST715 | 639 | 650-757 | LSM980 inverted 63 × oil | |
| | | MAP4-nirFAST715 | 633 | 650-800 | Leica SP5 63 × oil | |
| | | nirFAST715-Cb5 | 639 | 650-757 | LSM980 inverted 63 × oil | |
| | panels b-c | H2B-nirFAST680 (live and fixed) | 639 | 650-757 | LSM980 inverted 20 × dry | |
| | | H2B-nirFAST715 (live and fixed) | 639 | 650-757 | LSM980 inverted 20 × dry | |
| | panel d | ECFP-giantin | 405 | 420-500 | LSM980 inverted 63 × oil | |
| | | mito-pFAST540 | 488 | 508-570 | LSM980 inverted 63 × oil | |
| | | lyn11-mCherry | 561 | 600-632 | LSM980 inverted 63 × oil | |
| | | H2B-nirFAST715 | 639 | 680-757 | LSM980 inverted 63 × oil | |
| Fig. 3 | panels a,c | NIR reporter-P2A-EGFP | 488/639 | 500-600 / 650-790 | Leica SP5 63 × oil | |
| Fig. 4 | panel a | MAP4-nirFAST600 | 561 | Abberior STAR Orange | Abberior STEDYCON 775 depletion laser, 100 × glycerol | pixel size 25-30 nm |
| | panel b | MAP4-nirFAST680 | 640 | Abberior STAR Red | Abberior STEDYCON 775 depletion laser, 100 × glycerol | pixel size 25-30 nm |
| Fig. 5 | panel b | frFAST<br>nirFAST680<br>EGFP | 642<br>642<br>491 | 500-550<br>500-550<br>660-750 | Spinning-disk Confocal 10 × CFI Plan APO LBDA, NA 0.45 | |
| | panel c | frFAST<br>nirFAST715<br>EGFP | 642<br>642<br>491 | 500-550<br>500-550<br>660-750 | Spinning-disk Confocal 10 × CFI Plan APO LBDA, NA 0.45 | |
| | panel d | emiRFP670<br>nirFAST715<br>EGFP | 642<br>642<br>491 | 500-550<br>500-550<br>660-750 | Spinning-disk Confocal 10 × CFI Plan APO LBDA, NA 0.45 | |

| | | | | | | |
|---|---|---|---|---|---|---|
| | panel e | H2B-pFAST540<br>mCherry<br>mito-nirFAST715 | 491<br>561<br>642 | 500-550<br>580-653<br>660-750 | Spinning-disk Confocal, 100 × APO VC, NA 1.4 oil | Also for Supplementary Movie 1 |
| | panel g | nirFAST-P2A-eGFP #1 | 488 / 639 | 500-570/ 650-757 | LSM980 upright 20 × water | |
| | | nirFAST-P2A-eGFP#1 projection | 639 | 650-757 | LSM980 upright 20 × water | z-stack (12 stacks, Z-step = 5 µm ) |
| | | nirFAST-P2A-eGFP #2 | 488 / 639 | 500-570/ 650-757 | LSM980 upright 20 × water | |
| | | nirFAST-P2A-eGFP#2 projection | 639 | 650-757 | LSM980 upright 20 × water | z-stack (14 stacks, Z-step = 5 µm ) |
| **Fig. 6** | panel a | $^{FAST}$FUCCI | 488 / 639 | 508-570 and 650-757 | LSM980 inverted 20 × dry | 1 frame / 20 min Also for Supplementary Movie 3 |
| **Fig. 7** | panel b,c | EGFP | 488 | 508-570 | LSM980 inverted 63 × oil | 1 frame / min |
| | | nirCATCHFIRE680 | 639 | 650-757 | LSM980 inverted 63 × oil | 1 frame / min Also for Supplementary Movie 5 |
| | | nirCATCHFIRE715 | 639 | 650-757 | LSM980 inverted 63 × oil | 1 frame / min Also for Supplementary Movie 6 |
| | panel e,f | mCherry | 561 | 609-663 | Nikon Inverted Eclipse Ti-E, 100 × oil | 1 frame/min, Z-stack (11 stacks, 0.4 µm step) Also for Supplementary Movie 4 |
| | | nirCATCHFIRE715 | 633 | 700-775 | | |
| **Fig. S5** | panel a,b | NIR reporter-P2A-EGFP | 488 / 633 | 500-600 / 650-790 | Leica SP5 63 × oil | |
| **Fig S7** | panel a,b | nirFAST-P2A-EGFP | 639 | 650-757 | LSM980 inverted 40 × oil | Timelapse 51 frames 1 frame / 2s |
| **Fig. S9** | panels a-c | H2B-mCherry | 561 | 600-variable (indicated) | LSM980 inverted 63 × oil | |
| | | mito-NIR-reporter | 639 | 650-757 | LSM980 inverted 63 × oil | |
| **Fig. S10** | | mito-pFAST540 | 488 | 508-570 | LSM980 inverted 63 × oil | |
| | | H2B-nirFAST715 | 639 | 650-757 | LSM980 inverted 63 × oil | |
| **Fig. S11** | | NIR reporter-P2A-EGFP | 639 | 650-757 | LSM980 inverted 63 × oil | Timelapse, 500 frames, 1 frame / 2s, laser power 3.4 mW |
| **Fig. S12** | panel b | mito-nirFAST600 | 561 | 570-694 | LSM980 inverted 63 × oil | |
| | | mito-nirFAST680 | 633 | 650-790 | Leica SP5 63 × oil | |
| | | mito-nirFAST715 | 633 | 650-790 | Leica SP5 63 × oil | |
| **Fig. S13** | | $^{FAST}$FUCCI | 488 / 639 | 508-570 / 650-757 | LSM980 inverted 20 × dry | 1 frame / 20min Also for Supplementary Movie 3 |
| **Fig. S14** | panels c and f | split-nirFAST680 | 488 / 639 | 508-570 / 650-757 | LSM980 inverted 63 × oil | 1 frame / 30s |

| | | | | | | |
|---|---|---|---|---|---|---|
| | | split-nirFAST715 | 488 / 639 | 508-570 / 650-757 | LSM980 inverted 63 × oil | 1 frame / 30s |
| **Fig. S15** | panels c-g | Split-nirFAST / $^{nir}$CATCHFIRE | 445 | 473-500 | LSM980 inverted 63 × oil | |
| | | | 561 | 579-606 | LSM980 inverted 63 × oil | |
| | | | 639 | 680-757 | LSM980 inverted 63 × oil | |

**Supplementary Table 9. Plasmids used in this study**

| Vector | ORF | ORF sequences |
|---|---|---|
| | frFAST | atggagcatgttgcctttggcagtgaggacatcgagaacactttggccaaaatggacga<br>cggacaactggatgggttggcctttggcgcaattcagctcgatggtgacgggaatatcct<br>gcagtacaatgctgctgaaggagacatcacaggcagagatcccaaacaggtgattgg<br>gaagaacttattcaaggatgttgcacctggaacggtttcctccgggttttacggcaaattca<br>aggaaggcgtagcgtcagggaatctgaacaccatgttcgaatggatgataccgacaa<br>gcaggggaccaaccaaggtcaagatacacatgaagaaagccctttccggtgacagct<br>attgggtctttgtgaaacgggtg |
| | nirFAST | atggagcatgttgcctttggcagtgaggacatcgagaacactttggccaaaatggacga<br>cggacaactggatgggttggcctttggcgcagttcagctcgatggtaacgggaatatcct<br>gaagtacagtgctgctcagggaggcatcacaggcaaagatcccaaacaggtgattgg<br>gaagaacttattcaaggatgttgcaccaggatcggtttcctccgggttttacggcatattca<br>aggaaggcgtagcgtcagggaatctgaacaccatgttcgaatggatgataccgacaa<br>gcagggggccaactaaggtcaaggtgcatatgaagaaagccctttccggtgacagcta<br>ttgggtctttgtgaaacgggtg |
| | nirFAST$_{1-114}$ | atggagcatgttgcctttggcagtgaggacatcgagaacactttggccaaaatggacga<br>cggacaactggatgggttggcctttggcgcagttcagctcgatggtaacgggaatatcct<br>gaagtacagtgctgctcagggaggcatcacaggcaaagatcccaaacaggtgattgg<br>gaagaacttattcaaggatgttgcaccaggatcggtttcctccgggttttacggcatattca<br>aggaaggcgtagcgtcagggaatctgaacaccatgttcgaatggatgataccgacaa<br>gcagggggccaactaaggtcaaggtgcatatgaagaaagccctttcc |
| | nirFAST$_{115-125}$ | ggtgacagctattgggtctttgtgaaacgggtg |
| pAG1253 | 6 ×His-Thrombincs-nirFAST | atgggcagcagccatcatcatcatcatcacagcagcggcctggtgccgcgcggcagc<br>catatggctagcatggagcatgttgcctttggcagtgaggacatcgagaacactttggcc<br>aaaatggacgacggacaactggatgggttggcctttggcgcagttcagctcgatggtaa<br>cgggaatatcctgaagtacagtgctgctcagggaggcatcacaggcaaagatcccaa<br>acaggtgattgggaagaacttattcaaggatgttgcaccaggatcggtttcctccgggtttt<br>acggcatattcaaggaaggcgtagcgtcagggaatctgaacaccatgttcgaatggat<br>gataccgacaagcagggggccaactaaggtcaaggtgcatatgaagaaagccctttc<br>cggtgacagctattgggtctttgtgaaacgggtg |
| pAG1329 | nirFAST-P2A-EGFP-cMyc | atggagcatgttgcctttggcagtgaggacatcgagaacactttggccaaaatggacga<br>cggacaactggatgggttggcctttggcgcagttcagctcgatggtaacgggaatatcct<br>gaagtacagtgctgctcagggaggcatcacaggcaaagatcccaaacaggtgattgg<br>gaagaacttattcaaggatgttgcaccaggatcggtttcctccgggttttacggcatattca<br>aggaaggcgtagcgtcagggaatctgaacaccatgttcgaatggatgataccgacaa<br>gcagggggccaactaaggtcaaggtgcatatgaagaaagccctttccggtgacagcta<br>ttgggtctttgtgaaacgggtgggaagcggagctactaacttcagcctgctgaagcagg<br>ctggagacgtggaggagaaccctggacctatggtgagcaagggcgaggagctgttca<br>ccggggtggtgcccatcctggtcgagctggacggcgacgtaaacggccacaagttca<br>gcgtgtccggcgagggcgagggcgatgccacctacggcaagctgaccctgaagttca<br>tctgcaccaccggcaagctgcccgtgccctggcccaccctcgtgaccaccctgacctac<br>ggcgtgcagtgcttcagccgctaccccgaccacatgaagcagcacgacttcttcaagtc<br>cgccatgcccgaaggctacgtccaggagcgcaccatcttcttcaaggacgacggcaa<br>ctacaagacccgcgccgaggtgaagttcgagggcgacaccctggtgaaccgcatcga<br>gctgaagggcatcgacttcaaggaggacggcaacatcctggggcacaagctggagta<br>caactacaacagccacaacgtctatatcatggccgacaagcagaagaacggcatca<br>aggtgaacttcaagatccgccacaacatcgaggacggcagcgtgcagctcgccgacc<br>actaccagcagaacacccccatcggcgacggccccgtgctgctgcccgacaaccact<br>acctgagcacccagtccgccctgagcaaagaccccaacgagaagcgcgatcacatg<br>gtcctgctggagttcgtgaccgccgccgggatcactctcggcatggacgaggaacaaa<br>agcttatttctgaagaggacttg |

| | | |
|---|---|---|
| pAG1334 | frFAST-<br>P2A-<br>EGFP-<br>cMyc | atggagcatgttgcctttggcagtgaggacatcgagaacactttggccaaaatggacga<br>cggacaactggatgggttggcctttggcgcaattcagctcgatggtgacgggaatatcct<br>gcagtacaatgctgctgaaggagacatcacaggcagagatcccaaacaggtgattgg<br>gaagaacttattcaaggatgttgcacctggaacggtttcctccgggttttacggcaaattca<br>aggaaggcgtagcgtcagggaatctgaacaccatgttcgaatggatgataccgacaa<br>gcaggggaccaaccaaggtcaagatacacatgaagaaagccctttccggtgacagct<br>attgggtctttgtgaaacgggtgggaagcggagctactaacttcagcctgctgaagcag<br>gctggagacgtggaggagaaccctggacctatggtgagcaagggcgaggagctgttc<br>accggggtggtgcccatcctggtcgagctggacggcgacgtaaacggccacaagttc<br>agcgtgtccggcgagggcgagggcgatgccacctacggcaagctgaccctgaagttc<br>atctgcaccaccggcaagctgcccgtgccctggcccaccctcgtgaccaccctgaccta<br>cggcgtgcagtgcttcagccgctaccccgaccacatgaagcagcacgacttcttcaagt<br>ccgccatgcccgaaggctacgtccaggagcgcaccatcttcttcaaggacgacggca<br>actacaagacccgcgccgaggtgaagttcgagggcgacaccctggtgaaccgcatcg<br>agctgaagggcatcgacttcaaggaggacggcaacatcctggggcacaagctggagt<br>acaactacaacagccacaacgtctatatcatggccgacaagcagaagaacggcatc<br>aaggtgaacttcaagatccgccacaacatcgaggacggcagcgtgcagctcgccgac<br>cactaccagcagaacacccccatcggcgacggccccgtgctgctgcccgacaacca<br>ctacctgagcacccagtccgccctgagcaaagaccccaacgagaagcgcgatcaca<br>tggtcctgctggagttcgtgaccgccgccgggatcactctcggcatggacgaggaaca<br>aaagcttatttctgaagaggacttg |
| pAG1335 | emiRFP67<br>0-P2A-<br>EGFP-<br>cMyc | atggcggaaggatccgtcgccaggcagcctgacctcttgacctgcgaacatgaagag<br>atccacctcgccggctcgatccagccgcatggcgcgcttctggtcgtcagcgaacatga<br>tcatcgcgtcatccaggccagcgccaacgccgcggaatttctgaatctcggaagcgtac<br>tcggcgttccgctcgccgagatcgacggcgatctgttgatcaagatcctgccgcatctcg<br>atcccaccgccgaaggcatgccggtcgcggtgcgctgccggatcggcaatccctctac<br>ggagtactgcggtctgatgcatcggcctccggaaggcgggctgatcatcgaactcgaa<br>cgtgccggcccgtcgatcgatctgtcaggcacgctggcgccggcgctggagcggatcc<br>gcacggcgggttcactgcgcgcgctgtgcgatgacaccgtgctgctgtttcagcagtgca<br>ccggctacgaccgggtgatggtgtatcgtttcgatgagcaaggccacggcctggtattct<br>ccgagtgccatgtgcctgggctcgaatcctatttcggcaaccgctatccgtcgtcgactgt<br>cccgcagatggcgcggcagctgtacgtgcggcagcgcgtccgcgtgctggtcgacgtc<br>acctatcagccggtgccgctggagccgcggctgtcgccgctgaccgggcgcgatctcg<br>acatgtcgggctgcttcctgcgctcgatgtcgccgtgccatctgcagttcctgaaggacat<br>gggcgtgcgcgccaccctggcggtgtcgctggtggtcggcggcaagctgtggggcctg<br>gttgtctgtcaccattatctgccgcgcttcatccgtttcgagctgcgggcgatctgcaaacg<br>gctcgccgaaaggatcgcgacgcggatcaccgcgcttgagagcggaagcggagcta<br>ctaacttcagcctgctgaagcaggctggagacgtggaggagaaccctggacctatggt<br>gagcaagggcgaggagctgttcaccggggtggtgcccatcctggtcgagctggacgg<br>cgacgtaaacggccacaagttcagcgtgtccggcgagggcgagggcgatgccacct<br>acggcaagctgaccctgaagttcatctgcaccaccggcaagctgcccgtgccctggcc<br>caccctcgtgaccaccctgacctacggcgtgcagtgcttcagccgctaccccgaccac<br>atgaagcagcacgacttcttcaagtccgccatgcccgaaggctacgtccaggagcgca<br>ccatcttcttcaaggacgacggcaactacaagacccgcgccgaggtgaagttcgagg<br>gcgacaccctggtgaaccgcatcgagctgaagggcatcgacttcaaggaggacggc<br>aacatcctggggcacaagctggagtacaactacaacagccacaacgtctatatcatgg<br>ccgacaagcagaagaacggcatcaaggtgaacttcaagatccgccacaacatcgag<br>gacggcagcgtgcagctcgccgaccactaccagcagaacacccccatcggcgacgg<br>ccccgtgctgctgcccgacaaccactacctgagcacccagtccgccctgagcaaaga<br>ccccaacgagaagcgcgatcacatggtcctgctggagttcgtgaccgccgccgggatc<br>actctcggcatggacgaggaacaaaagcttatttctgaagaggacttg |

| | | |
|---|---|---|
| pAG1337 | miRFP713<br>-P2A-<br>EGFP-<br>cMyc | atggcggaaggctccgtcgccaggcagcctgacctcttgacctgcgacgatgagccga<br>tccatatccccggtgccatccaaccgcatggactgctgctcgccctcgccgccgacatg<br>acgatcgttgccggcagcgacaaccttcccgaactcaccggactggcgatcggcgccc<br>tgatcggccgctctgcggccgatgtcttcgactcggagacgcacaaccgtctgacgatc<br>gccttggccgagcccggggcggccgtcggagcaccgatcactgtcggcttcacgatgc<br>gaaaggacgcaggcttcatcggctcctggcatcgccatgatcagctcatcttcctcgagc<br>tcgagcctccccagcgggacgtcgccgagccgcaggcgttcttccgccgcaccaaca<br>gcgccatccgccgcctgcaggccgccgaaaccttggaaagcgcctgcgccgccgcg<br>gcgcaagaggtgcggaagattaccggcttcgatcgggtgatgatctatcgcttcgcctcc<br>gacttcagcggcgaagtgatcgcagaggatcggtgcgccgaggtcgagtcaaaacta<br>ggcctgcactatcctgcctcaaccgtgccggcgcaggcccgtcggctctataccatcaa<br>cccggtacggatcattcccgatatcaattatcggccggtgccggtcaccccagacctca<br>atccggtcaccgggcggccgattgatcttagcttcgccatcctgcgcagcgtctcgcccgt<br>ccatctggagttcatgcgcaacataggcatgcacggcacgatgtcgatctcgattttgcg<br>cggcgagcgactgtggggattgatcgtttgccatcaccgaacgccgtactacgtcgatct<br>cgatggccgccaagcctgcaagagggtcgccgagaggctggccactcagatcggcgt<br>gatggaagagggaagcggagctactaacttcagcctgctgaagcaggctggagacgt<br>ggaggagaaccctggacctatggtgagcaagggcgaggagctgttcaccggggtggt<br>gcccatcctggtcgagctggacggcgacgtaaacggccacaagttcagcgtgtccggc<br>gagggcgagggcgatgccacctacggcaagctgaccctgaagttcatctgcaccacc<br>ggcaagctgcccgtgccctggcccaccctcgtgaccaccctgacctacggcgtgcagt<br>gcttcagccgctaccccgaccacatgaagcagcacgacttcttcaagtccgccatgccc<br>gaaggctacgtccaggagcgcaccatcttcttcaaggacgacggcaactacaagacc<br>cgcgccgaggtgaagttcgagggcgacaccctggtgaaccgcatcgagctgaaggg<br>catcgacttcaaggaggacggcaacatcctggggcacaagctggagtacaactacaa<br>cagccacaacgtctatatcatggccgacaagcagaagaacggcatcaaggtgaactt<br>caagatccgccacaacatcgaggacggcagcgtgcagctcgccgaccactaccagc<br>agaacacccccatcggcgacggccccgtgctgctgcccgacaaccactacctgagca<br>cccagtccgccctgagcaaagaccccaacgagaagcgcgatcacatggtcctgctgg<br>agttcgtgaccgccgccgggatcactctcggcatggacgaggaacaaaagcttatttct<br>gaagaggacttg |
| pAG1372 | H2B-<br>nirFAST-<br>cMyc | atgcccgaacctgcgaagtcagcgcccgctcccaaaaaaggctctaaaaaagctgtc<br>gccaagacccagaagaaggggataagaaaaggcgtaagaccaggaaagagagt<br>tacgccatttacgtgtacaaagtactaaaacaagtccacccggacactggcatctcctca<br>aaggcgatgggcattatgaactcatttgtaaacgacatcttcgagcgcatcgccggaga<br>agcgtcgcgcctggcgcattacaacaagcgctccactatcacatcccgggagatccag<br>acggccgtgcgcctgctcctgcccggagaactggccaaacacgctgtgtctgagggca<br>caaaggccgtgaccaagtacaccagctccaagggcggaggctccggaggcggatct<br>gccaccatggagcatgttgcctttggcagtgaggacatcgagaacactttggccaaaat<br>ggacgacggacaactggatgggttggcctttggcgcagttcagctcgatggtaacggg<br>aatatcctgaagtacagtgctgctcagggaggcatcacaggcaaagatcccaaacag<br>gtgattgggaagaacttattcaaggatgttgcaccaggatcggtttcctccgggttttacgg<br>catattcaaggaaggcgtagcgtcagggaatctgaacaccatgttcgaatggatgatac<br>cgacaagcaggggccaactaaggtcaaggtgcatatgaagaaagccctttccggtg<br>acagctattgggtctttgtgaaacgggtgggatccgaacaaaagcttatttctgaagagg<br>acttg |
| pAG1375 | mito-<br>nirFAST-<br>cMyc | atgtccgtcctgacgccgctgctgctgcggggcttgacaggctcggcccggcggctccc<br>agtgccgcgcgccaagatccattcgttgagatctgccaccatggagcatgttgcctttggc<br>agtgaggacatcgagaacactttggccaaaatggacgacggacaactggatgggttg<br>gcctttggcgcagttcagctcgatggtaacgggaatatcctgaagtacagtgctgctcag<br>ggaggcatcacaggcaaagatcccaaacaggtgattgggaagaacttattcaaggat<br>gttgcaccaggatcggtttcctccgggttttacggcatattcaaggaaggcgtagcgtcag<br>ggaatctgaacaccatgttcgaatggatgataccgacaagcaggggccaactaagg<br>tcaaggtgcatatgaagaaagccctttccggtgacagctattgggtctttgtgaaacgggt<br>gggatccgaacaaaagcttatttctgaagaggacttg |

| | | |
|---|---|---|
| pAG1377 | nirFAST-Cb5 | atggagcatgttgcctttggcagtgaggacatcgagaacactttggccaaaatggacgacggacaactggatgggttggcctttggcgcagttcagctcgatggtaacgggaatatcctgaagtacagtgctgctcagggaggcatcacaggcaaagatcccaaacaggtgattgggaagaacttattcaaggatgttgcaccaggatcggtttcctccgggttttacggcatattcaaggaaggcgtagcgtcagggaatctgaacaccatgttcgaatggatgataccgacaagcaggggggccaactaaggtcaaggtgcatatgaagaaagccctttccggtgacagctattgggtctttgtgaaacgggtgtccggactcagatctatcaccaccgtggagtccaactcctcctggtggaccaactgggtgatccccgccatctccgccctggtggtggccctgatgtaccgcctatacatggccgaggac |
| pAG1380 | MAP4-nirFAST-cMyc | atggtgtcccggcaagaagaagcaaaggctgctgtaggtgtgactggaaatgacatcactaccccgccaaacaaggagccaccaccaagcccagaaaagaaagcaaagcctttggccaccactcaacctgcaaagacttcaacatcgaaagccaaaacacagcccacttctctccctaagcaaccagctcccaccacctctggtgggttgaataaaaaacccatgagcctcgcctcaggctcagtgccagctgccccacacaaacgccctgctgctgccactgctactgccaggccttccaccctacctgccagagacgtgaagccaaagccaattacagaagctaaggttgccgaaaagcggacctctccatccaagccttcatctgccccagccctcaaacctggacctaaaaccaccccaaccgtttcaaaagccacatctccctcaactcttgtttccactggaccaagtagtagaagtccagctacaactctgcctaagaggccaaccagcatcaagactgaggggaaacctgctgatgtcaaaaggatgactgctaagtctgcctcagctgacttgagtcgctcaaagaccacctctgccagttctgtgaagagaaacaccactcccactggggcagcacccccagcagggatgacttccactcgagtcaagcccatgtctgcacctagccgctcttctggggctctttctgtggacaagaagcccacttccactaagcctagctcctctgctcccagggtgagccgcctggccacaactgtttctgcccctgacctgaagagtgttcgctccaaggtcggctctacagaaaacatcaaacaccagcctggaggaggccgggccaaggtagagaaaaaaacagaggcagctaccacagctgggaagcctgaacctaatgcagtcactaaagcagccggctccattgcgagtgcacagaaaccgcctgctgggaaagtccagatagtatccaaaaaagtgagctacagtcatattcaatccaagtgtgtttccaaggacaatattaagcatgtccctggatgtggcaatgttcagattcagaacaagaaagtggacatatccaaggtctcctccaagtgtgggtccaaagctaatatcaagcacaagcctggtggaggagatgtcaagattgaaagtcagaagttgaacttcaaggagaaggcccaagccaaagtgggaggcggattcgcggatccaccggtcgccaccatgagtgtgattaaaccagacatggagcatgttgcctttggcagtgaggacatcgagaacactttggccaaaatggacgacggacaactggatgggttggcctttggcgcagttcagctcgatggtaacgggaatatcctgaagtacagtgctgctcagggaggcatcacaggcaaagatcccaaacaggtgattgggaagaacttattcaaggatgttgcaccaggatcggtttcctccgggttttacggcatattcaaggaaggcgtagcgtcagggaatctgaacaccatgttcgaatggatgataccgacaagcaggggggccaactaaggtcaaggtgcatatgaagaaagccctttccggtgacagctattgggtctttgtgaaacgggtgggatccgaacaaaagcttatttctgaagaggacttg |

| | | |
|---|---|---|
| pAG1384 | cMyc-FKBP-nirFAST$_{115-125}$-IRES-HA-EGFP | atggaacaaaagcttatttctgaagaggacttggaattcggagtgcaggtggaaaccat<br>ctccccaggagacgggcgcaccttccccaagcgcggccagacctgcgtggtgcacta<br>caccgggatgcttgaagatggaaagaaatttgattcctcccgggacagaaacaagccc<br>tttaagtttatgctaggcaagcaggaggtgatccgaggctgggaagaaggggttgccca<br>gatgagtgtgggtcagagagccaaactgactatatctccagattatgcctatggtgccact<br>gggcacccaggcatcatcccaccacatgccactctcgtcttcgatgtggagcttctaaaa<br>ctggaagaatccggaggaggcggcagcggcggagggggatccggtgacagctattg<br>ggtctttgtgaaacgggtgtaactcgaggactacaaggacgacgacgacaagcccgg<br>gatccgcccctctccctcccccccccctaacgttactggccgaagccgcttggaataag<br>gccggtgtgcgtttgtctatatgttattttccaccatattgccgtcttttggcaatgtgagggcc<br>cggaaacctggccctgtcttcttgacgagcattcctaggggtctttcccctctcgccaaag<br>gaatgcaaggtctgttgaatgtcgtgaaggaagcagttcctctggaagcttcttgaagac<br>aaacaacgtctgtagcgaccctttgcaggcagcggaaccccccacctggcgacaggt<br>gcctctgcggccaaaagccacgtgtataagatacacctgcaaaggcggcacaacccc<br>agtgccacgttgtgagttggatagttgtggaaagagtcaaatggctctcctcaagcgtatt<br>caacaaggggctgaaggatgcccagaaggtaccccattgtatgggatctgatctgggg<br>cctcggtacacatgctttacatgtgtttagtcgaggttaaaaaaacgtctaggccccccga<br>accacggggacgtggttttcctttgaaaaacacgatgataatatggccacaaccatgcg<br>atcgtacccatacgatgttccagattacgctgaattcgtgagcaagggcgaggagctgtt<br>caccggggtggtgcccatcctggtcgagctggacggcgacgtaaacggccacaagttc<br>agcgtgtccggcgagggcgagggcgatgccacctacggcaagctgaccctgaagttc<br>atctgcaccaccggcaagctgcccgtgccctggcccaccctcgtgaccaccctgaccta<br>cggcgtgcagtgcttcagccgctaccccgaccacatgaagcagcacgacttcttcaagt<br>ccgccatgcccgaaggctacgtccaggagcgcaccatcttcttcaaggacgacggca<br>actacaagacccgcgccgaggtgaagttcgagggcgacaccctggtgaaccgcatcg<br>agctgaagggcatcgacttcaaggaggacggcaacatcctggggcacaagctggagt<br>acaactacaacagccacaacgtctatatcatggccgacaagcagaagaacggcatc<br>aaggtgaacttcaagatccgccacaacatcgaggacggcagcgtgcagctcgccgac<br>cactaccagcagaacacccccatcggcgacggccccgtgctgctgcccgacaacca<br>ctacctgagcacccagtccgccctgagcaaagaccccaacgagaagcgcgatcaca<br>tggtcctgctggagttcgtgaccgccgccgggatcactctcggcatggacgagctgtaca<br>ag |

| | | |
|---|---|---|
| pAG1385 | cMyc-FRB-nirFAST$_{1-114}$-IRES-HA-mTurquoise2 | atggaacaaaagcttatttctgaagaggacttggaattcgagatgtggcatgaaggcctg<br>gaagaggcatctcgtttgtactttggggaaaggaacgtgaaaggcatgtttgaggtgctg<br>gagcccttgcatgctatgatggaacggggcccccagactctgaaggaaacatcctttaa<br>tcaggcctatggtcgagatttaatggaggcccaagagtggtgcaggaagtacatgaaat<br>cagggaatgtcaaggacctcacccaagcctgggacctctattatcatgtgttccgacga<br>atctcaaagcaggtc—<br>tccggaggaggcggcagcggcggaggggatccatggagcatgttgcctttggcagt<br>gaggacatcgagaacactttggccaaaatggacgacggacaactggatgggttggcc<br>tttggcgcagttcagctcgatggtaacgggaatatcctgaagtacagtgctgctcaggga<br>ggcatcacaggcaaagatcccaaacaggtgattgggaagaacttattcaaggatgttg<br>caccaggatcggtttcctccgggttttacggcatattcaaggaaggcgtagcgtcaggga<br>atctgaacaccatgttcgaatggatgataccgacaagcaggggggccaactaaggtcaa<br>ggtgcatatgaagaaagccctttcctaacctcgaggactacaaggacgacgacgaca<br>agcccgggatccgcccctctccctcccccccccctaacgttactggccgaagccgcttg<br>gaataaggccggtgtgcgtttgtctatatgttattttccaccatattgccgtcttttggcaatgt<br>gagggcccggaaacctggccctgtcttcttgacgagcattcctaggggtctttcccctctc<br>gccaaaggaatgcaaggtctgttgaatgtcgtgaaggaagcagttcctctggaagcttct<br>tgaagacaaacaacgtctgtagcgaccctttgcaggcagcggaaccccccacctggc<br>gacaggtgcctctgcggccaaaagccacgtgtataagatacacctgcaaaggcggca<br>caaccccagtgccacgttgtgagttggatagttgtggaaagagtcaaatggctctcctca<br>agcgtattcaacaaggggctgaaggatgcccagaaggtaccccattgtatgggatctg<br>atctggggcctcggtacacatgctttacatgtgtttagtcgaggttaaaaaaacgtctagg<br>ccccccgaaccacggggacgtggttttcctttgaaaaacacgatgataatatggccaca<br>accatgcgatcgtacccatacgatgttccagattacgct<br>gaattcatggtgagcaagggcgaggagctgttcaccggggtggtgcccatcctggtcga<br>gctggacggcgacgtaaacggccacaagttcagcgtgtccggcgagggcgagggcg<br>atgccacctacggcaagctgaccctgaagttcatctgcaccaccggcaagctgcccgt<br>gccctggcccaccctcgtgaccaccctgtcctggggcgtgcagtgcttcgcccgctaccc<br>cgaccacatgaagcagcacgacttcttcaagtccgccatgcccgaaggctacgtccag<br>gagcgcaccatcttcttcaaggacgacggcaactacaagacccgcgccgaggtgaag<br>ttcgagggcgacaccctggtgaaccgcatcgagctgaagggcatcgacttcaaggag<br>gacggcaacatcctggggcacaagctggagtacaactacttcagcgacaacgtctata<br>tcaccgccgacaagcagaagaacggcatcaaggccaacttcaagatccgccacaac<br>atcgaggacggcggcgtgcagctcgccgaccactaccagcagaacacccccatcgg<br>cgacggccccgtgctgctgcccgacaaccactacctgagcacccagtccaagctgag<br>caaagaccccaacgagaagcgcgatcacatggtcctgctggagttcgtgaccgccgc<br>cgggatcactctcggcatggacgagctgtacaag |

| | | |
|---|---|---|
| pAG1501 | TOM20(1-34)-ECFP-nirFAST$_{1-114}$-linker | atggtgggtcggaacagcgccatcgccgcgggcgtgtgcggtgccctcttcatagggta<br>ctgcatctactttgaccgcaaaagacgaagtgaccccaacttcggatccagtgctggtg<br>gtagtgctggtggtatggtgagcaagggcgaggagctgttcaccggggtggtgcccatc<br>ctggtcgagctggacggcgacgtaaacggccacaagttcagcgtgtccggcgagggc<br>gagggcgatgccacctacggcaagctgaccctgaagttcatctgcaccaccggcaag<br>ctgcccgtgccctggcccaccctcgtgaccaccctgacctggggcgtgcagtgcttcag<br>ccgctaccccgaccacatgaagcagcacgacttcttcaagtccgccatgcccgaaggc<br>tacgtccaggagcgcaccatcttcttcaaggacgacggcaactacaagacccgcgcc<br>gaggtgaagttcgagggcgacaccctggtgaaccgcatcgagctgaagggcatcgac<br>ttcaaggaggacggcaacatcctggggcacaagctggagtacaactacatcagccac<br>aacgtctatatcaccgccgacaagcagaagaacggcatcaaggccaacttcaagatc<br>cgccacaacatcgaggacggcagcgtgcagctcgccgaccactaccagcagaaca<br>cccccatcggcgacggccccgtgctgctgcccgacaaccactacctgagcacccagtc<br>cgccctgagcaaagaccccaacgagaagcgcgatcacatggtcctgctggagttcgtg<br>accgccgccgggatcactctcggcatggacgagctgtacaagggagcaagtggaatg<br>gagcatgttgcctttggcagtgaggacatcgagaacactttggccaaaatggacgacg<br>gacaactggatgggttggcctttggcgcagttcagctcgatggtaacgggaatatcctga<br>agtacagtgctgctcagggaggcatcacaggcaaagatcccaaacaggtgattggga<br>agaacttattcaaggatgttgcaccaggatcggtttcctccgggttttacggcatattcaag<br>gaaggcgtagcgtcagggaatctgaacaccatgttcgaatggatgataccgacaagc<br>agggggccaactaaggtcaaggtgcatatgaagaaagccctttccagtgctggtggta<br>gtgctggtggtagtgctggtggtagtgctggtggtactggtggtcctcgagct |

| | | |
|---|---|---|
| | KIF17MD-flag-nirFAST$_{1-114}$-linker | atggcctccgaggcggtgaaggttgtcgtgcgctgccgtcccatgaaccagcgggagc<br>gagagctgcgctgccagcccgtggtgactgtggactgcgcgcgcgcccagtgctgcat<br>ccagaacccgggcgccgccgacgagccgcccaagcagttcaccttcgacggcgcct<br>accacgtggaccacgtcaccgagcagatctacaacgagatcgcctatccgctggtgga<br>gggcgtcactgagggctacaatggcaccatctttgcctacggccagacaggcagcgg<br>gaagtccttcaccatgcagggcctgccggatccgccctcccagagaggcatcatcccc<br>agggccttcgagcacgtgttcgagagcgtccagtgtgcagagaacactaagttcctggt<br>ccgggcctcctacctggagatctacaatgaagatgtccgggacctccttggggctgaca<br>ccaagcagaagctggagctgaaggagcacccagagaagggcgtgtacgtgaaggg<br>gctgtccatgcacacggtgcacagcgtggcccagtgtgagcacatcatggagactggc<br>tggaagaaccgttcggtcggctacacgctgatgaacaaggattcctcacgctcgcactc<br>catcttcaccatcagcatcgagatgtctgccgtggatgagcggggcaaggaccacctcc<br>gggcgggcaagctgaacctggtggacctggcgggcagcgagcggcagtccaagac<br>cggggccacgggcgagcggctcaaggaggccaccaagatcaacctgtcgctctcgg<br>cactgggcaatgtcatctcggcgctggtggacgggcgctgtaagcacgtcccctaccgt<br>gactcgaagctgacgcggctgctgcaggactcactgggcggcaacaccaagacgctc<br>atggtggcctgcctgtcgcctgcggacaacaactacgatgagacactcagcacgctgc<br>gctacgccaaccgggccaagaacatcaggaacaagccgcgcatcaatgaggaccc<br>caaggatgcgctgcttcgcgagtaccaggaggagatcaagaagctcaaggccatcct<br>gacacagcagatgagccccagcagcctgtcagccctgctgtccaggcaggtgccccc<br>agaccctgtgcaggtggaggagaagctgttgccccaacctgtgatccagcatgacgtg<br>gaggccgagaagcagctgatccgggaggagtatgaagagcgcctggcccggctgaa<br>agccgactataaggccgagcaggagtctcgggccaggctggaggaagacatcactg<br>ccatgcgcaactcatatgacgtcaggctgtccacgctggaggagaacctgcggaagg<br>agacagaggctgtcctgcaggtgggagtcctctacaaggctgaggtcatgtccagggct<br>gagtttgccagcagcgctgagtacccgcctgcttttcagtatgagacagtggtgaaaccc<br>aaggtcttctccacgactgacactctgcccagtgacgatgtctccaagactcaggtttcct<br>ccaggtttgcggagctgcccaaggtggaaccctccaaatctgagatttctctgggctcca<br>gtgagtcatcctcgctcgaagaaacccttgcggccgctgattacaaggatgacgatgac<br>aagttaattaatggagcaagtggaatggagcatgttgcctttggcagtgaggacatcga<br>gaacactttggccaaaatggacgacggacaactggatgggttggcctttggcgcagttc<br>agctcgatggtaacgggaatatcctgaagtacagtgctgctcagggaggcatcacagg<br>caaagatcccaaacaggtgattgggaagaacttattcaaggatgttgcaccaggatcg<br>gtttcctccgggttttacggcatattcaaggaaggcgtagcgtcagggaatctgaacacc<br>atgttcgaatggatgataccgacaagcaggggggccaactaaggtcaaggtgcatatga<br>agaaagccctttccagtgctggtggtagtgctggtggtagtgctggtggtactggtggtcct<br>cgagct |

| | | |
|---|---|---|
| | pFAST-zGem(1-100)-P2A-nirFAST-zCdt1(1-190) | atggagcatgttgcctttggcagtgaggacatcgagaacactctcgctaatatggacgac<br>gaacaactggataggttggcctttggcgtgattcagctcgatggtgacgggaatatcctg<br>ctgtataatgctgctgaaggagacatcacaggcagagatcccaaacaggtgattggga<br>agaacttcttcaaggatgttgcacctggaacagatactcccgagttttacggcaaattcaa<br>ggaaggcgcagcgtcaggaaatctgaacaccatgttcgaatggacaataccgacaag<br>caggggacctaccaaggtcaaggtgcacctgaagaaagccctttccggtgacagatat<br>tgggtctttgtgaaacgggtg-<br>cccgggtctcctggaccccgcggatacccttctcactggaggccactggag-<br>atgagttccatcagaaggccaaagaacgcagagaacccttctgagaacatcaagaaa<br>ttcctggtggctcctacatctggagcaggaatggaagaaggacacttcaggtgctcca<br>gccctctgctgtgaacaaaaacctgggtcgcatcattgagaatggcaaagctgtggcta<br>aaaggaagatgtggagcgccgagcaagtgaagggttcaaagagagtcaaagcaga<br>ggtggctgtcaaatccacaaatgcagagaatgagaaccagccagagggagtcacac<br>aggaggcttatgag-<br>agcgctgtgagcggctccggagccacaaacttctctctgctcaagcaggctggagatgt<br>ggaagaaaaccctggacctaggagcggaatggccacaactccgtctcac-<br>atggagcatgttgcctttggcagtgaggacatcgagaacactctcgcgaaaatggacg<br>acggacaactggatgggttggcctttggcgcagttcagctcgatggtaacgggaatatcc<br>tgaagtacagtgctgctcagggaggcatcacaggcaaagatcccaaacaggtgattg<br>ggaagaacttattcaaggatgttgcaccaggatcggtttcctccgggttttacggcatattc<br>aaggaaggcgtagcgtcagggaatctgaacaccatgttcgaatggatgataccgaca<br>agcagggggccaactaaggtcaaggtgcatatgaagaaagccctttccggtgacagct<br>attgggtctttgtgaaacgggtg-<br>cccgggtctcctggagaattctcaatgctgccttctctgggaggctatccatcacattggag<br>gccactggaa-<br>atggcccaggctcgcgttactgattattttgcacagagtaaaaaagctggtgtttctcgatct<br>ttacggtctaagggacaaaaggtgtccggggatgtagtagaatcagcggtgattaacaa<br>acctagaagttcgagcagagcgtccggttcttcacgcaaagcgacgcacagtgagatc<br>accacagcagagccgcagaaacaaactcaactggagtttctcaaagtcatcgacgag<br>gctttatcgacggagactgctgataccgtcgcggattcgcgtgatgtgaagattgagggc<br>ttaacggcgagtcccagaacccctaagagaagttcaccggagttcgatgtgtgttcggtg<br>ttgtttccgtccacagccgagctgcatagcagcgccaagaagcggcagagactcaacg<br>cgggccacaactgtagaagctcaccggaggagagagctggacagaaaactgcgag<br>gaaaaaactctatctactggcaagcgacgacaaagtaaagacaaccgagcctctggc<br>ttctagcagcccacaggcacctcaacaaactgcccgcaaagag |
| pAG721 | lyn11-mCherry-cMyc | atgggctgcatcaagtccaagggcaaggactccgccggcggcggctccatggtgagc<br>aagggcgaggaggataacatggccatcatcaaggagttcatgcgcttcaaggtgcac<br>atggagggctccgtgaacggccacgagttcgagatcgagggcgagggcgagggccg<br>cccctacgagggcacccagaccgccaagctgaaggtgaccaagggtggccccctgc<br>ccttcgcctgggacatcctgtcccctcagttcatgtacggctccaaggcctacgtgaagc<br>accccgccgacatccccgactacttgaagctgtccttccccgagggcttcaagtgggag<br>cgcgtgatgaacttcgaggacggcggcgtggtgaccgtgacccaggactcctccctgc<br>aggacggcgagttcatctacaaggtgaagctgcgcggcaccaacttcccctccgacgg<br>ccccgtaatgcagaagaagaccatgggctgggaggcctcctccgagcggatgtaccc<br>cgaggacggcgccctgaagggcgagatcaagcagaggctgaagctgaaggacggc<br>ggccactacgacgctgaggtcaagaccacctacaaggccaagaagcccgtgcagct<br>gcccggcgcctacaacgtcaacatcaagttggacatcacctcccacaacgaggacta<br>caccatcgtggaacagtacgaacgcgccgagggccgccactccaccggcggcatgg<br>acgagctgtacaagggatccgaacaaaagcttatttctgaagaggacttg |

### Supplementary Methods

#### Construction of yeast display libraries by random mutagenesis

The two combinatorial yeast libraries (called library A and B in this study) of nirFAST0.1 and nirFAST1.1 variants were constructed by error-prone PCR using Genemorph II kit (Agilent). They were constructed by randomization of the sequences of nirFAST0.1 and nirFAST1.1 by error-prone PCR using primers ag216 and ag217. PCR product was digested using restriction enzymes NheI and BamHI (New England Biolabs) and ligated into the pCTCON2 vector using NheI and BamHI restriction sites. For the two libraries, small-scale transformations were performed by electroporation in DH10B *E. coli* to optimize the number of transformants and control the mutation rate. Large-scale transformation performed by electroporation in DH10B *E. coli* cells led to 3.5 $\times 10^7$ transformants for library A and 2 $\times 10^7$ transformants for Library B. DNA was then minipreped, and retransformed in EBY100 yeast strain using large-scale high-efficiency transformation protocol[13,14], leading to $10^6$ transformants for Library A and 3.9 $\times 10^6$ transformants for library B.

#### Selection

Libraries (typically 1 $\times 10^{10}$ cells) were grown overnight (30°C, 280 rpm) in 1 L of SD (20 g/L dextrose, 6.7 g/L yeast nitrogen base, 1.92 g/L yeast synthetic dropout without tryptophan, 7.44 g/L $NaH_2PO_4$ and 10.2 g/L $Na_2HPO_4$-7$H_2O$, 1% penicillin-streptomycin 34 10,000 U/mL). 1 $\times 10^{10}$ cells yeast cells were then collected and grown for 36 h (23°C, 280 rpm) in 1L SG (20 g/L galactose, 2 g/L dextrose, 6.7 g/L yeast nitrogen base, 1.92 g/L yeast synthetic dropout without tryptophane, 7.44 g/L $NaH_2PO_4$, 10.2 g/L $Na_2HPO_4$-7$H_2O$, 1% penicillin-streptomycin 10,000 U/mL). 6 $\times 10^8$ induced cells were then pelleted by centrifugation (25°C, 3 min, 2,500 $\times$ g), washed with 10 mL DPBS-BSA (137 mM NaCl, 2.7 mM KCl, 4.3 mM $Na_2HPO_4$, 1.4 mM $KH_2PO_4$, 1 g/L bovine serum albumin, pH 7.4), and incubated for 30 min at room temperature in 200 µL of 1/250 primary antibody chicken antic-CMyc IgY (Life Technologies) solution in DPBS-BSA. Cells were then washed with 10 mL DPBS-BSA, and incubated in 200 µL of 1/100 secondary antibody Alexa Fluor® 488–goat anti-rabbit IgG (Life Technologies) solution in DPBS-BSA for 30 min on ice. After washing with DPBS-BSA, cells were incubated in 10 mL DPBS-BSA supplemented with indicated concentration of HPAR-3,5DOM, and sorted on a MoFlo™ Astrios Cell sorter (Beckman Coulter) equipped with a 488 nm and a 640 nm laser (see **Supplementary Figure 16** for the gating strategy). The sorted cells, displaying high green and NIR fluorescence were collected in SD, grown overnight (30°C, 240 rpm) and spread on SD plates (SD supplemented with 182 g/L sorbitol, 15 g/L agar). Plates were incubated for 60 h at 30°C. The cell lawn was collected in SD supplemented with 30% glycerol, aliquoted and frozen or directly used in the next round.

**Cloning of the different variants**

The plasmids used in this study have been generated using isothermal Gibson assembly or restriction enzymes cloning.

The plasmids pAG985 and pAG1150 for the yeast expression of Aga2p-HA-nirFAST0.1-cMyc and Aga2p-nirFAST1.1-cMyc were obtained by introducing the sequence of nirFAST0.1 and nirFAST1.1 in the pCTCON2 vector using NheI and BamHI restriction sites.

The plasmid pAG456 for bacterial expression of 6×His–nirFAST0.1 was obtained by introduction of the mutation E46Q in the plasmid pAG419 for bacterial expression of 6×His–frFAST$^{I107V}$[1]. The plasmid pAG922 for bacterial expression of 6×His–frFAST$^{E46Q}$ was obtained by introduction of the point mutation E46Q in the sequence of frFAST in the plasmid pAG455 for bacterial expression of 6×His–frFAST[1]. The plasmids pAG1103, pAG1104, pAG1105 (**nirFAST1.0**), pAG1132 and pAG1130 for bacterial expression of clones A.R5-5, A.R5-22, A.R5-48, A.R5-24 and A.R5-18 respectively were obtained by restriction enzyme cloning (NheI and XhoI sites) by insertion of their coding sequences in the pET28 vector. The plasmid pAG1135 for bacterial expression of 6×His–nirFAST1.0$^{N43S}$ was obtained by introduction of the point mutation N43S in the plasmid pAG1105 for bacterial expression of 6×His–nirFAST1.0. The plasmid pAG1136 for bacterial expression of nirFAST1.0$^{K78I}$ was obtained by introduction of the point mutation K78I in the sequence of the plasmid pAG1105 for bacterial expression of 6×His–nirFAST1.0. The plasmid pAG1137 (**nirFAST1.1**) for bacterial expression of 6×His–nirFAST1.1 (a.k.a. nirFAST1.0$^{N43S,K78I}$) was obtained by introduction of the mutation K78I in the plasmid pAG1135 for bacterial expression of 6×His–nirFAST1.0$^{N43S}$. The plasmid pAG1139 for bacterial expression of A.R5-18.1 was obtained by introduction of the mutation T70S in the plasmid pAG1130 for bacterial expression of 6×His–A.R5-18. The plasmid pAG1140 for bacterial expression of A.R5-22.1 was obtained by introduction of the mutation T70S in the plasmid pAG1104 for bacterial expression of 6×His–A.R5-22. The plasmid pAG1208 for bacterial expression of 6×His–B.R5-24 (**nirFAST2.0**) was obtained by replacing the sequence of nirFAST1.1 by the sequence of clone B.R5-24 in the plasmid pAG1137 for bacterial expression of 6×His–nirFAST1.1. The plasmids pAG1225, pAG1226, pAG1227 and pAG1228 for bacterial expression of clones B.R5-06, B.R5-67, B.R5-60 and B.R5-37 were obtained by introducing their coding sequence in the plasmid pAG1137 for bacterial expression of 6×His–nirFAST1.1. The plasmids pAG1236, pAG1249, pAG1250, pAG1251, pAG1252, pAG1253, pAG1254 and pAG1255 for bacterial expression of the single variants of nirFAST2.0 were obtained by introducing respectively the mutations V107I, G21R, G25R, K41R, G47R, R52K (mutant corresponding to **nirFAST**), M95T and S117R in the sequence of pAG1208 for bacterial expression of expression of 6×His–nirFAST2.0. Plasmids 1304, 1305, 1327, 1328, 1330 for mammalian expression of nirFAST2.0, nirFAST2.0$^{M95T}$, nirFAST2.0$^{K41R}$, nirFAST2.0$^{G47R}$ and nirFAST2.0$^{V107I}$ were obtained by replacing the sequence of FAST by the sequences of each mutant in the plasmid pAG453 allowing the mammalian expression of FAST-P2A-EGFP (not published).

### Protein Sequences

#### frFAST

MEHVAFGSEDIENTLAKMDDGQLDGLAFGAIQLDGDGNILQYNAAEGDITGRDPKQVIGKNL
FKDVAPGTVSSGFYGKFKEGVASGNLNTMFEWMIPTSRGPTKVKIHMKKALSGDSYWVFV
KRV

#### nirFAST

MEHVAFGSEDIENTLAKMDDGQLDGLAFGA**V**QLDG**N**GNIL**K**Y**S**AA**Q**G**G**ITG**K**DPKQVIGKN
LFKDVAPG**S**VSSGFYG**I**FKEGVASGNLNTMFEWMIPTSRGPTKVK**V**HMKKALSGDSYWVF
VKRV
